# White matter micro- and macrostructural properties in midlife individuals at risk for Alzheimer’s disease: Associations with sex and menopausal status

**DOI:** 10.1101/2025.06.09.658686

**Authors:** Adam C. Raikes, Jonathan P. Dyke, Matilde Nerattini, Camila Boneu, Trisha Ajila, Francesca Fauci, Michael Battista, Silky Pahlajani, Schantel Williams, Roberta Diaz Brinton, Lisa Mosconi

## Abstract

Women are at greater lifetime risk for Alzheimer’s disease (AD), potentially due to midlife fractional anisotropy (FA) and lower mean diffusivity in fornix and corpus callosum, indicating more densely organized white matter. Perimenopausal women were the exception, with white matter profiles closely resembling those of men. Perimenopausal women exhibited minimal or absent fiber cross-section and FDC sex differences and a reversal of the fornix FA advantage observed in pre- and postmenopausal women. These cross-sectional results are consistent with sex differences in white matter organization. Importantly, the perimenopause emerges as a critical window of neural reorganization in the female midlife aging brain characterized by temporary convergence toward male-like white matter organization. Longitudinal analyses are key to identifying women who do or do not revert to a premenopausal profile, which may inform AD risk.

## Introduction

Women have a greater prevalence of late-onset Alzheimer’s disease, independent of age and survival rates, with postmenopausal women representing more than 60% of AD cases^1^. AD has a long prodromal period lasting 10-20 years prior to clinical diagnosis. This prodromal phase aligns with the menopausal timeline, suggesting that menopause may play a role in elevated risk. This is supported by cross-sectional studies demonstrating increased amyloid-beta and tau pathology, decreased gray matter volume, cerebral glucose dysregulation, and altered mitochondrial function in AD-vulnerable brain regions in peri- and postmenopausal women compared to either premenopausal women or age-matched men^2–7^. Less well-described than these AD-relevant markers is the role of sex and menopause on white matter.

Fractional anisotropy (FA) is the most frequently reported metric of white matter microstructure, with higher values generally associated with greater white matter health. Across the literature, there is substantial variability regarding sex differences in FA, with studies reporting men having higher FA than women^8–11^, women having higher FA than men^12,13^, or limited to no sex differences at all ^14,15^. Regions most commonly associated with sex differences include the fornix, corpus callosum, cingulum, corona radiata, and thalamic radiata. Women typically show greater FA in the fornix^2,16–19^ while men often show greater FA in the cingulum and thalamic radiata^2,12,18,20,21^. The corpus callosum exhibits mixed profiles across studies^12,15–17,22,23^.

One understudied area is differences between men and women in the macrostructural organization (fiber density, cross sectional area) of white matter. One method for probing such macrostructural features using diffusion MRI is a “fixel-based” approach^24^. This approach characterizes the fiber orientations present within a voxel, providing measures of both fiber density and fiber-cross section^24^. This provides complementary information to traditional diffusion MRI tensor metrics, capturing measures sensitive to fiber loss, bundle atrophy, and a combined metric that indexes the simultaneous contributions of density and cross-section (FDC)^24,25^. Recent work using this approach has demonstrated reduced fiber density, fiber cross-section, and FDC in both mild cognitive impairment and mild amnestic AD compared to cognitively unimpaired individuals, primarily in limbic and callosal fiber bundles^26–28^ indicating both fiber loss and bundle atrophy to be disease-related features. However, sex-specific analyses are lacking. One study has provided some evidence indicating modest sex differences in both fiber density and cross-section in several white matter tracts, but sex differences were not a primary focus of the analyses and comprehensive reporting is lacking^29^. The absence of more comprehensive evaluations of these macrostructural differences limits our understanding of how sex differences may contribute to disease risk and onset.

Herein we conducted a cross-sectional analysis of sex differences in both macro- and microstructural white matter properties in cognitively normal women and men in midlife at risk for AD (e.g. AD family history and/or APOE-e4 genotype), and tested for associations with menopausal status (pre-, peri-, and post-menopausal). Given the lack of consensus on the presence, absence, and spatial distribution of sex differences in white matter, we employed a Bayesian hierarchical analysis framework in order to more completely describe the evidence for and against, as well as the directionality of, sex differences by incorporating all tracts into singular models in order to leverage partial pooling, mitigating statistical concerns of multiplicity^30,31^. Using this strategy, we aimed to provide a robust framework for identifying sex and menopause status differences as well as a platform for future longitudinal analyses.

## Results

### Demographics

A total of 137 individuals with complete datasets were included in these analyses. These included 34 pre-, 39 peri-, and 27 post-menopausal women and a total of 37 men. All participants were in good general health. Demographic data are presented in Table 1.

**Table 1.** Demographic characteristics.

| Characteristic | Whole dataset |  | Menopause subgroups |  |  |
| --- | --- | --- | --- | --- | --- |
|  | Male,<br>N = 37 | Female,<br>N = 100 | Pre,<br>N = 34 | Peri,<br>N = 39 | Post,<br>N = 27 |
| Age<sup>1</sup> | 51.35 (7.68) | 49.49 (6.20) | 44.06 (3.13) | 49.00 (3.74) | 57.04 (3.95) |
| Years of education<sup>1</sup> | 17.51 (2.02) | 17.10 (1.54) | 17.00 (1.23) | 16.97 (1.77) | 17.41 (1.55) |
| Total cholesterol (mg/dL)<sup>1</sup> | 180.81 (39.18) | 199.87<br>(32.27) | 190.56<br>(36.54) | 204.97<br>(28.20) | 204.41<br>(30.42) |
| BMI<sup>1</sup> | 27.29 (4.60) | 25.03 (5.53) | 24.19 (4.90) | 26.36 (6.03) | 24.09 (5.28) |
| Total intracranial volume<br>(cm<sup>3</sup>)<sup>1</sup> | 1,624.45<br>(134.20) | 1,411.37<br>(95.57) | 1,417.00<br>(99.21) | 1,414.82<br>(98.36) | 1,399.30<br>(89.09) |
| APOE e4 carriers<sup>2</sup> | 21 (57%) | 48 (48%) | 19 (56%) | 16 (41%) | 13 (48%) |
| Family history of<br>Alzheimer's<sup>2</sup> | 25 (68%) | 82 (82%) | 25 (74%) | 36 (92%) | 21 (78%) |
| History of hypertension<sup>2</sup> | 8 (22%) | 6 (6.1%) | 0 (0%) | 3 (7.7%) | 3 (11%) |

<sup>1</sup> Mean (SD)<sup>2</sup> n (%)

### Sex differences

#### Full dataset

Figure 1A provides an overview of the number of tracts with identified sex differences at both levels of evidence. Fiber density and FDC were greater throughout the majority of the brain in women, with no tracts identified with men having greater values. Women had evidence of greater fiber cross-section in only 22.2% of the tracts, indicating that differences in FDC are driven by differences in fiber density, independent of fiber cross-section as well as differences in total brain volume.

**Figure 1:**
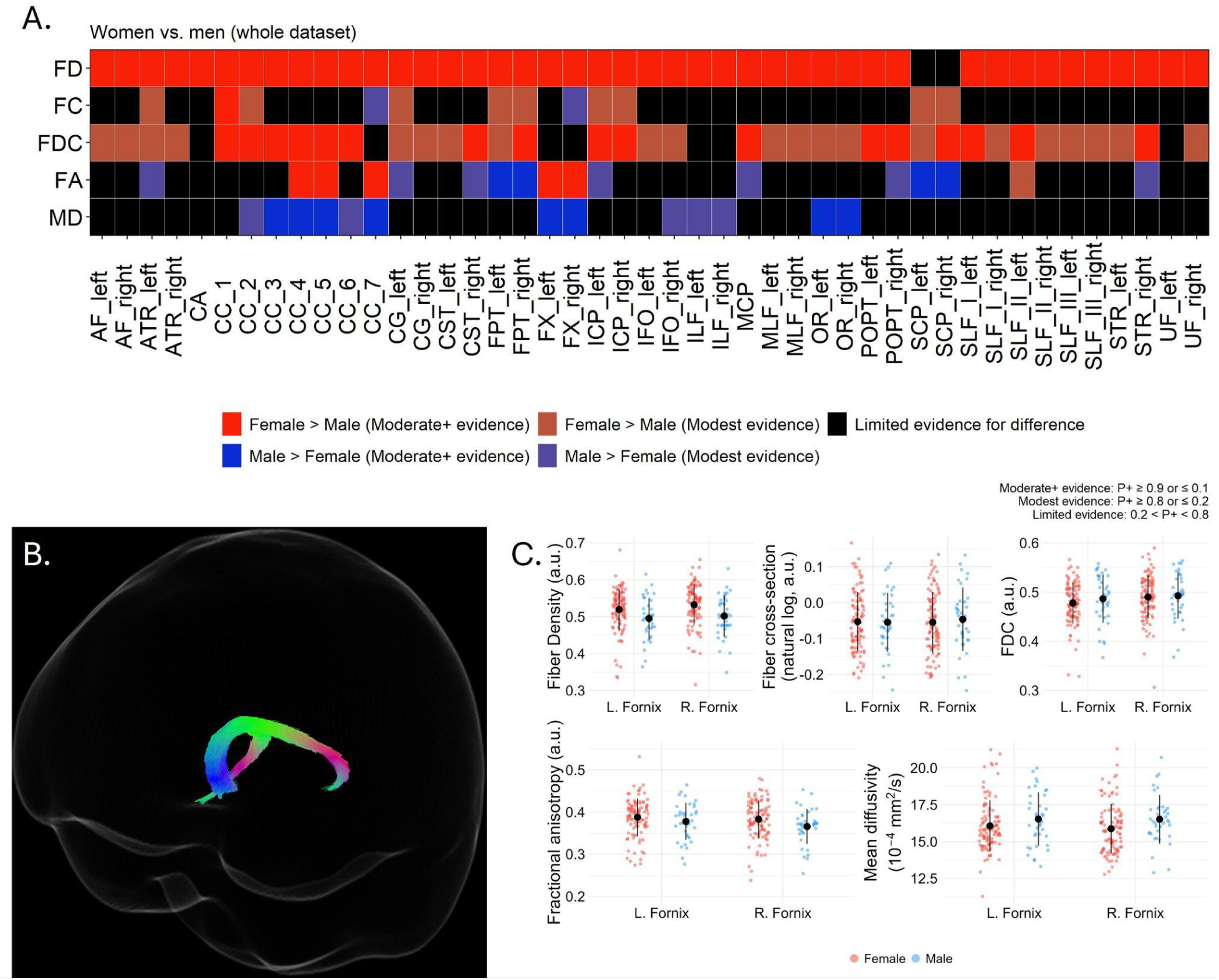
Sex differences in white matter metrics between men and women. **(A)** Heatmap showing tract-wise sex differences for FD, FC, FDC, FA, and MD across 45 bundles. Colors denote Female > Male (reds) or Male > Female (blues), with darker shades indicating stronger evidence (P+ > 0.9 or < 0.1) and lighter shades indicating modest evidence (P+ > 0.8 or < 0.2). Black indicates limited evidence of difference. **(B)** Segmentation of the bilateral fornix. **(C)** Boxplots of residualized mean FD, logFC, FDC, FA, and MD in the corpus callosum for females (red) and males (blue), adjusted for covariates.

Additionally, men had greater FA in a total of 9 tracts ( P+ < 0.1: 4 tracts; Figure 1A), which included the left anterior and right superior thalamic radiata, frontopontine tracts, and cerebellar peduncle tracts. Women had evidence of greater white matter structural coherence in 12 tracts, exhibiting either greater FA, lower MD, or both compared to men. These were distributed in the corpus callosum, bilateral fornix, bilateral superior and inferior longitudinal fasciculi, and optic radiata.Exemplar data in 1B shows the bilateral fornix and 1C presents data for each scalar type residualized for APOE carriership, family ADRD history, age, education, and total brain volume (for fiber cross-section and FDC only). As evidenced by the moderate-to-strong evidence in the hierarchical Bayesian models (Figure 1A), mean values for FD and FA were higher in the women and lower in the men, while men had greater average mean diffusivity (MD) values in both fornices.

#### Pre-menopause

Figure 2A provides an overview of the number of tracts with identified sex differences at both levels of evidence for the pre-menopausal women and age-matched men. See Supplemental Tables 6-10 for full details including tract-wise mean values, model identified differences, 95%

**Figure 2:**
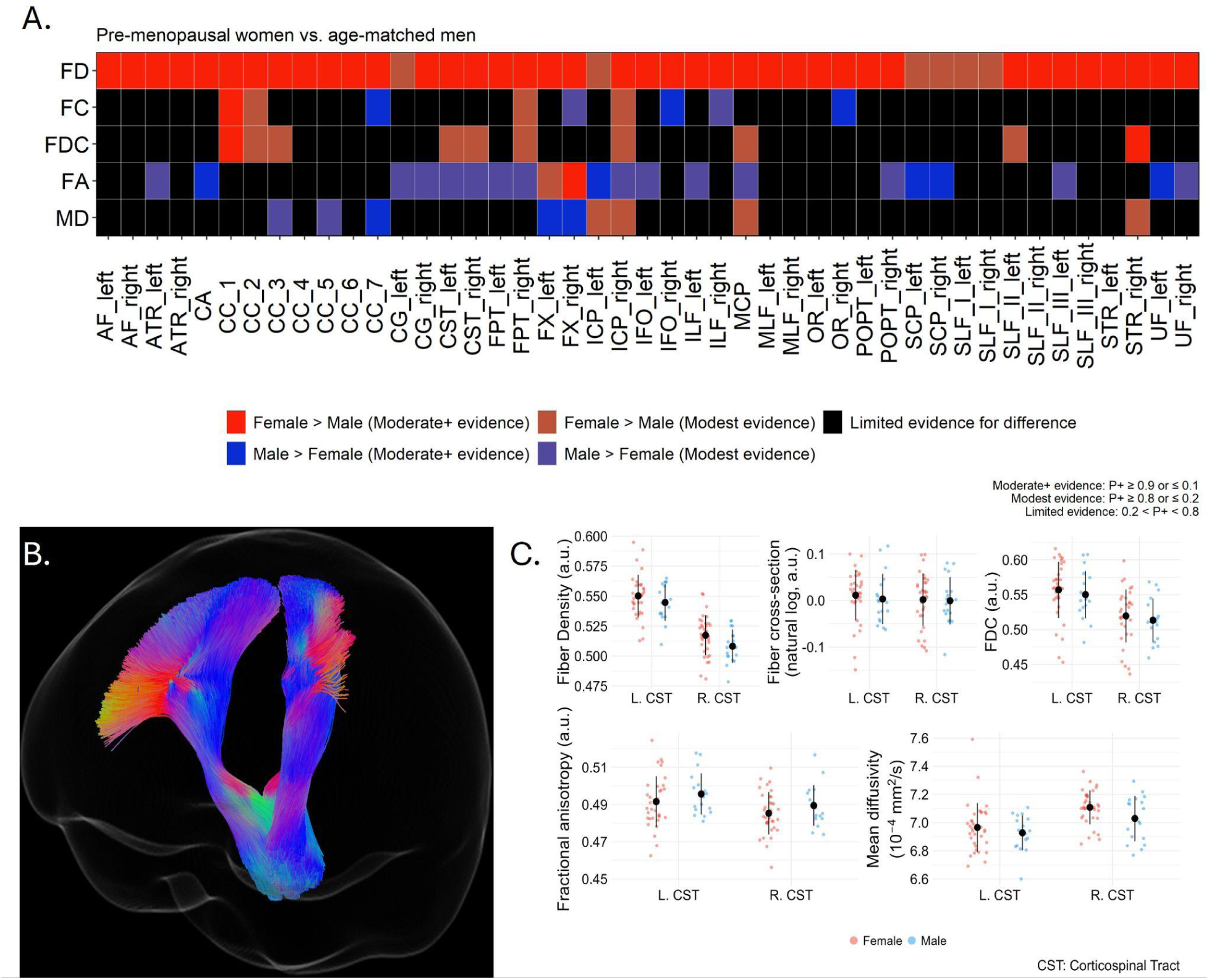
Sex differences in white matter metrics between premenopausal women and age-matched men. **(A)** Heatmap showing tract-wise sex differences for FD, FC, FDC, FA, and MD across 45 bundles. Colors denote Female > Male (reds) or Male > Female (blues), with darker shades indicating stronger evidence (P+ > 0.9 or < 0.1) and lighter shades indicating modest evidence (P+ > 0.8 or < 0.2). Black indicates limited evidence of difference. **(B)** Segmentation of the bilateral corticospinal tract. **(C)** Boxplots of residualized mean FD, logFC, FDC, FA, and MD in the corpus callosum for females (red) and males (blue), adjusted for covariates

HDI, and P+ values. As with the full dataset, women had greater fiber density in all tracts. By contrast, the number of tracts with greater fiber cross section or FDC in these women was less than that for the whole dataset (4 and 10 tracts compared to 8 and 38 respectively), suggesting that the effects observed at the whole dataset level are not driven by this younger age group. Age-controlled men exhibited evidence of greater fiber cross-section in five tracts, notably including the splenium of the corpus callosum and the right fornix where men also had greater cross-section at the whole dataset level.

As shown in Figure 2A, women exhibited greater FA in the bilateral fornix, as well as greater microstructural coherence in segments of the corpus callosum, where men had greater MD. By contrast, men had greater FA in a wider range of tracts including left anterior thalamic radiation, cingulum bundle, corticospinal tracts, inferior and superior cerebellar peduncles, and uncinate fasciculi.

Representative evidence of these differences in the bilateral corticospinal tract is presented in Figure 2B-C. After residualizing for other covariates, mean values for fiber density and FDC were greater in the women while men had greater FA.

#### Peri-menopause

The patterns observed in peri-menopause were substantially different from those observed at pre-menopause. The peri-menopausal group continued to have strong evidence of greater fiber density throughout the brain (45/45 tracts) compared to age-controlled men (Figure 3A). However, in contrast to the pre-menopausal group comparison, the peri-menopausal group did not exhibit greater fiber cross-section, widespread greater FDC, or greater FA in any tracts compared to men. For this comparison, men had greater FA in the bilateral fornix, which is a reversal from the whole dataset and the pre-menopausal analysis.

**Figure 3:**
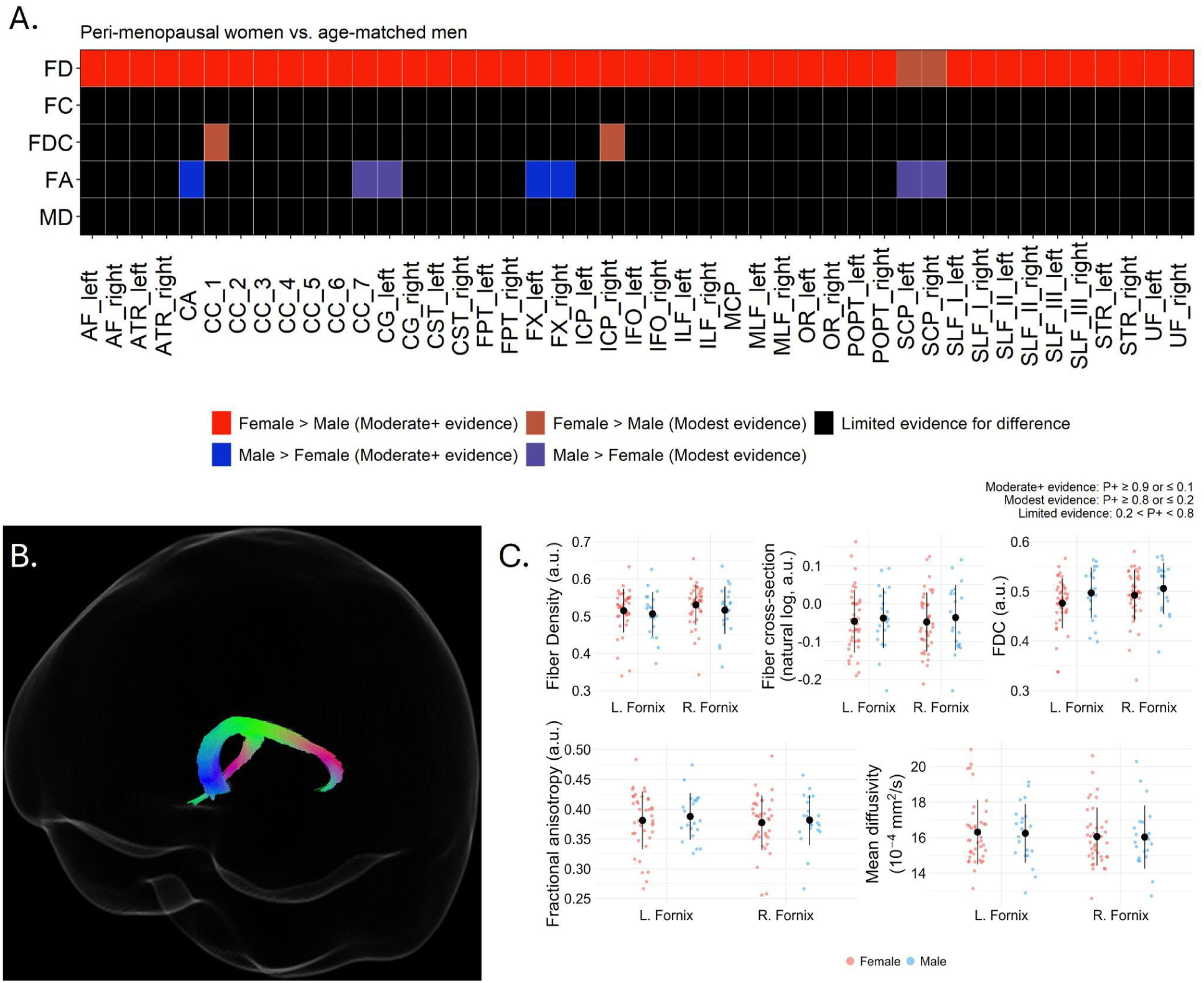
Sex differences in white matter metrics between perimenopausal women and age-matched men. **(A)** Heatmap showing tract-wise sex differences for FD, FC, FDC, FA, and MD across 45 bundles. Colors denote Female > Male (reds) or Male > Female (blues), with darker shades indicating stronger evidence (P+ > 0.9 or < 0.1) and lighter shades indicating modest evidence (P+ > 0.8 or < 0.2). Black indicates limited evidence of difference. **(B)** Segmentation of the bilateral fornix. **(C)** Boxplots of residualized mean FD, logFC, FDC, FA, and MD in the corpus callosum for females (red) and males (blue), adjusted for covariates.

Figure 3B-C provides representative findings from the bilateral fornix. Fiber density is greater in the women whale men had greater bilateral FA. Findings from the hierarchical models demonstrate that fiber cross-section, FDC, and mean diffusivity were comparable between men and women. See Supplemental Tables 11-15 for full details including tract-wise mean values, model identified differences, 95% HDI, and P+ values.

#### Post-menopause

At the age-range for the post-menopausal women, fiber bundle macrostructural differences were more extensive. The post-menopausal group had evidence of either greater fiber density, fiber cross-section, or FDC in every tract compared to age-controlled men (Figure 4A). It is, however, noteworthy for this group that whereas pre- and perimenopausal women had strong evidence greater fiber density in every nearly every tract, here there are 7 tracts without evidence of a difference and, while still the majority of the tracts, the number with moderate-to-strong evidence of a difference is down to 33/45.

**Figure 4:**
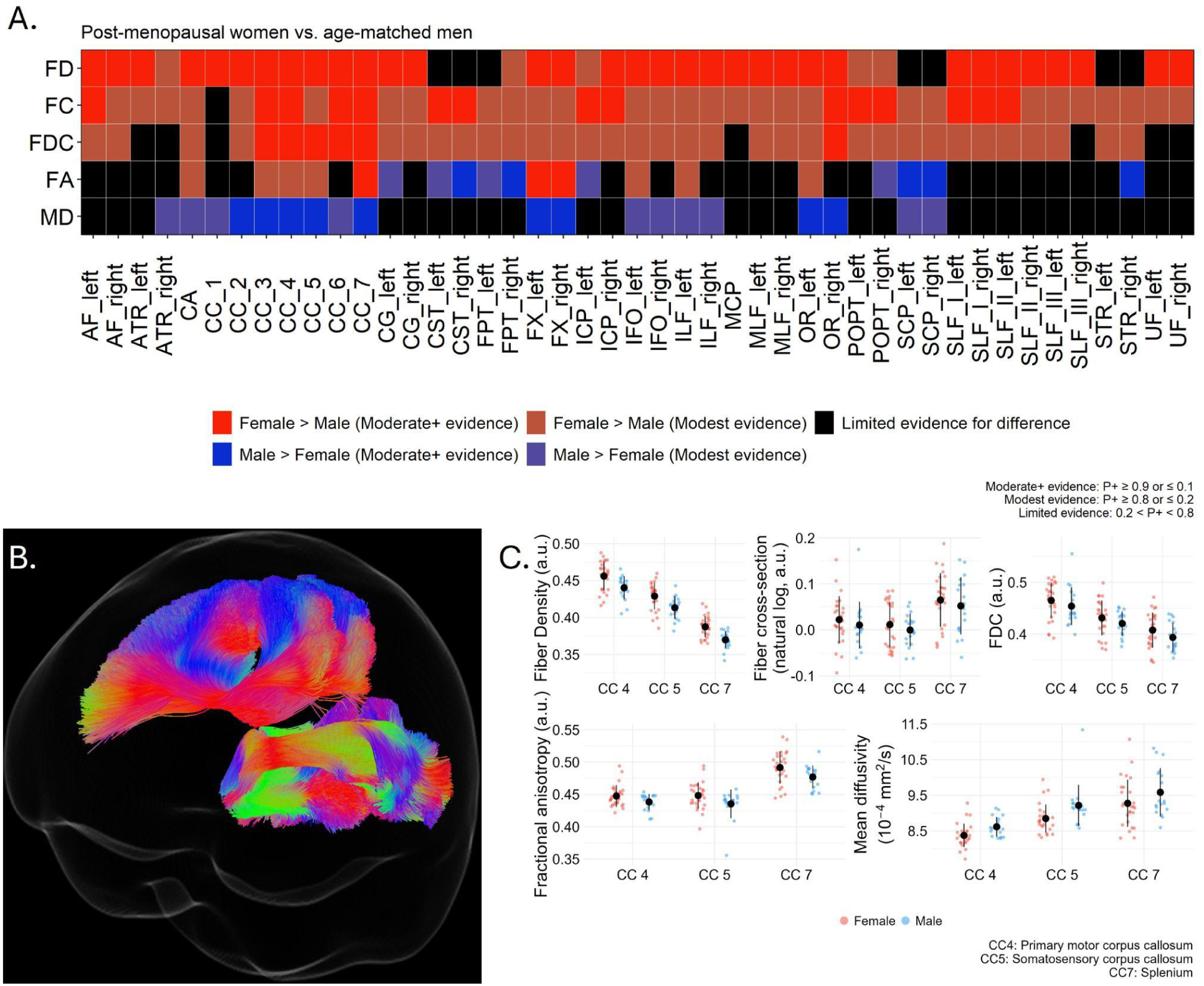
Sex differences in white matter metrics between postmenopausal women and age-matched men. **(A)** Heatmap showing tract-wise sex differences for FD, FC, FDC, FA, and MD across 45 bundles. Colors denote Female > Male (reds) or Male > Female (blues), with darker shades indicating stronger evidence (P+ > 0.9 or < 0.1) and lighter shades indicating modest evidence (P+ > 0.8 or < 0.2). Black indicates limited evidence of difference. **(B)** Segments 4, 5, and 7 of the corpus callosum. **(C)** Boxplots of residualized mean FD, logFC, FDC, FA, and MD in the corpus callosum for females (red) and males (blue), adjusted for covariates.

As with findings from the whole dataset as well as the pre-menopausal age group comparison, post-menopausal women had greater FA in the bilateral fornix and splenium of the corpus callosum compared to men, while men continued to exhibit greater FA in frontopontine tracts and cerebellar peduncle-related fiber bundles as well as the corticospinal tracts, left cingulum, and the right superior thalamic radiation.There was further evidence of greater microstructural coherence in the post-menopausal group, evidenced by greater mean diffusivity in the men throughout commissural tracts (corpus callosum, anterior commissure) as well as the optic radiata, and bilateral inferior orbital and longitudinal tracts.

Figure 4B-C highlights representative findings from three segments of the corpus callosum: The primary motor segment, somatosensory segment, and the splenium. In all three segments, women had greater fiber density, fiber cross-section, FDC, and FA while the men had greater mean diffusivity. See Supplemental Tables 15-20 for full details including tract-wise mean values, model identified differences, 95% HDI, and P+ values.

## Discussion

Present findings indicate that, among midlife individuals at risk for AD, women have greater fiber density throughout the brain relative to men. These findings were consistent in pre- and perimenopausal groups, while the number of the tracts exhibiting moderate-to-strong evidence of greater values was decreased by 27% in the postmenopausal group. Sex differences in white matter microstructural coherence were sparse and generally weaker and modulated by menopausal transition status, which are broadly consistent with frequently reported sex differences throughout the literature. Women on average, and specifically pre- and post-menopausal women, had greater FA and lower MD in the bilateral fornix as well as the corpus callosum compared to age-controlled men, whereas men had greater FA in the cerebellar peduncles, cingulum, and anterior commissure. Collectively, the findings indicate more globally tightly packed white matter along with greater white matter microstructural coherence in the fornix and corpus callosum in pre- and post-menopausal women. Results were independent of age, years of education, APOE genotype, and family history of AD, all of which may additionally and critically impact white matter health across midlife and later adulthood.

Fixel-based analyses are increasingly being used to characterize aging and disease-related changes in white matter micro- and macrostructure. Recent work in normal aging provides evidence that fiber density, cross-section, and FDC generally decrease with advancing age^29,32,33^. Findings from those studies suggest changes to both the microstructural environment due a loss of fibers (decreased FD) as well as atrophy (decreasing FC), culminating in less densely packed and thinner fiber bundles (FDC)^34^. These observations are further reflected in recent work in neurodegenerative diseases, where patients demonstrate reduced fiber density, fiber cross-section, and FDC in both mild cognitive impairment and mild AD compared to cognitively unimpaired individuals, primarily in limbic and callosal fiber bundles^26–28^. Those findings are independent of age, indicating both fiber loss and bundle atrophy to be disease-related features. However, the role of sex on fixed-based metrics in both aging and disease is relatively absent from the literature.

Choy et al (2020) used fixel-based metrics to examine aging in 293 individuals aged 21-86 and found a general pattern of age-related decreases in FD, FC, and FDC^29^. Across these three metrics, sex differences were only observed in fiber density for the anterior thalamic radiation and the hippocampal portion of the cingulum; however the authors did not indicate which sex had the greater mean value for either of these bundles. By contrast, we observed consistent evidence supporting the interpretation that women have greater fiber density throughout the brain and that there were no tracts in which men had greater density. Additionally we observed sparse evidence for fiber cross-section differences while FDC was greater in the women throughout the brain.

Interestingly, this greater fiber bundle packing in women did not manifest as extensively in the diffusion tensor metrics for these fiber bundles. Women across the menopausal transition represented here had evidence of greater fractional anisotropy in only 6 bundles. It is noteworthy that these findings agree with past work demonstrating greater FA in women in the fornix^12,13,16,17,19,35^ and the corpus callosum^12,17,23,36^. Men in this sample had greater mean diffusivity in these same tracts. This would be consistent with an interpretation that, for comparable fiber cross-sections, males’ less tightly packed bundles may allow for increased omnidirectional diffusivity. Men additionally had greater FA in tracts including the inferior, middle, and superior cerebral peduncles, superior and anterior thalamic radiata, as well as corticospinal and the frontopontine tracts, consistent with previous work^10,16,17,23,36^.

Collectively, these findings suggest that after accounting for both age and total brain volume, men and women in this sample had generally comparable bundle cross-section although women have more tightly packed white matter. This female advantage in fiber density (FD) may, additionally, be reflected by greater fractional anisotropy and lower mean diffusivity in women in tracts that are susceptible to both aging and neurodegenerative effects^9,12,27,28,33,37^. In light of the decreased FD observed in MCI and AD^27,28^ and the vulnerability of the female brain to AD, the findings in this cognitively normal, at-risk population may suggest either that white matter changes occur after the prodromal period or that these particular women are, on average, potentially resilient despite their risk factors. While either of these interpretations is plausible, both would require longitudinal data to substantiate them.

### Menopause as a modulator

Our sensitivity analyses provided additional insights into the nature of the observed differences. The observed fiber density patterns were largely consistent across all three sub-analyses by menopause status. By contrast, differences in fiber cross-section, FDC, FA, and mean diffusivity observed at both premenopause and postmenopause were largely absent in perimenopause, including a complete reversal of the directionality of the difference in FA in the fornix. These findings point to a potential U-shaped distribution with advancing endocrinological age, whereby perimenopausal women exhibit similar microstructural characteristics to comparably aged men. Additionally, the strength of evidence for fiber density differences between men and women was diminished at postmenopause compared to premenopause, suggesting permanent, and potentially destructive, remodeling of the white matter that occurs during perimenopause and those who are least able to recover those microstructural differences may be those at greatest risk for earlier neurodegenerative onset.

Several lines of work lend support for this interpretation. First, recent work has identified that circulating hormone levels, including 17B-estradiol and progesterone, are associated with dynamic changes in white matter diffusion and anisotropy properties across the menstrual cycle^38^, highlighting the temporal role of these hormones on white matter. This is further reflected by work suggesting that androgen and estrogen may confer masculinizing and feminizing effects, respectively, on white matter whereby the changing availability of and sensitivity to estrogen in the perimenopausal brain may briefly appear more like the male brain^23^. Second, estrogen receptors are highly expressed in oligodendrocytes and astrocytes in white matter^39,40^ and are critical orchestrators of estrogen-mediated neuroprotective signaling pathways^41^, oxidative processes, and metabolism^42,43^. Estrogen receptor density increases during menopausal transition in estrogen-regulated networks^44^, in agreement with findings of higher density in ovariectomized rats^45^. Finally, menopause-specific work demonstrates that the postmenopausal female brain undergoes multiple adaptations in both gray and white matter structure and organization, bioenergetics, and cognition^2–4^. Collectively, these results align with the notion that women’s brains undergo a dynamic remodeling during the menopause transition, which in this study extends to white matter fiber bundles and some connectivity measures.

Sex differences in cerebral metabolism have also been implicated in the sex-specific AD prevalence and differences in disease onset, pathophysiology and pathology progression as well as symptom presentation^2–6,46–50^. The declining regulatory effect of estrogen on glucose metabolism during menopause is thought to be a key driver of AD onset in women^40,51^. Animal studies demonstrate a catabolic effect on white matter in response to glucose hypometabolic states^52^, necessitating a remodeling of white matter pathways and a further bioenergetic adaptive response to preserve cognitive functioning^2^. Collectively, this body of work suggests that the bioenergetic adaptations required during menopause may, in part, be fulfilled through myelin catabolism until a more complete adaptive response can be mounted. The speed of this response likely dictates the extent to which white matter is remodeled and degraded, and as such may be a critical tipping point between neurodegenerative disease and resilience.

The current analysis excluded active menopausal hormone therapy (MHT) users and women with hysterectomy or oophorectomy, as these variables could confound detection of biomarker associations with menopause status. A natural extension of this work will be to explore interventions that could stabilize estrogen levels to support neuroendocrine aging in midlife women. Current guidelines recommend considering estrogen therapy, along with progestogen if the uterus is present, for cognitive support in women experiencing early menopause or premature ovarian insufficiency^53^. This is particularly the case following oophorectomy, where it is recommended to start hormone therapy as soon as possible after surgery and continue treatment until the average age of spontaneous menopause (around age 51), although the duration can vary based on individual factors and ongoing reassessment of benefits and risks. For women undergoing spontaneous menopause, as in this study, MHT is not recommended for AD risk reduction^53,54^. By identifying biological substrates of the menopause transition, present findings provide novel endpoints for clinical trials and observational research aimed at mitigating AD risk in menopausal women.

While our findings do not indicate that the women in our cohort have AD, evidence of AD brain biomarker phenotypes provide insights into the possible intersection of menopause and AD risk, supporting the notion that menopause-related hypoestrogenism may activate existing vulnerabilities to AD in some women^55^. Further research is warranted to simultaneously explore biomarkers of connectivity / diffusivity and AD pathology in this population. We caution that present results are cross-sectional and obtained from relatively small samples of carefully screened, highly educated participants at risk for AD. Replication in community-based populations with different risks and more diverse racial or socioeconomic backgrounds is warranted.

### Strengths and limitations

The present study has a number of strengths. First, we focused on carefully screened, healthy men and women ages 40-65 years, with comprehensive clinical and cognitive exams and menopause assessments. All of the individuals in this sample were cognitively normal at the time of participation, and our sample is enriched for AD risk factors, including APOE e4 carriership and family history. These factors increase the likelihood that the observed differences may be related to AD and therefore facilitate a midlife interpretation of potentially associated factors. Second, the fixel-based approach employed here is largely absent in the neuroimaging sex differences literature, providing novel information Finally, our hierarchical Bayesian approach enabled us to more completely characterize the probability of differences in these white matter metrics. This provides prior information to be incorporated systematically into future studies examining white matter characteristics in aging and disease.

Several limitations should be noted. First, the diffusion weighted imaging data collected were single-shell with a b-value of 1000mm^2^/s. Recommendations for fixel-based analyses include multi-shell data and higher b-values to better resolve crossing fibers^24^. However, past work, including some by the developers of the fixel-based metrics^25,56,57^, has successfully applied this to single-shell data, and therefore future studies may benefit from multi-shell data as an opportunity for refining the specificity of the findings here. Additionally, there are other contributors to white matter health that are not represented in these analyses, including lifestyle indicators (smoking, alcohol, exercise) as well as environmental considerations (socioeconomic status and childhood deprivation, pollution). Future work with larger samples should include these as potential factors that may alter the trajectory of white matter remodeling in midlife. Finally, cross-sectional data do not allow for precise causal inference about the effect of endocrinological aging in women. Future studies should include longitudinal studies that enable more precise modeling of the time course of these changes. Additional studies are needed in order to examine effects of menopause hormone therapy and as well as surgically induced menopause on the observed sex differences.

## Conclusions

Our findings demonstrate a dynamic trajectory of white matter remodeling across the menopausal transition, characterized by micro- and macrostructural changes during perimenopause. The dynamic restructuring of white matter complexity and integrity represents a critical period in the midlife female aging brain. As with all neural critical periods, there is the potential for unmasking of risk factors. Perimenopause has been previously identified as a critical window of intervention for Alzheimer’s disease risk reduction and our findings extend this need to white matter health as well. Identifying and tracking changes in white matter during endocrinological transition could facilitate timely interventions that preserve neuronal integrity and mitigate future cognitive decline.

## Methods

### Participants

This is a cross-sectional study of cognitively normal men and women ages 36-65 years, with a family history of late-onset AD and/or APOE-4 genotype. Participants were recruited at the Weill Cornell Medicine (WCM) Alzheimer’s Prevention Program between 2018-2024 by self-referral, flyers, and word of mouth.

Our inclusion and exclusion criteria were previously described^2–4^. Briefly, all participants had Montreal Cognitive Assessment (MoCA) score≥26 and normal cognitive test performance by age and education^2–4^. Exclusion criteria included medical conditions that may affect brain structure or function (e.g., stroke, any neurodegenerative diseases, major psychiatric disorders, hydrocephalus, demyelinating disorders such as Multiple Sclerosis, intracranial mass, and infarcts on MRI), use of psychoactive medications, and contraindications to brain imaging. All received medical, neurological, laboratory, cognitive and MRI exams, including volumetric MRI and DTI, within 3 months of each other.

A family history of late-onset AD was elicited using standardized questionnaires^2–4^. APOE-4 genotype was determined using standard qPCR procedures. Those carrying one or two APOE4 alleles were grouped as APOE-4 carriers.

The patients’ sex was determined by self-report. Our study protocol involves a ~1:3 enrollment ratio of men to women, with approximately equal representation of premenopausal, perimenopausal, and postmenopausal statuses among women. Determination of menopausal status was based on the Stages of Reproductive Aging Workshop (STRAW-10) criteria^58^ with hormone assessments as supportive criteria^5^. Participants were classified as premenopausal (regular cycler), perimenopausal (irregular cyclers with interval of amenorrhea ≥60 days or ≥2 skipped cycles) and postmenopausal (no cycle for ≥12 consecutive months)^5^. Information on hysterectomy/oophorectomy status and menopause hormone therapy (MHT) usage was obtained through review of medical history. MHT users and surgical menopausal patients were excluded.

### Image acquisition

All participants underwent MRI neuroimaging on a 3.0 Tesla G.E. Discovery MR750 scanner equipped with a 32-channel coil. Acquired sequences included a 3D sagittal brain volume imaging (BRAVO) sequence acquired with isotropic 1×1×1 mm resolution (TR: 8.2ms; TE: 3.2ms; TI: 450ms; Flip angle: 12°; FOV: 25.6cm on a 256 × 256 matrix; ARC acceleration). Diffusion Tensor Imaging (DTI) was acquired with one b=0 s/mm^2^ volume and 55 b=1000 s/mm^2^ directions (TR: 8s; TE: 65ms; acquisition matrix: 128×128 reconstructed to 256×256; resolution: 0.9×0.9×1.8 mm).

### Image minimal pre-processing

#### CAT12

To estimate total brain volume (TBV), all T1-weighted images were processed using CAT12 (v. 12.8.2) in SPM12 using default parameters^59^. Outputs from the process included spatially normalized gray matter, white matter, and CSF maps as well as tissue-type specific volume measures and total brain volume.

#### SynB0-DISCO

Prior to pre-processing the diffusion weighted images, an undistorted, diffusion image was synthesized from the b=0 diffusion image and the T1-weighted image using SyNB0-DISCO^60^. The purpose of this step was to enable susceptibility distortion correction.

#### QSIPrep

Pre-processed diffusion weighted data were produced using QSIPrep (v. 0.19.1)^61^, which is based on Nipype 1.8.6 (RRID:SCR_002502)^62,63^. Descriptions of the preprocessing steps are provided by boilerplates distributed with QSIPrep’s outputs under a CC0 license for the purpose of transparency and reproducibility in published works. These are presented below with minimal modifications for formatting and clarity.

#### Anatomical preprocessing

The T1-weighted (T1w) image was corrected for intensity non-uniformity (INU) using N4BiasFieldCorrection (ANTs 2.4.3)^64^, and used as an anatomical reference throughout the workflow. The anatomical reference image was reoriented into AC-PC alignment via a 6-DOF transform extracted from a full affine registration to the MNI152NLin2009cAsym template. A full nonlinear registration to the template from AC-PC space was estimated via symmetric nonlinear registration (SyN) using antsRegistration. Brain extraction was performed on the T1w image using SynthStrip^65^ and automated segmentation was performed using SynthSeg^66,67^ from FreeSurfer version 7.3.1.

#### Diffusion preprocessing

Any images with a b-value less than 100 s/mm^2 were treated as a b=0 image. MP-PCA denoising as implemented in MRtrix3’s *dwidenoise*^68,69^ was applied with a 5-voxel window. After MP-PCA, Gibbs unringing was performed using MRtrix3’s *mrdegibbs*^70^. Following unringing, the mean intensity of the DWI series was adjusted so all the mean intensity of the b=0 images matched across each separate DWI scanning sequence. B1 field inhomogeneity was corrected using *dwibiascorrect* from MRtrix3 with the N4 algorithm^64^ after corrected images were resampled.

FSL (version 6.0.5.1:57b01774)’s *eddy* was used for head motion correction and Eddy current correction^71^. Eddy was configured with a q-space smoothing factor of 10, a total of 5 iterations, and 1000 voxels used to estimate hyperparameters. A linear first level model and a linear second level model were used to characterize Eddy current-related spatial distortion. q-space coordinates were forcefully assigned to shells. Field offset was attempted to be separated from subject movement. Shells were aligned post-eddy. Eddy’s outlier replacement was run^72^. Data were grouped by slice, only including values from slices determined to contain at least 250 intracerebral voxels. Groups deviating by more than 4 standard deviations from the prediction had their data replaced with imputed values. Data was provided with reversed phase-encode blips, resulting in pairs of images with distortions going in opposite directions. Here, b=0 reference images with reversed phase encoding directions were used along with an equal number of b=0 images extracted from the DWI scans. From these pairs the susceptibility-induced off-resonance field was estimated using a method similar to that described in (Andersson, Skare, and Ashburner 2003)^73^. The fieldmaps were ultimately incorporated into the Eddy current and head motion correction interpolation. Final interpolation was performed using the *jac* method.

Several confounding time-series were calculated based on the preprocessed DWI: framewise displacement (FD) using the implementation in Nipype (following the definitions by Power et al. 2014)^74^. The head-motion estimates calculated in the correction step were also placed within the corresponding confounds file. Slicewise cross correlation was also calculated. The DWI time-series were resampled to ACPC, generating a preprocessed DWI run in ACPC space with 1mm isotropic voxels.

Many internal operations of QSIPrep use Nilearn 0.10.2 (RRID:SCR_001362)^75,76^ and DIPY^77^. For more details of the pipeline, see the section corresponding to workflows in QSIPrep’s documentation (https://qsiprep.readthedocs.io).

### Fixel-based analyses

The fixel-based analysis pipeline was implemented in MRtrix 3 (v. 3.0.4)^78^, has been described in detail in other publications^24,25^, and the present processing approach was adapted from previous publication^79^. Briefly, all QSIPrep preprocessed T1w and DWI volumes and vector files as well as the brain mask were reoriented to FSL standard orientation. The preprocessed DWI volumes were denoised using DIPY’s *Patch2Self* algorithm to minimize residual noise and improve SNR^80^. Response functions for grey matter, white matter, and CSF were estimated in MRtrix3 using the *dhollander* algorithm^81,82^ and an average response function for each tissue type was created across the whole dataset^34^. Fiber orientation directions (FODs) for each participant were generated using the *MRtrix3Tissue* Single-Shell 3-Tissue Constrained Spherical Deconvolution method^56^ (v. 5.2.9; https://github.com/3Tissue/MRtrix3Tissue) and log-domain intensity normalized^24,25^.

A randomly selected sample of 10 women in each menopausal transition state (pre-, peri-, post-) as well as 10 age-range matched men for each transition state were used to generate an unbiased, study-specific FOD template. All participant FODs were then independently registered and warped to the template brain along with their brain masks. A whole-brain fixel-wise analysis mask was created from the FOD template within the intersection of the warped brain masks using MRtrix’s fod2fixel using a peak FOD amplitude minimum threshold of 0.09 for inclusion. Participant’s warped FODs were then segmented, reoriented, and mapped to template space. For each participant, fiber density (FD), fiber cross-section (FC), and the product of FD and FC (FDC) were calculated for each fixel. FC was additionally log-transformed (logFC) prior to analyses due to a known non-normal distribution.

### Tract segmentation

The three primary spherical harmonic peaks from the FOD template brain were passed to TractSeg (v. 2.3)^83–86^. Tractograms were generated for 45 commissural and bilateral fiber bundles available in TractSeg. For each fiber bundle, the tractograms were converted to fixel masks using MRtrix’s tck2fixel. Finally mean values for FD, logFC, and FDC were calculated for each participant for each fiber bundle.

### DTI scalar preparation

From the reoriented DWI volumes, four diffusion tensor scalars were also computed (FA, MD, AD, and RD) using an iteratively weighted least squares approach (Mrtrix ref). These were computed in the participant’s native space, warped to the FOD template space, and then mapped to the TractSeg fiber bundle using TractSeg’s *Tractometry* function^86^. The *Tractometry* function divides each fiber bundle into 100 equally sized segments and computes the mean scalar value for each segment. For the present analyses, the mean value for the middle 98 segments (discarding the first and last segments) was computed as the summary statistic for each fiber bundle.

### Statistical analyses

All preprocessing on and analyses were conducted on the high performance computing cluster at the University of Arizona. Analyses were conducted using R (v. 4.4.0)^87^ and the *groundhog* package (v. 3.1.2)^88^ was used for package version control, loading packages from 2024-05-01. The primary aim was to identify sex-specific differences in fiber bundles micro- and macrostructural properties. We employed hierarchical Bayesian regression models to assess these differences at the level of both the whole brain and individual fiber bundles simultaneously. Five models were fit (FDC, logFC, FD, FA, and MD) using *brms*^89,90^, implementing 8 parallel chains and 10,000 iterations, plus an initial 5,000 tuning samples which were discarded, resulting in 80,000 total samples per model. The primary outcome of interest was the per-bundle differences between males and females. Each model incorporated previously identified predictors of white matter health, including age, years of education, APOE e4 carriership status (carrier vs. non-carrier), and family history of AD (positive vs. negative). Age, years of education, and total intracranial volume (FDC and logFC models only) were Z-score normalized before modeling.

Bundle-specific differences between sexes were identified from the posterior distributions of each model. These outcomes are summarized across the entire posterior distribution, presenting sex-specific mean values adjusted for all other covariates, along with the 95% highest density interval (HDI) of the between-sex difference and the proportion of differences greater than zero (P+). P+ provides a continuum of evidence, indicating the probability that the difference is in the observed direction^30,31^. As opposed to *p*-values, P+ is not interpreted through dichotomization (e.g., significant vs. not significant). Instead P+ in the present study indicates the probability of a female > male difference given the collected data. For example, P+ = 0.9 for FA in a tract indicates a 90% probability of female FA > male FA while P+ = 0.1 would indicate only a 10% probability, which implies that the probability of male FA > female FA is 90%. In line with the intention to provide a comprehensive view of sex differences in our data, descriptive statistics of all posterior distributions are provided in Supplementary Materials while the main reporting focuses on differences identified at both modest (P+ > 0.8, P+ < 0.2) and stronger (P+ > 0.9, P+ < 0.1) levels of evidence^91–93^.

Sensitivity analyses were then conducted to determine whether the observed sex differences could be attributed to menopausal status (pre-, peri-, and post-menopausal) or if they remained stable across the age range in our data, adjusting by the same confounders as above. To this end, the dataset was subdivided into three subsets, each containing women in a specific menopausal transition state and men matched to the same age range as the women in each of the menopausal status groups, consistent with prior research^2,6,94^. Due to overlapping age ranges for these transition state groups, a total of 20 men were included in more than one of the secondary analysis datasets; however these sensitivity datasets were analyzed separately and are thus treated as independent. The same modeling approach was applied to each subset, and results were reported similarly.

## Acknowledgements

This study was supported by grants from NIH/NIA (P01AG026572 and R01AG057931), NIH/NCATS UL1TR002384, the Cure Alzheimer’s Fund, the Women’s Alzheimer’s Movement, and philanthropic support to the Weill Cornell Alzheimer’s Prevention Program.

## Notes

### Competing Interest Statement

The authors have declared no competing interest.

