## Supplemental Tables for "White matter micro- and macrostructural properties in midlife individuals at risk for Alzheimer’s disease: Associations with sex and menopausal status"

### Supplemental table 1: Fiber density differences between women and men

| | Raw means[1] | | Covariate adjusted means[1] | | Hierarchical Bayesian model[1] | | |
|---|---|---|---|---|---|---|---|
| Tract | Females | Males | Females | Males | Mean Difference | 95% HDI | P+ |
| AF_left | 0.369±0.015 | 0.358±0.014 | 0.368±0.014 | 0.360±0.013 | 0.010 | [0.004, 0.015] | 0.998 |
| AF_right | 0.393±0.018 | 0.382±0.015 | 0.393±0.018 | 0.383±0.015 | 0.010 | [0.004, 0.015] | 0.998 |
| ATR_left | 0.367±0.017 | 0.359±0.018 | 0.366±0.017 | 0.361±0.015 | 0.009 | [0.003, 0.014] | 0.993 |
| ATR_right | 0.366±0.016 | 0.359±0.016 | 0.365±0.015 | 0.360±0.015 | 0.008 | [0.002, 0.013] | 0.987 |
| CA | 0.442±0.027 | 0.434±0.025 | 0.442±0.026 | 0.435±0.024 | 0.009 | [0.003, 0.014] | 0.994 |
| CC_1 | 0.454±0.020 | 0.435±0.018 | 0.453±0.019 | 0.438±0.017 | 0.014 | [0.009, 0.020] | 1.000 |
| CC_2 | 0.384±0.016 | 0.371±0.014 | 0.384±0.015 | 0.373±0.012 | 0.011 | [0.006, 0.017] | 0.999 |
| CC_3 | 0.402±0.022 | 0.384±0.021 | 0.401±0.022 | 0.386±0.019 | 0.014 | [0.008, 0.020] | 1.000 |
| CC_4 | 0.457±0.023 | 0.437±0.018 | 0.456±0.023 | 0.440±0.017 | 0.014 | [0.009, 0.020] | 1.000 |
| CC_5 | 0.433±0.021 | 0.412±0.018 | 0.432±0.021 | 0.414±0.016 | 0.015 | [0.009, 0.021] | 1.000 |
| CC_6 | 0.410±0.018 | 0.392±0.013 | 0.409±0.017 | 0.393±0.013 | 0.013 | [0.008, 0.019] | 1.000 |
| CC_7 | 0.387±0.017 | 0.372±0.013 | 0.387±0.017 | 0.373±0.012 | 0.012 | [0.007, 0.018] | 1.000 |
| CG_left | 0.352±0.014 | 0.345±0.014 | 0.352±0.014 | 0.346±0.013 | 0.007 | [0.002, 0.013] | 0.979 |
| CG_right | 0.365±0.017 | 0.355±0.013 | 0.364±0.017 | 0.357±0.012 | 0.009 | [0.003, 0.014] | 0.994 |
| CST_left | 0.553±0.020 | 0.549±0.019 | 0.553±0.020 | 0.550±0.019 | 0.006 | [0.000, 0.011] | 0.939 |
| CST_right | 0.519±0.019 | 0.512±0.017 | 0.519±0.019 | 0.513±0.017 | 0.007 | [0.001, 0.012] | 0.973 |
| FPT_left | 0.477±0.017 | 0.472±0.016 | 0.476±0.017 | 0.473±0.016 | 0.006 | [0.001, 0.012] | 0.959 |
| FPT_right | 0.467±0.017 | 0.460±0.018 | 0.467±0.017 | 0.461±0.018 | 0.007 | [0.001, 0.012] | 0.975 |
| FX_left | 0.523±0.059 | 0.486±0.068 | 0.520±0.056 | 0.495±0.056 | 0.027 | [0.021, 0.033] | 1.000 |
| FX_right | 0.536±0.061 | 0.492±0.070 | 0.532±0.056 | 0.502±0.057 | 0.033 | [0.027, 0.039] | 1.000 |
| ICP_left | 0.416±0.018 | 0.411±0.018 | 0.416±0.018 | 0.412±0.017 | 0.006 | [0.001, 0.012] | 0.961 |
| ICP_right | 0.381±0.018 | 0.373±0.015 | 0.381±0.018 | 0.374±0.014 | 0.008 | [0.003, 0.014] | 0.990 |
| IFO_left | 0.389±0.015 | 0.375±0.016 | 0.388±0.015 | 0.377±0.013 | 0.012 | [0.006, 0.017] | 1.000 |
| IFO_right | 0.386±0.016 | 0.372±0.012 | 0.385±0.016 | 0.373±0.011 | 0.011 | [0.006, 0.017] | 1.000 |
| ILF_left | 0.389±0.017 | 0.378±0.016 | 0.389±0.016 | 0.380±0.015 | 0.010 | [0.004, 0.015] | 0.998 |
| ILF_right | 0.393±0.018 | 0.385±0.018 | 0.393±0.018 | 0.386±0.017 | 0.008 | [0.003, 0.014] | 0.990 |
| MCP | 0.456±0.019 | 0.445±0.016 | 0.455±0.019 | 0.447±0.014 | 0.009 | [0.004, 0.015] | 0.995 |
| MLF_left | 0.377±0.016 | 0.362±0.013 | 0.377±0.016 | 0.364±0.012 | 0.011 | [0.006, 0.017] | 0.999 |
| MLF_right | 0.382±0.018 | 0.364±0.014 | 0.381±0.018 | 0.366±0.013 | 0.013 | [0.007, 0.019] | 1.000 |
| OR_left | 0.395±0.016 | 0.383±0.015 | 0.394±0.016 | 0.384±0.013 | 0.010 | [0.004, 0.015] | 0.997 |
| OR_right | 0.404±0.018 | 0.389±0.014 | 0.404±0.018 | 0.391±0.013 | 0.011 | [0.006, 0.017] | 0.999 |
| POPT_left | 0.464±0.017 | 0.452±0.014 | 0.464±0.017 | 0.453±0.014 | 0.010 | [0.004, 0.015] | 0.997 |
| POPT_right | 0.463±0.017 | 0.450±0.014 | 0.462±0.017 | 0.451±0.014 | 0.010 | [0.005, 0.016] | 0.998 |
| SCP_left | 0.464±0.017 | 0.468±0.015 | 0.464±0.016 | 0.467±0.015 | 0.001 | [-0.005, 0.007] | 0.622 |
| SCP_right | 0.440±0.015 | 0.440±0.015 | 0.440±0.015 | 0.440±0.015 | 0.003 | [-0.003, 0.008] | 0.790 |
| SLF_III_left | 0.370±0.017 | 0.360±0.017 | 0.369±0.016 | 0.362±0.015 | 0.010 | [0.004, 0.015] | 0.997 |
| SLF_III_right | 0.378±0.020 | 0.367±0.017 | 0.377±0.019 | 0.369±0.017 | 0.009 | [0.004, 0.015] | 0.997 |
| SLF_II_left | 0.338±0.017 | 0.325±0.016 | 0.337±0.017 | 0.326±0.016 | 0.011 | [0.005, 0.016] | 0.999 |
| SLF_II_right | 0.332±0.018 | 0.322±0.014 | 0.332±0.017 | 0.323±0.014 | 0.009 | [0.004, 0.015] | 0.996 |
| SLF_I_left | 0.310±0.015 | 0.305±0.013 | 0.310±0.015 | 0.306±0.013 | 0.006 | [0.001, 0.012] | 0.965 |
| SLF_I_right | 0.329±0.017 | 0.323±0.011 | 0.329±0.017 | 0.324±0.010 | 0.007 | [0.002, 0.013] | 0.979 |
| STR_left | 0.495±0.022 | 0.488±0.019 | 0.495±0.022 | 0.488±0.019 | 0.007 | [0.001, 0.013] | 0.978 |
| STR_right | 0.471±0.021 | 0.457±0.019 | 0.471±0.021 | 0.458±0.019 | 0.011 | [0.005, 0.016] | 0.999 |
| UF_left | 0.426±0.018 | 0.416±0.019 | 0.426±0.018 | 0.417±0.018 | 0.010 | [0.004, 0.015] | 0.998 |
| UF_right | 0.416±0.019 | 0.405±0.017 | 0.416±0.019 | 0.406±0.016 | 0.010 | [0.005, 0.016] | 0.999 |

[1] Units for raw and adjusted means, difference, and 95% HDI are arbitrary.

Data are presented as mean ± SD. Raw values are unadjusted sex-wise means. Adjusted values are sex-wise mean residual values after adjusting for age, education, APOE e4 carriership, and family history of Alzheimer's disease. Bayesian model outputs include the mean difference and 95% highest density interval estimates as well as P+. P+ indicates the proportion of the Bayesian samples (N = 80000) where the difference between women and men was > 0. P+ values approaching 1 indicate stronger evidence of female > male while P+ values approaching 0 indicate stronger evidence of male > female. The hierarchical model includes all tracts in a singular model and leverages partial pooling of the data, therefore the mean difference will not always be exactly equal to the difference in covariate adjusted means.

### Supplemental table 2: Fiber cross-section differences between women and men

| | Raw means[1] | | Covariate adjusted means[1] | | Hierarchical Bayesian model[1] | | |
|---|---|---|---|---|---|---|---|
| Tract | Females | Males | Females | Males | Mean Difference | 95% HDI | P+ |
| AF_left | 0.008±0.056 | 0.088±0.073 | 0.031±0.035 | 0.025±0.039 | 0.004 | [-0.009, 0.017] | 0.695 |
| AF_right | 0.006±0.055 | 0.089±0.073 | 0.029±0.032 | 0.026±0.042 | 0.004 | [-0.009, 0.017] | 0.708 |
| ATR_left | -0.017±0.070 | 0.069±0.082 | 0.009±0.047 | -0.001±0.043 | 0.008 | [-0.005, 0.022] | 0.833 |
| ATR_right | -0.013±0.070 | 0.073±0.083 | 0.012±0.048 | 0.004±0.053 | 0.007 | [-0.007, 0.020] | 0.784 |
| CA | -0.011±0.062 | 0.068±0.083 | 0.009±0.051 | 0.014±0.063 | 0.001 | [-0.012, 0.014] | 0.557 |
| CC_1 | -0.022±0.073 | 0.044±0.084 | 0.000±0.053 | -0.015±0.061 | 0.014 | [0.000, 0.029] | 0.945 |
| CC_2 | -0.003±0.066 | 0.074±0.081 | 0.021±0.043 | 0.008±0.047 | 0.010 | [-0.004, 0.023] | 0.867 |
| CC_3 | 0.010±0.079 | 0.087±0.101 | 0.035±0.065 | 0.020±0.067 | 0.006 | [-0.008, 0.020] | 0.757 |
| CC_4 | -0.012±0.063 | 0.059±0.081 | 0.007±0.055 | 0.006±0.058 | 0.003 | [-0.010, 0.016] | 0.641 |
| CC_5 | -0.019±0.059 | 0.051±0.066 | 0.000±0.046 | -0.002±0.044 | 0.005 | [-0.008, 0.018] | 0.745 |
| CC_6 | -0.026±0.052 | 0.057±0.070 | -0.003±0.033 | -0.003±0.048 | 0.003 | [-0.010, 0.015] | 0.627 |
| CC_7 | 0.017±0.070 | 0.129±0.092 | 0.043±0.058 | 0.058±0.070 | -0.009 | [-0.024, 0.006] | 0.162 |
| CG_left | -0.031±0.059 | 0.048±0.071 | -0.008±0.036 | -0.013±0.044 | 0.009 | [-0.004, 0.023] | 0.865 |
| CG_right | -0.027±0.056 | 0.051±0.066 | -0.005±0.034 | -0.007±0.043 | 0.007 | [-0.007, 0.020] | 0.791 |
| CST_left | -0.015±0.061 | 0.036±0.064 | 0.000±0.054 | -0.005±0.048 | 0.006 | [-0.007, 0.019] | 0.781 |
| CST_right | -0.021±0.059 | 0.036±0.071 | -0.004±0.051 | -0.010±0.052 | 0.006 | [-0.007, 0.019] | 0.763 |
| FPT_left | -0.016±0.052 | 0.046±0.067 | 0.003±0.037 | -0.004±0.041 | 0.009 | [-0.004, 0.023] | 0.870 |
| FPT_right | -0.011±0.055 | 0.047±0.069 | 0.008±0.041 | -0.004±0.043 | 0.009 | [-0.004, 0.022] | 0.863 |
| FX_left | -0.092±0.107 | 0.050±0.120 | -0.053±0.083 | -0.055±0.082 | -0.004 | [-0.019, 0.012] | 0.353 |
| FX_right | -0.092±0.104 | 0.055±0.124 | -0.055±0.084 | -0.046±0.088 | -0.009 | [-0.026, 0.007] | 0.179 |
| ICP_left | -0.027±0.064 | 0.018±0.055 | -0.014±0.056 | -0.017±0.050 | 0.008 | [-0.005, 0.022] | 0.830 |
| ICP_right | -0.026±0.061 | 0.017±0.056 | -0.013±0.053 | -0.019±0.049 | 0.008 | [-0.006, 0.021] | 0.818 |
| IFO_left | 0.005±0.057 | 0.097±0.071 | 0.029±0.037 | 0.032±0.036 | 0.001 | [-0.012, 0.014] | 0.543 |
| IFO_right | 0.005±0.055 | 0.103±0.073 | 0.029±0.038 | 0.038±0.050 | -0.003 | [-0.016, 0.011] | 0.372 |
| ILF_left | 0.023±0.057 | 0.120±0.072 | 0.048±0.038 | 0.054±0.040 | -0.004 | [-0.019, 0.009] | 0.307 |
| ILF_right | 0.016±0.061 | 0.121±0.077 | 0.042±0.042 | 0.049±0.046 | -0.005 | [-0.018, 0.009] | 0.302 |
| MCP | -0.012±0.063 | 0.038±0.061 | 0.002±0.055 | -0.001±0.052 | 0.005 | [-0.008, 0.019] | 0.746 |
| MLF_left | -0.007±0.052 | 0.077±0.066 | 0.015±0.031 | 0.016±0.041 | 0.001 | [-0.012, 0.014] | 0.560 |
| MLF_right | -0.017±0.052 | 0.069±0.070 | 0.006±0.034 | 0.009±0.045 | 0.002 | [-0.010, 0.016] | 0.622 |
| OR_left | 0.013±0.066 | 0.110±0.086 | 0.038±0.052 | 0.043±0.053 | -0.004 | [-0.018, 0.010] | 0.326 |
| OR_right | -0.004±0.066 | 0.102±0.087 | 0.021±0.054 | 0.034±0.069 | -0.006 | [-0.020, 0.008] | 0.254 |
| POPT_left | -0.025±0.051 | 0.038±0.057 | -0.006±0.037 | -0.011±0.037 | 0.005 | [-0.008, 0.018] | 0.752 |
| POPT_right | -0.028±0.050 | 0.033±0.060 | -0.011±0.037 | -0.015±0.042 | 0.006 | [-0.007, 0.019] | 0.778 |
| SCP_left | -0.030±0.049 | 0.009±0.057 | -0.017±0.041 | -0.026±0.044 | 0.010 | [-0.004, 0.024] | 0.880 |
| SCP_right | -0.032±0.048 | 0.009±0.055 | -0.019±0.041 | -0.026±0.043 | 0.009 | [-0.005, 0.022] | 0.860 |
| SLF_III_left | 0.005±0.061 | 0.091±0.086 | 0.029±0.043 | 0.026±0.059 | 0.003 | [-0.010, 0.016] | 0.624 |
| SLF_III_right | 0.009±0.060 | 0.084±0.076 | 0.030±0.041 | 0.026±0.057 | 0.005 | [-0.008, 0.018] | 0.727 |
| SLF_II_left | 0.017±0.061 | 0.090±0.073 | 0.039±0.041 | 0.029±0.042 | 0.005 | [-0.009, 0.018] | 0.708 |
| SLF_II_right | 0.010±0.065 | 0.095±0.075 | 0.034±0.044 | 0.030±0.045 | 0.004 | [-0.010, 0.017] | 0.670 |
| SLF_I_left | 0.009±0.053 | 0.072±0.066 | 0.028±0.040 | 0.023±0.044 | 0.003 | [-0.010, 0.017] | 0.655 |
| SLF_I_right | 0.000±0.056 | 0.073±0.064 | 0.021±0.039 | 0.018±0.042 | 0.004 | [-0.009, 0.017] | 0.693 |
| STR_left | -0.034±0.061 | 0.033±0.070 | -0.017±0.054 | -0.011±0.056 | 0.005 | [-0.009, 0.018] | 0.714 |
| STR_right | -0.019±0.057 | 0.034±0.073 | -0.004±0.050 | -0.007±0.053 | 0.007 | [-0.006, 0.020] | 0.799 |
| UF_left | -0.011±0.056 | 0.074±0.062 | 0.011±0.038 | 0.015±0.031 | 0.004 | [-0.010, 0.017] | 0.669 |
| UF_right | 0.002±0.053 | 0.083±0.066 | 0.023±0.036 | 0.026±0.041 | 0.002 | [-0.011, 0.015] | 0.617 |

[1] Units for raw and adjusted means, difference, and 95% HDI are arbitrary.

Data are presented as mean ± SD. Raw values are unadjusted sex-wise means. Adjusted values are sex-wise mean residual values after adjusting for age, education, intracranial volume, APOE e4 carriership, and family history of Alzheimer's disease. Bayesian model outputs include the mean difference and 95% highest density interval estimates as well as P+. P+ values approaching 1 indicate stronger evidence of female > male while P+ values approaching 0 indicate stronger evidence of male > female. The hierarchical model includes all tracts in a singular model and leverages partial pooling of the data, therefore the mean difference will not always be exactly equal to the difference in covariate adjusted means.

### Supplemental table 3: FDC differences between women and men

| | Raw means[1] | | Covariate adjusted means[1] | | Hierarchical Bayesian model[1] | | |
|---|---|---|---|---|---|---|---|
| Tract | Females | Males | Females | Males | Mean Difference | 95% HDI | P+ |
| AF_left | 0.375±0.029 | 0.394±0.035 | 0.382±0.024 | 0.375±0.024 | 0.007 | [-0.002, 0.017] | 0.888 |
| AF_right | 0.397±0.029 | 0.419±0.036 | 0.405±0.025 | 0.400±0.027 | 0.007 | [-0.003, 0.017] | 0.873 |
| ATR_left | 0.362±0.029 | 0.385±0.035 | 0.369±0.023 | 0.364±0.022 | 0.006 | [-0.003, 0.017] | 0.852 |
| ATR_right | 0.362±0.028 | 0.386±0.035 | 0.369±0.023 | 0.366±0.028 | 0.006 | [-0.004, 0.016] | 0.845 |
| CA | 0.438±0.042 | 0.466±0.053 | 0.445±0.039 | 0.446±0.045 | 0.004 | [-0.007, 0.014] | 0.712 |
| CC_1 | 0.447±0.039 | 0.453±0.048 | 0.453±0.035 | 0.440±0.040 | 0.011 | [0.000, 0.021] | 0.952 |
| CC_2 | 0.385±0.031 | 0.399±0.036 | 0.391±0.026 | 0.383±0.027 | 0.009 | [-0.001, 0.018] | 0.920 |
| CC_3 | 0.407±0.043 | 0.418±0.044 | 0.414±0.039 | 0.401±0.036 | 0.009 | [-0.001, 0.019] | 0.928 |
| CC_4 | 0.455±0.043 | 0.466±0.042 | 0.459±0.042 | 0.454±0.036 | 0.010 | [0.000, 0.020] | 0.941 |
| CC_5 | 0.427±0.038 | 0.434±0.035 | 0.430±0.037 | 0.424±0.029 | 0.010 | [0.000, 0.020] | 0.946 |
| CC_6 | 0.402±0.031 | 0.417±0.034 | 0.407±0.028 | 0.402±0.027 | 0.008 | [-0.002, 0.018] | 0.906 |
| CC_7 | 0.393±0.037 | 0.420±0.042 | 0.400±0.034 | 0.400±0.033 | 0.005 | [-0.005, 0.015] | 0.799 |
| CG_left | 0.343±0.025 | 0.364±0.032 | 0.349±0.021 | 0.348±0.027 | 0.007 | [-0.003, 0.017] | 0.859 |
| CG_right | 0.357±0.026 | 0.376±0.030 | 0.362±0.022 | 0.361±0.024 | 0.006 | [-0.003, 0.016] | 0.852 |
| CST_left | 0.547±0.044 | 0.568±0.040 | 0.554±0.042 | 0.549±0.033 | 0.007 | [-0.003, 0.018] | 0.882 |
| CST_right | 0.512±0.039 | 0.530±0.039 | 0.518±0.037 | 0.513±0.034 | 0.009 | [-0.001, 0.019] | 0.918 |
| FPT_left | 0.471±0.034 | 0.493±0.039 | 0.478±0.030 | 0.473±0.029 | 0.007 | [-0.002, 0.017] | 0.888 |
| FPT_right | 0.463±0.033 | 0.481±0.040 | 0.470±0.029 | 0.462±0.032 | 0.008 | [-0.002, 0.018] | 0.913 |
| FX_left | 0.472±0.044 | 0.501±0.057 | 0.478±0.042 | 0.487±0.049 | 0.001 | [-0.010, 0.012] | 0.575 |
| FX_right | 0.485±0.048 | 0.507±0.058 | 0.490±0.044 | 0.493±0.047 | 0.002 | [-0.009, 0.013] | 0.617 |
| ICP_left | 0.408±0.034 | 0.421±0.026 | 0.412±0.032 | 0.411±0.025 | 0.009 | [-0.001, 0.019] | 0.931 |
| ICP_right | 0.373±0.030 | 0.380±0.024 | 0.377±0.028 | 0.370±0.021 | 0.011 | [0.001, 0.021] | 0.959 |
| IFO_left | 0.390±0.028 | 0.410±0.032 | 0.396±0.024 | 0.392±0.020 | 0.006 | [-0.004, 0.016] | 0.844 |
| IFO_right | 0.385±0.028 | 0.407±0.031 | 0.392±0.025 | 0.390±0.022 | 0.006 | [-0.004, 0.016] | 0.848 |
| ILF_left | 0.398±0.030 | 0.424±0.036 | 0.405±0.026 | 0.405±0.025 | 0.004 | [-0.006, 0.014] | 0.756 |
| ILF_right | 0.398±0.035 | 0.431±0.040 | 0.407±0.029 | 0.405±0.028 | 0.004 | [-0.006, 0.014] | 0.731 |
| MCP | 0.455±0.040 | 0.468±0.034 | 0.460±0.038 | 0.454±0.030 | 0.010 | [0.000, 0.020] | 0.945 |
| MLF_left | 0.377±0.028 | 0.392±0.029 | 0.382±0.024 | 0.377±0.022 | 0.008 | [-0.002, 0.018] | 0.899 |
| MLF_right | 0.377±0.030 | 0.392±0.032 | 0.383±0.027 | 0.378±0.024 | 0.008 | [-0.002, 0.018] | 0.895 |
| OR_left | 0.397±0.032 | 0.422±0.037 | 0.404±0.028 | 0.402±0.026 | 0.005 | [-0.004, 0.015] | 0.808 |
| OR_right | 0.399±0.033 | 0.426±0.037 | 0.407±0.031 | 0.407±0.030 | 0.006 | [-0.004, 0.016] | 0.829 |
| POPT_left | 0.454±0.031 | 0.469±0.030 | 0.460±0.028 | 0.453±0.023 | 0.009 | [-0.001, 0.018] | 0.920 |
| POPT_right | 0.452±0.032 | 0.465±0.032 | 0.457±0.030 | 0.451±0.025 | 0.009 | [-0.001, 0.019] | 0.931 |
| SCP_left | 0.451±0.032 | 0.473±0.028 | 0.458±0.029 | 0.455±0.024 | 0.008 | [-0.002, 0.018] | 0.899 |
| SCP_right | 0.427±0.030 | 0.445±0.027 | 0.433±0.027 | 0.429±0.023 | 0.009 | [-0.001, 0.018] | 0.919 |
| SLF_III_left | 0.375±0.031 | 0.398±0.043 | 0.382±0.026 | 0.378±0.033 | 0.006 | [-0.003, 0.016] | 0.853 |
| SLF_III_right | 0.384±0.034 | 0.403±0.040 | 0.390±0.029 | 0.386±0.035 | 0.007 | [-0.002, 0.017] | 0.887 |
| SLF_II_left | 0.348±0.032 | 0.360±0.031 | 0.354±0.027 | 0.344±0.026 | 0.009 | [-0.001, 0.019] | 0.927 |
| SLF_II_right | 0.340±0.031 | 0.359±0.034 | 0.346±0.027 | 0.342±0.029 | 0.007 | [-0.003, 0.016] | 0.860 |
| SLF_I_left | 0.317±0.025 | 0.331±0.029 | 0.322±0.023 | 0.319±0.024 | 0.008 | [-0.002, 0.018] | 0.910 |
| SLF_I_right | 0.334±0.028 | 0.352±0.028 | 0.340±0.025 | 0.337±0.022 | 0.008 | [-0.002, 0.018] | 0.894 |
| STR_left | 0.483±0.043 | 0.503±0.040 | 0.489±0.041 | 0.485±0.034 | 0.007 | [-0.003, 0.017] | 0.876 |
| STR_right | 0.463±0.040 | 0.472±0.042 | 0.468±0.038 | 0.459±0.037 | 0.010 | [0.000, 0.020] | 0.941 |
| UF_left | 0.422±0.032 | 0.447±0.034 | 0.429±0.027 | 0.428±0.024 | 0.005 | [-0.005, 0.015] | 0.782 |
| UF_right | 0.417±0.031 | 0.439±0.035 | 0.423±0.028 | 0.422±0.026 | 0.005 | [-0.004, 0.015] | 0.815 |

[1] Units for raw and adjusted means, difference, and 95% HDI are arbitrary.

Data are presented as mean ± SD. Raw values are unadjusted sex-wise means. Adjusted values are sex-wise mean residual values after adjusting for age, education, intracranial volume, APOE e4 carriership, and family history of Alzheimer's disease. Bayesian model outputs include the mean difference and 95% highest density interval estimates as well as P+. P+ values approaching 1 indicate stronger evidence of female > male while P+ values approaching 0 indicate stronger evidence of male > female. The hierarchical model includes all tracts in a singular model and leverages partial pooling of the data, therefore the mean difference will not always be exactly equal to the difference in covariate adjusted means.

### Supplemental table 4: Fractional anisotropy differences between women and men

| Tract | Raw means[1] | | Covariate adjusted means[1] | | Hierarchical Bayesian model[1] | | |
|---|---|---|---|---|---|---|---|
| | Females | Males | Females | Males | Mean Difference | 95% HDI | P+ |
| AF_left | 0.387±0.015 | 0.385±0.017 | 0.387±0.015 | 0.386±0.016 | 0.001 | [-0.004, 0.005] | 0.608 |
| AF_right | 0.387±0.014 | 0.385±0.015 | 0.387±0.014 | 0.385±0.015 | 0.001 | [-0.004, 0.005] | 0.581 |
| ATR_left | 0.362±0.012 | 0.367±0.013 | 0.362±0.012 | 0.367±0.014 | -0.003 | [-0.007, 0.002] | 0.197 |
| ATR_right | 0.344±0.011 | 0.345±0.014 | 0.343±0.011 | 0.345±0.014 | -0.001 | [-0.006, 0.004] | 0.375 |
| CA | 0.393±0.046 | 0.396±0.052 | 0.393±0.046 | 0.396±0.052 | -0.002 | [-0.006, 0.003] | 0.297 |
| CC_1 | 0.435±0.018 | 0.435±0.023 | 0.435±0.017 | 0.436±0.022 | 0.001 | [-0.004, 0.006] | 0.652 |
| CC_2 | 0.411±0.017 | 0.407±0.016 | 0.410±0.016 | 0.408±0.015 | 0.002 | [-0.003, 0.007] | 0.744 |
| CC_3 | 0.403±0.016 | 0.400±0.016 | 0.403±0.015 | 0.400±0.016 | 0.002 | [-0.003, 0.006] | 0.721 |
| CC_4 | 0.450±0.017 | 0.442±0.016 | 0.450±0.016 | 0.443±0.015 | 0.004 | [-0.001, 0.009] | 0.914 |
| CC_5 | 0.451±0.018 | 0.442±0.023 | 0.451±0.018 | 0.443±0.021 | 0.005 | [0.000, 0.010] | 0.951 |
| CC_6 | 0.479±0.014 | 0.477±0.017 | 0.479±0.014 | 0.477±0.016 | 0.001 | [-0.004, 0.005] | 0.620 |
| CC_7 | 0.494±0.024 | 0.485±0.025 | 0.493±0.023 | 0.487±0.022 | 0.005 | [0.000, 0.010] | 0.947 |
| CG_left | 0.411±0.018 | 0.418±0.021 | 0.411±0.018 | 0.418±0.020 | -0.004 | [-0.008, 0.001] | 0.116 |
| CG_right | 0.401±0.016 | 0.407±0.022 | 0.401±0.016 | 0.407±0.022 | -0.002 | [-0.007, 0.002] | 0.207 |
| CST_left | 0.493±0.016 | 0.495±0.014 | 0.493±0.016 | 0.495±0.014 | -0.002 | [-0.007, 0.003] | 0.231 |
| CST_right | 0.486±0.015 | 0.491±0.013 | 0.487±0.014 | 0.491±0.013 | -0.003 | [-0.008, 0.001] | 0.131 |
| FPT_left | 0.458±0.014 | 0.465±0.014 | 0.458±0.014 | 0.464±0.014 | -0.004 | [-0.009, 0.001] | 0.079 |
| FPT_right | 0.449±0.014 | 0.455±0.016 | 0.449±0.014 | 0.454±0.016 | -0.004 | [-0.009, 0.001] | 0.086 |
| FX_left | 0.389±0.045 | 0.373±0.052 | 0.388±0.045 | 0.378±0.044 | 0.012 | [0.007, 0.017] | 1.000 |
| FX_right | 0.385±0.047 | 0.360±0.051 | 0.383±0.045 | 0.366±0.042 | 0.017 | [0.012, 0.022] | 1.000 |
| ICP_left | 0.414±0.014 | 0.421±0.019 | 0.414±0.014 | 0.421±0.019 | -0.004 | [-0.009, 0.001] | 0.107 |
| ICP_right | 0.421±0.017 | 0.423±0.019 | 0.421±0.017 | 0.424±0.018 | -0.002 | [-0.007, 0.003] | 0.257 |
| IFO_left | 0.420±0.014 | 0.417±0.016 | 0.419±0.014 | 0.417±0.014 | 0.001 | [-0.003, 0.006] | 0.669 |
| IFO_right | 0.426±0.015 | 0.424±0.016 | 0.426±0.015 | 0.425±0.015 | 0.001 | [-0.004, 0.006] | 0.636 |
| ILF_left | 0.412±0.017 | 0.408±0.019 | 0.412±0.017 | 0.409±0.017 | 0.002 | [-0.003, 0.007] | 0.745 |
| ILF_right | 0.411±0.016 | 0.409±0.014 | 0.411±0.016 | 0.410±0.013 | 0.000 | [-0.004, 0.005] | 0.520 |
| MCP | 0.480±0.013 | 0.483±0.016 | 0.480±0.012 | 0.483±0.016 | -0.003 | [-0.008, 0.002] | 0.163 |
| MLF_left | 0.385±0.014 | 0.380±0.015 | 0.385±0.014 | 0.381±0.014 | 0.002 | [-0.003, 0.007] | 0.732 |
| MLF_right | 0.376±0.014 | 0.374±0.015 | 0.376±0.014 | 0.374±0.015 | 0.000 | [-0.005, 0.005] | 0.521 |
| OR_left | 0.421±0.017 | 0.418±0.016 | 0.421±0.016 | 0.418±0.015 | 0.001 | [-0.004, 0.006] | 0.633 |
| OR_right | 0.430±0.016 | 0.428±0.017 | 0.430±0.016 | 0.428±0.016 | 0.001 | [-0.004, 0.005] | 0.601 |
| POPT_left | 0.453±0.013 | 0.455±0.012 | 0.453±0.013 | 0.454±0.012 | -0.002 | [-0.007, 0.002] | 0.220 |
| POPT_right | 0.459±0.013 | 0.463±0.013 | 0.459±0.013 | 0.463±0.013 | -0.003 | [-0.008, 0.002] | 0.133 |
| SCP_left | 0.430±0.014 | 0.439±0.012 | 0.430±0.013 | 0.438±0.012 | -0.007 | [-0.012, -0.002] | 0.010 |
| SCP_right | 0.409±0.014 | 0.420±0.017 | 0.410±0.013 | 0.419±0.015 | -0.008 | [-0.013, -0.003] | 0.004 |
| SLF_III_left | 0.395±0.015 | 0.394±0.016 | 0.394±0.015 | 0.395±0.014 | 0.001 | [-0.004, 0.005] | 0.605 |
| SLF_III_right | 0.385±0.014 | 0.386±0.016 | 0.385±0.013 | 0.386±0.015 | 0.000 | [-0.005, 0.005] | 0.507 |
| SLF_II_left | 0.353±0.017 | 0.345±0.018 | 0.353±0.017 | 0.345±0.017 | 0.003 | [-0.002, 0.008] | 0.836 |
| SLF_II_right | 0.359±0.017 | 0.354±0.014 | 0.359±0.016 | 0.354±0.014 | 0.001 | [-0.003, 0.006] | 0.679 |
| SLF_I_left | 0.377±0.016 | 0.376±0.016 | 0.377±0.015 | 0.376±0.015 | 0.000 | [-0.005, 0.005] | 0.522 |
| SLF_I_right | 0.376±0.016 | 0.373±0.014 | 0.376±0.015 | 0.373±0.014 | 0.000 | [-0.004, 0.005] | 0.551 |
| STR_left | 0.396±0.018 | 0.398±0.016 | 0.396±0.018 | 0.397±0.016 | -0.002 | [-0.007, 0.003] | 0.259 |
| STR_right | 0.398±0.017 | 0.402±0.018 | 0.398±0.017 | 0.402±0.018 | -0.003 | [-0.008, 0.002] | 0.167 |
| UF_left | 0.363±0.014 | 0.367±0.020 | 0.362±0.014 | 0.368±0.020 | -0.002 | [-0.007, 0.003] | 0.277 |
| UF_right | 0.353±0.013 | 0.354±0.017 | 0.353±0.013 | 0.355±0.017 | 0.000 | [-0.005, 0.005] | 0.462 |

[1] Units for raw and adjusted means, difference, and 95% HDI are arbitrary.

Data are presented as mean ± SD. Raw values are unadjusted sex-wise means. Adjusted values are sex-wise mean residual values after adjusting for age, education, APOE e4 carriership, and family history of Alzheimer's disease. Bayesian model outputs include the mean difference and 95% highest density interval estimates as well as P+. P+ indicates the proportion of the Bayesian samples (N = 80000) where the difference between women and men was > 0. P+ values approaching 1 indicate stronger evidence of female > male while P+ values approaching 0 indicate stronger evidence of male > female. The hierarchical model includes all tracts in a singular model and leverages partial pooling of the data, therefore the mean difference will not always be exactly equal to the difference in covariate adjusted means.

### Supplemental table 5: Mean diffusivity differences between women and men

| | Raw means[1] | | Covariate adjusted means[1] | | Hierarchical Bayesian model[1] | | |
|---|---|---|---|---|---|---|---|
| Tract | Females | Males | Females | Males | Mean Difference | 95% HDI | P+ |
| AF_left | 7.128±0.205 | 7.137±0.206 | 7.130±0.204 | 7.133±0.195 | -0.004 | [-0.095, 0.088] | 0.476 |
| AF_right | 7.192±0.187 | 7.154±0.231 | 7.191±0.184 | 7.157±0.220 | 0.010 | [-0.079, 0.104] | 0.568 |
| ATR_left | 7.549±0.217 | 7.597±0.226 | 7.553±0.215 | 7.586±0.211 | -0.031 | [-0.121, 0.062] | 0.294 |
| ATR_right | 7.821±0.205 | 7.890±0.245 | 7.826±0.203 | 7.877±0.228 | -0.042 | [-0.133, 0.049] | 0.226 |
| CA | 8.130±0.359 | 8.088±0.375 | 8.136±0.359 | 8.071±0.355 | -0.013 | [-0.107, 0.081] | 0.402 |
| CC_1 | 8.136±0.326 | 8.070±0.326 | 8.136±0.321 | 8.070±0.328 | -0.009 | [-0.103, 0.085] | 0.431 |
| CC_2 | 8.451±0.362 | 8.512±0.263 | 8.452±0.360 | 8.508±0.259 | -0.054 | [-0.147, 0.039] | 0.172 |
| CC_3 | 8.723±0.409 | 8.867±0.349 | 8.727±0.405 | 8.856±0.339 | -0.091 | [-0.184, 0.003] | 0.059 |
| CC_4 | 8.374±0.366 | 8.526±0.340 | 8.382±0.366 | 8.503±0.306 | -0.087 | [-0.178, 0.005] | 0.063 |
| CC_5 | 8.786±0.421 | 9.014±0.562 | 8.804±0.419 | 8.967±0.496 | -0.128 | [-0.221, -0.036] | 0.013 |
| CC_6 | 7.888±0.263 | 7.957±0.311 | 7.895±0.261 | 7.937±0.283 | -0.048 | [-0.140, 0.042] | 0.197 |
| CC_7 | 9.113±0.658 | 9.498±0.796 | 9.134±0.644 | 9.442±0.765 | -0.189 | [-0.286, -0.090] | 0.001 |
| CG_left | 7.273±0.194 | 7.232±0.243 | 7.271±0.191 | 7.239±0.237 | 0.011 | [-0.080, 0.103] | 0.577 |
| CG_right | 7.245±0.201 | 7.209±0.244 | 7.244±0.197 | 7.213±0.235 | 0.010 | [-0.083, 0.101] | 0.570 |
| CST_left | 6.954±0.160 | 6.958±0.127 | 6.953±0.159 | 6.960±0.121 | 0.007 | [-0.085, 0.098] | 0.552 |
| CST_right | 7.103±0.138 | 7.046±0.156 | 7.102±0.137 | 7.051±0.152 | 0.021 | [-0.070, 0.112] | 0.644 |
| FPT_left | 7.138±0.156 | 7.119±0.152 | 7.136±0.155 | 7.125±0.147 | 0.010 | [-0.080, 0.102] | 0.570 |
| FPT_right | 7.340±0.163 | 7.284±0.171 | 7.337±0.162 | 7.293±0.166 | 0.016 | [-0.076, 0.107] | 0.606 |
| FX_left | 15.978±1.810 | 16.751±2.067 | 16.062±1.739 | 16.525±1.816 | -0.551 | [-0.673, -0.433] | 0.000 |
| FX_right | 15.773±1.790 | 16.777±1.990 | 15.871±1.680 | 16.510±1.653 | -0.641 | [-0.764, -0.521] | 0.000 |
| ICP_left | 7.138±0.177 | 7.063±0.184 | 7.136±0.174 | 7.066±0.190 | 0.029 | [-0.064, 0.120] | 0.692 |
| ICP_right | 6.890±0.205 | 6.820±0.202 | 6.888±0.198 | 6.824±0.201 | 0.039 | [-0.054, 0.131] | 0.749 |
| IFO_left | 7.829±0.264 | 7.896±0.273 | 7.832±0.264 | 7.888±0.258 | -0.040 | [-0.130, 0.052] | 0.240 |
| IFO_right | 7.863±0.266 | 7.954±0.321 | 7.870±0.263 | 7.936±0.305 | -0.050 | [-0.140, 0.040] | 0.189 |
| ILF_left | 7.811±0.307 | 7.917±0.300 | 7.815±0.306 | 7.905±0.271 | -0.052 | [-0.144, 0.039] | 0.179 |
| ILF_right | 7.706±0.284 | 7.828±0.299 | 7.713±0.277 | 7.810±0.285 | -0.055 | [-0.145, 0.038] | 0.166 |
| MCP | 6.892±0.145 | 6.826±0.174 | 6.888±0.142 | 6.836±0.164 | 0.038 | [-0.054, 0.130] | 0.747 |
| MLF_left | 7.579±0.218 | 7.620±0.245 | 7.581±0.213 | 7.617±0.234 | -0.024 | [-0.115, 0.067] | 0.337 |
| MLF_right | 7.403±0.210 | 7.373±0.274 | 7.403±0.209 | 7.374±0.257 | 0.000 | [-0.091, 0.091] | 0.499 |
| OR_left | 8.113±0.354 | 8.376±0.503 | 8.124±0.356 | 8.345±0.471 | -0.109 | [-0.204, -0.013] | 0.029 |
| OR_right | 7.921±0.330 | 8.082±0.441 | 7.931±0.324 | 8.057±0.414 | -0.081 | [-0.173, 0.012] | 0.078 |
| POPT_left | 7.635±0.174 | 7.692±0.183 | 7.638±0.172 | 7.684±0.171 | -0.029 | [-0.120, 0.062] | 0.303 |
| POPT_right | 7.618±0.156 | 7.618±0.220 | 7.620±0.155 | 7.612±0.211 | -0.012 | [-0.104, 0.078] | 0.416 |
| SCP_left | 8.028±0.220 | 8.081±0.229 | 8.026±0.216 | 8.087±0.228 | -0.026 | [-0.120, 0.068] | 0.322 |
| SCP_right | 8.303±0.243 | 8.286±0.238 | 8.301±0.239 | 8.294±0.242 | -0.016 | [-0.112, 0.081] | 0.386 |
| SLF_III_left | 7.091±0.225 | 7.104±0.216 | 7.093±0.224 | 7.098±0.200 | -0.006 | [-0.098, 0.087] | 0.464 |
| SLF_III_right | 7.336±0.212 | 7.317±0.246 | 7.336±0.211 | 7.317±0.232 | 0.001 | [-0.090, 0.091] | 0.503 |
| SLF_II_left | 7.084±0.191 | 7.115±0.227 | 7.085±0.187 | 7.113±0.219 | -0.008 | [-0.101, 0.083] | 0.447 |
| SLF_II_right | 7.208±0.188 | 7.201±0.239 | 7.207±0.183 | 7.205±0.232 | 0.000 | [-0.091, 0.093] | 0.504 |
| SLF_I_left | 7.282±0.216 | 7.289±0.255 | 7.284±0.209 | 7.284±0.249 | -0.011 | [-0.103, 0.081] | 0.423 |
| SLF_I_right | 7.384±0.216 | 7.380±0.269 | 7.384±0.208 | 7.380±0.266 | -0.008 | [-0.099, 0.084] | 0.446 |
| STR_left | 7.020±0.192 | 7.050±0.183 | 7.021±0.189 | 7.048±0.169 | -0.007 | [-0.100, 0.084] | 0.454 |
| STR_right | 7.027±0.162 | 6.977±0.196 | 7.026±0.159 | 6.979±0.182 | 0.016 | [-0.076, 0.107] | 0.607 |
| UF_left | 7.675±0.236 | 7.562±0.227 | 7.673±0.234 | 7.567±0.227 | 0.021 | [-0.073, 0.115] | 0.632 |
| UF_right | 7.895±0.211 | 7.854±0.191 | 7.895±0.209 | 7.854±0.186 | -0.005 | [-0.098, 0.088] | 0.460 |

[1] Units for raw and adjusted means, difference, and 95% HDI are $10^{-4}$ mm²/s.

Data are presented as mean ± SD. Raw values are unadjusted sex-wise means. Adjusted values are sex-wise mean residual values after adjusting for age, education, APOE e4 carriership, and family history of Alzheimer's disease. Bayesian model outputs include the mean difference and 95% highest density interval estimates as well as P+. P+ indicates the proportion of the Bayesian samples (N = 80000) where the difference between women and men was > 0. P+ values approaching 1 indicate stronger evidence of female > male while P+ values approaching 0 indicate stronger evidence of male > female. The hierarchical model includes all tracts in a singular model and leverages partial pooling of the data, therefore the mean difference will not always be exactly equal to the difference in covariate adjusted means.

### Supplemental table 6: Fiber density differences between pre-menopausal women and age-matched men

| | Raw means[1] | | Covariate adjusted means[1] | | Hierarchical Bayesian model[1] | | |
|---|---|---|---|---|---|---|---|
| Tract | Females | Males | Females | Males | Mean Difference | 95% HDI | P+ |
| AF_left | 0.368±0.014 | 0.360±0.014 | 0.368±0.013 | 0.361±0.012 | 0.009 | [0.001, 0.017] | 0.960 |
| AF_right | 0.392±0.016 | 0.381±0.016 | 0.391±0.016 | 0.382±0.012 | 0.011 | [0.003, 0.019] | 0.985 |
| ATR_left | 0.368±0.017 | 0.361±0.015 | 0.368±0.016 | 0.362±0.012 | 0.008 | [0.000, 0.016] | 0.948 |
| ATR_right | 0.366±0.016 | 0.358±0.013 | 0.366±0.016 | 0.358±0.012 | 0.008 | [0.000, 0.016] | 0.942 |
| CA | 0.447±0.025 | 0.440±0.022 | 0.447±0.025 | 0.439±0.021 | 0.007 | [-0.001, 0.015] | 0.909 |
| CC_1 | 0.458±0.021 | 0.436±0.018 | 0.457±0.020 | 0.437±0.018 | 0.016 | [0.008, 0.025] | 0.999 |
| CC_2 | 0.386±0.016 | 0.371±0.012 | 0.385±0.016 | 0.373±0.011 | 0.012 | [0.004, 0.020] | 0.989 |
| CC_3 | 0.403±0.022 | 0.384±0.018 | 0.402±0.022 | 0.386±0.015 | 0.015 | [0.007, 0.023] | 0.998 |
| CC_4 | 0.455±0.025 | 0.436±0.017 | 0.455±0.025 | 0.437±0.013 | 0.015 | [0.007, 0.024] | 0.999 |
| CC_5 | 0.434±0.023 | 0.412±0.019 | 0.434±0.023 | 0.413±0.015 | 0.016 | [0.008, 0.024] | 0.999 |
| CC_6 | 0.410±0.018 | 0.392±0.013 | 0.409±0.018 | 0.394±0.012 | 0.014 | [0.006, 0.022] | 0.997 |
| CC_7 | 0.389±0.019 | 0.376±0.011 | 0.389±0.019 | 0.377±0.010 | 0.011 | [0.002, 0.019] | 0.982 |
| CG_left | 0.353±0.015 | 0.348±0.014 | 0.352±0.014 | 0.349±0.014 | 0.006 | [-0.002, 0.014] | 0.876 |
| CG_right | 0.365±0.017 | 0.358±0.013 | 0.364±0.017 | 0.360±0.012 | 0.008 | [-0.001, 0.016] | 0.931 |
| CST_left | 0.550±0.018 | 0.544±0.016 | 0.550±0.018 | 0.545±0.015 | 0.009 | [0.001, 0.018] | 0.963 |
| CST_right | 0.518±0.017 | 0.508±0.015 | 0.517±0.017 | 0.508±0.014 | 0.010 | [0.002, 0.019] | 0.977 |
| FPT_left | 0.476±0.016 | 0.470±0.016 | 0.475±0.016 | 0.471±0.015 | 0.008 | [0.000, 0.016] | 0.943 |
| FPT_right | 0.466±0.018 | 0.457±0.017 | 0.466±0.017 | 0.458±0.016 | 0.010 | [0.002, 0.018] | 0.972 |
| FX_left | 0.535±0.050 | 0.505±0.059 | 0.532±0.049 | 0.510±0.052 | 0.026 | [0.018, 0.035] | 1.000 |
| FX_right | 0.552±0.053 | 0.507±0.068 | 0.549±0.053 | 0.512±0.063 | 0.033 | [0.023, 0.041] | 1.000 |
| ICP_left | 0.415±0.016 | 0.411±0.017 | 0.415±0.015 | 0.411±0.015 | 0.006 | [-0.003, 0.014] | 0.866 |
| ICP_right | 0.380±0.017 | 0.371±0.016 | 0.379±0.016 | 0.371±0.014 | 0.009 | [0.000, 0.017] | 0.954 |
| IFO_left | 0.391±0.015 | 0.378±0.014 | 0.391±0.016 | 0.379±0.011 | 0.012 | [0.003, 0.020] | 0.988 |
| IFO_right | 0.388±0.016 | 0.373±0.011 | 0.387±0.016 | 0.374±0.009 | 0.012 | [0.003, 0.020] | 0.988 |
| ILF_left | 0.390±0.016 | 0.382±0.014 | 0.390±0.016 | 0.382±0.013 | 0.008 | [0.000, 0.016] | 0.949 |
| ILF_right | 0.396±0.016 | 0.385±0.016 | 0.395±0.016 | 0.386±0.013 | 0.010 | [0.002, 0.018] | 0.975 |
| MCP | 0.456±0.019 | 0.447±0.015 | 0.455±0.018 | 0.447±0.012 | 0.010 | [0.002, 0.018] | 0.973 |
| MLF_left | 0.379±0.015 | 0.363±0.012 | 0.378±0.014 | 0.365±0.011 | 0.012 | [0.004, 0.020] | 0.989 |
| MLF_right | 0.382±0.017 | 0.364±0.014 | 0.381±0.017 | 0.366±0.011 | 0.014 | [0.005, 0.022] | 0.996 |
| OR_left | 0.395±0.017 | 0.384±0.014 | 0.395±0.018 | 0.385±0.011 | 0.010 | [0.002, 0.018] | 0.977 |
| OR_right | 0.404±0.018 | 0.389±0.014 | 0.403±0.018 | 0.390±0.011 | 0.012 | [0.004, 0.020] | 0.991 |
| POPT_left | 0.465±0.015 | 0.450±0.015 | 0.464±0.014 | 0.451±0.015 | 0.013 | [0.004, 0.021] | 0.993 |
| POPT_right | 0.462±0.016 | 0.445±0.012 | 0.461±0.016 | 0.446±0.011 | 0.013 | [0.005, 0.022] | 0.995 |
| SCP_left | 0.462±0.016 | 0.463±0.015 | 0.462±0.016 | 0.463±0.014 | 0.005 | [-0.004, 0.013] | 0.815 |
| SCP_right | 0.438±0.015 | 0.435±0.014 | 0.437±0.015 | 0.435±0.013 | 0.006 | [-0.002, 0.014] | 0.877 |
| SLF_III_left | 0.368±0.016 | 0.363±0.015 | 0.368±0.016 | 0.364±0.012 | 0.008 | [0.000, 0.016] | 0.932 |
| SLF_III_right | 0.379±0.019 | 0.368±0.016 | 0.378±0.018 | 0.369±0.015 | 0.010 | [0.002, 0.018] | 0.975 |
| SLF_II_left | 0.339±0.017 | 0.325±0.013 | 0.339±0.017 | 0.325±0.013 | 0.011 | [0.003, 0.019] | 0.983 |
| SLF_II_right | 0.331±0.016 | 0.323±0.014 | 0.330±0.016 | 0.324±0.011 | 0.009 | [0.001, 0.017] | 0.955 |
| SLF_I_left | 0.311±0.014 | 0.306±0.015 | 0.311±0.013 | 0.306±0.014 | 0.005 | [-0.003, 0.014] | 0.850 |
| SLF_I_right | 0.326±0.015 | 0.323±0.011 | 0.326±0.015 | 0.324±0.010 | 0.005 | [-0.003, 0.014] | 0.849 |
| STR_left | 0.495±0.020 | 0.485±0.017 | 0.494±0.020 | 0.486±0.016 | 0.010 | [0.002, 0.018] | 0.976 |
| STR_right | 0.473±0.019 | 0.451±0.016 | 0.472±0.018 | 0.452±0.016 | 0.017 | [0.009, 0.025] | 0.999 |
| UF_left | 0.430±0.019 | 0.420±0.015 | 0.429±0.019 | 0.420±0.015 | 0.010 | [0.002, 0.018] | 0.970 |
| UF_right | 0.418±0.019 | 0.408±0.014 | 0.418±0.019 | 0.409±0.014 | 0.009 | [0.001, 0.018] | 0.969 |

[1] Units for raw and adjusted means, difference, and 95% HDI are arbitrary.

Data are presented as mean ± SD. Raw values are unadjusted sex-wise means. Adjusted values are sex-wise mean residual values after adjusting for age, education, APOE e4 carriership, and family history of Alzheimer's disease. Bayesian model outputs include the mean difference and 95% highest density interval estimates as well as P+. P+ indicates the proportion of the Bayesian samples (N = 80000) where the difference between women and men was > 0. P+ values approaching 1 indicate stronger evidence of female > male while P+ values approaching 0 indicate stronger evidence of male > female. The hierarchical model includes all tracts in a singular model and leverages partial pooling of the data, therefore the mean difference will not always be exactly equal to the difference in covariate adjusted means.

### Supplemental table 7: Fiber cross-section differences between pre-menopausal women and age-matched men

| Tract | Raw means[1] | | Covariate adjusted means[1] | | Hierarchical Bayesian model[1] | | |
|---|---|---|---|---|---|---|---|
| | Females | Males | Females | Males | Mean Difference | 95% HDI | P+ |
| AF_left | 0.017±0.058 | 0.111±0.067 | 0.051±0.029 | 0.047±0.034 | 0.002 | [-0.023, 0.026] | 0.549 |
| AF_right | 0.012±0.059 | 0.109±0.068 | 0.046±0.033 | 0.044±0.039 | 0.000 | [-0.025, 0.024] | 0.490 |
| ATR_left | -0.009±0.077 | 0.084±0.077 | 0.029±0.043 | 0.012±0.044 | 0.011 | [-0.014, 0.037] | 0.761 |
| ATR_right | -0.003±0.079 | 0.089±0.089 | 0.036±0.049 | 0.016±0.054 | 0.012 | [-0.013, 0.038] | 0.779 |
| CA | -0.001±0.066 | 0.094±0.082 | 0.030±0.057 | 0.037±0.062 | -0.008 | [-0.033, 0.017] | 0.301 |
| CC_1 | -0.012±0.074 | 0.058±0.093 | 0.021±0.048 | -0.004±0.064 | 0.023 | [-0.003, 0.049] | 0.924 |
| CC_2 | 0.004±0.073 | 0.086±0.078 | 0.039±0.047 | 0.020±0.044 | 0.014 | [-0.011, 0.040] | 0.814 |
| CC_3 | 0.019±0.094 | 0.110±0.096 | 0.056±0.073 | 0.040±0.061 | 0.009 | [-0.016, 0.035] | 0.718 |
| CC_4 | -0.014±0.069 | 0.069±0.065 | 0.013±0.057 | 0.018±0.050 | 0.000 | [-0.025, 0.025] | 0.497 |
| CC_5 | -0.019±0.056 | 0.062±0.061 | 0.009±0.040 | 0.010±0.046 | 0.003 | [-0.021, 0.028] | 0.581 |
| CC_6 | -0.026±0.050 | 0.078±0.072 | 0.007±0.033 | 0.016±0.046 | -0.007 | [-0.032, 0.018] | 0.318 |
| CC_7 | 0.008±0.068 | 0.160±0.082 | 0.049±0.063 | 0.083±0.054 | -0.036 | [-0.064, -0.008] | 0.017 |
| CG_left | -0.030±0.060 | 0.064±0.076 | 0.004±0.032 | 0.000±0.044 | 0.006 | [-0.019, 0.031] | 0.643 |
| CG_right | -0.020±0.060 | 0.059±0.075 | 0.011±0.032 | 0.001±0.041 | 0.011 | [-0.014, 0.037] | 0.763 |
| CST_left | -0.010±0.067 | 0.043±0.061 | 0.011±0.055 | 0.003±0.054 | 0.013 | [-0.012, 0.038] | 0.799 |
| CST_right | -0.022±0.068 | 0.045±0.068 | 0.002±0.056 | -0.001±0.051 | 0.007 | [-0.017, 0.032] | 0.683 |
| FPT_left | -0.013±0.057 | 0.059±0.059 | 0.014±0.041 | 0.008±0.035 | 0.009 | [-0.016, 0.033] | 0.717 |
| FPT_right | -0.006±0.064 | 0.055±0.059 | 0.021±0.046 | 0.005±0.028 | 0.016 | [-0.009, 0.041] | 0.851 |
| FX_left | -0.095±0.105 | 0.051±0.102 | -0.046±0.077 | -0.040±0.076 | -0.012 | [-0.039, 0.015] | 0.247 |
| FX_right | -0.095±0.098 | 0.058±0.112 | -0.046±0.077 | -0.035±0.082 | -0.019 | [-0.046, 0.008] | 0.135 |
| ICP_left | -0.038±0.064 | 0.021±0.050 | -0.018±0.050 | -0.016±0.054 | 0.004 | [-0.022, 0.029] | 0.589 |
| ICP_right | -0.031±0.060 | 0.012±0.058 | -0.012±0.044 | -0.023±0.054 | 0.014 | [-0.012, 0.040] | 0.811 |
| IFO_left | 0.011±0.056 | 0.118±0.073 | 0.048±0.033 | 0.049±0.027 | -0.005 | [-0.030, 0.020] | 0.384 |
| IFO_right | 0.000±0.054 | 0.132±0.071 | 0.038±0.040 | 0.059±0.041 | -0.022 | [-0.047, 0.004] | 0.087 |
| ILF_left | 0.033±0.055 | 0.144±0.077 | 0.068±0.044 | 0.077±0.029 | -0.013 | [-0.038, 0.013] | 0.206 |
| ILF_right | 0.015±0.063 | 0.143±0.079 | 0.054±0.048 | 0.070±0.042 | -0.018 | [-0.043, 0.008] | 0.126 |
| MCP | -0.015±0.060 | 0.031±0.063 | 0.004±0.047 | -0.004±0.056 | 0.011 | [-0.015, 0.036] | 0.752 |
| MLF_left | 0.001±0.054 | 0.098±0.066 | 0.033±0.032 | 0.037±0.040 | -0.004 | [-0.029, 0.021] | 0.403 |
| MLF_right | -0.011±0.050 | 0.092±0.074 | 0.021±0.034 | 0.031±0.045 | -0.008 | [-0.033, 0.017] | 0.309 |
| OR_left | 0.019±0.070 | 0.133±0.091 | 0.057±0.057 | 0.063±0.047 | -0.011 | [-0.036, 0.014] | 0.242 |
| OR_right | -0.016±0.068 | 0.140±0.074 | 0.024±0.062 | 0.064±0.056 | -0.039 | [-0.066, -0.010] | 0.012 |
| POPT_left | -0.024±0.053 | 0.053±0.059 | 0.004±0.030 | 0.000±0.039 | 0.005 | [-0.019, 0.030] | 0.629 |
| POPT_right | -0.029±0.050 | 0.051±0.063 | -0.002±0.032 | 0.000±0.046 | 0.002 | [-0.023, 0.026] | 0.546 |
| SCP_left | -0.033±0.056 | 0.021±0.053 | -0.011±0.042 | -0.019±0.044 | 0.013 | [-0.013, 0.037] | 0.790 |
| SCP_right | -0.033±0.055 | 0.021±0.053 | -0.012±0.041 | -0.018±0.044 | 0.011 | [-0.014, 0.036] | 0.760 |
| SLF_III_left | 0.008±0.065 | 0.112±0.096 | 0.045±0.042 | 0.043±0.064 | 0.001 | [-0.024, 0.025] | 0.518 |
| SLF_III_right | 0.017±0.063 | 0.098±0.078 | 0.048±0.039 | 0.039±0.058 | 0.008 | [-0.017, 0.033] | 0.702 |
| SLF_II_left | 0.030±0.065 | 0.105±0.069 | 0.060±0.039 | 0.049±0.044 | 0.011 | [-0.015, 0.035] | 0.762 |
| SLF_II_right | 0.021±0.075 | 0.118±0.075 | 0.055±0.051 | 0.053±0.044 | 0.003 | [-0.022, 0.028] | 0.567 |
| SLF_I_left | 0.010±0.054 | 0.081±0.062 | 0.035±0.041 | 0.034±0.041 | 0.006 | [-0.019, 0.030] | 0.644 |
| SLF_I_right | 0.001±0.063 | 0.092±0.063 | 0.031±0.046 | 0.035±0.040 | 0.000 | [-0.025, 0.024] | 0.493 |
| STR_left | -0.028±0.059 | 0.044±0.068 | -0.005±0.049 | 0.001±0.058 | 0.004 | [-0.021, 0.029] | 0.601 |
| STR_right | -0.013±0.057 | 0.046±0.072 | 0.009±0.046 | 0.004±0.055 | 0.012 | [-0.013, 0.037] | 0.775 |
| UF_left | -0.003±0.055 | 0.095±0.061 | 0.029±0.039 | 0.035±0.026 | -0.004 | [-0.028, 0.021] | 0.407 |
| UF_right | 0.005±0.058 | 0.096±0.072 | 0.038±0.037 | 0.034±0.040 | 0.002 | [-0.023, 0.026] | 0.544 |

[1] Units for raw and adjusted means, difference, and 95% HDI are arbitrary.

Data are presented as mean ± SD. Raw values are unadjusted sex-wise means. Adjusted values are sex-wise mean residual values after adjusting for age, education, intracranial volume, APOE e4 carriership, and family history of Alzheimer's disease. Bayesian model outputs include the mean difference and 95% highest density interval estimates as well as P+. P+ values approaching 1 indicate stronger evidence of female > male while P+ values approaching 0 indicate stronger evidence of male > female. The hierarchical model includes all tracts in a singular model and leverages partial pooling of the data, therefore the mean difference will not always be exactly equal to the difference in covariate adjusted means.

### Supplemental table 8: FDC differences between pre-menopausal women and age-matched men

| | Raw means[1] | | Covariate adjusted means[1] | | Hierarchical Bayesian model[1] | | |
|---|---|---|---|---|---|---|---|
| Tract | Females | Males | Females | Males | Mean Difference | 95% HDI | P+ |
| AF_left | 0.378±0.026 | 0.406±0.035 | 0.389±0.020 | 0.385±0.023 | 0.004 | [-0.013, 0.021] | 0.658 |
| AF_right | 0.399±0.026 | 0.427±0.038 | 0.409±0.022 | 0.407±0.026 | 0.003 | [-0.014, 0.020] | 0.620 |
| ATR_left | 0.366±0.028 | 0.393±0.033 | 0.378±0.019 | 0.370±0.020 | 0.006 | [-0.011, 0.023] | 0.711 |
| ATR_right | 0.365±0.028 | 0.392±0.040 | 0.377±0.022 | 0.369±0.029 | 0.006 | [-0.011, 0.023] | 0.718 |
| CA | 0.447±0.041 | 0.484±0.048 | 0.457±0.041 | 0.465±0.044 | -0.003 | [-0.021, 0.015] | 0.399 |
| CC_1 | 0.455±0.039 | 0.461±0.052 | 0.463±0.032 | 0.445±0.047 | 0.018 | [0.000, 0.036] | 0.948 |
| CC_2 | 0.389±0.031 | 0.405±0.037 | 0.398±0.027 | 0.388±0.027 | 0.010 | [-0.007, 0.027] | 0.834 |
| CC_3 | 0.412±0.047 | 0.430±0.045 | 0.422±0.042 | 0.411±0.037 | 0.012 | [-0.005, 0.030] | 0.871 |
| CC_4 | 0.453±0.046 | 0.469±0.035 | 0.458±0.045 | 0.459±0.032 | 0.008 | [-0.010, 0.025] | 0.757 |
| CC_5 | 0.428±0.036 | 0.440±0.037 | 0.433±0.035 | 0.431±0.034 | 0.008 | [-0.009, 0.025] | 0.781 |
| CC_6 | 0.402±0.027 | 0.428±0.036 | 0.410±0.027 | 0.414±0.031 | 0.001 | [-0.016, 0.019] | 0.548 |
| CC_7 | 0.392±0.034 | 0.440±0.038 | 0.404±0.036 | 0.417±0.028 | -0.009 | [-0.027, 0.009] | 0.215 |
| CG_left | 0.344±0.024 | 0.375±0.036 | 0.354±0.020 | 0.356±0.029 | 0.001 | [-0.016, 0.018] | 0.544 |
| CG_right | 0.360±0.025 | 0.384±0.033 | 0.368±0.022 | 0.368±0.024 | 0.003 | [-0.014, 0.020] | 0.630 |
| CST_left | 0.548±0.044 | 0.567±0.039 | 0.557±0.040 | 0.550±0.034 | 0.010 | [-0.007, 0.027] | 0.824 |
| CST_right | 0.510±0.041 | 0.531±0.037 | 0.520±0.038 | 0.513±0.031 | 0.009 | [-0.008, 0.026] | 0.801 |
| FPT_left | 0.471±0.036 | 0.497±0.036 | 0.483±0.031 | 0.476±0.025 | 0.007 | [-0.010, 0.024] | 0.758 |
| FPT_right | 0.464±0.036 | 0.482±0.037 | 0.474±0.032 | 0.463±0.025 | 0.011 | [-0.006, 0.029] | 0.854 |
| FX_left | 0.484±0.037 | 0.524±0.051 | 0.495±0.035 | 0.504±0.046 | -0.004 | [-0.022, 0.014] | 0.358 |
| FX_right | 0.500±0.041 | 0.526±0.057 | 0.509±0.038 | 0.509±0.052 | 0.003 | [-0.015, 0.020] | 0.595 |
| ICP_left | 0.402±0.029 | 0.422±0.023 | 0.408±0.025 | 0.411±0.024 | 0.004 | [-0.014, 0.022] | 0.637 |
| ICP_right | 0.370±0.027 | 0.376±0.025 | 0.375±0.024 | 0.366±0.024 | 0.012 | [-0.006, 0.029] | 0.858 |
| IFO_left | 0.395±0.025 | 0.423±0.034 | 0.406±0.022 | 0.402±0.018 | 0.004 | [-0.013, 0.021] | 0.640 |
| IFO_right | 0.387±0.024 | 0.421±0.033 | 0.398±0.024 | 0.401±0.022 | -0.001 | [-0.017, 0.017] | 0.481 |
| ILF_left | 0.403±0.028 | 0.439±0.036 | 0.415±0.028 | 0.418±0.021 | -0.002 | [-0.019, 0.015] | 0.424 |
| ILF_right | 0.402±0.034 | 0.441±0.040 | 0.415±0.032 | 0.417±0.026 | -0.001 | [-0.018, 0.016] | 0.455 |
| MCP | 0.454±0.035 | 0.465±0.032 | 0.459±0.032 | 0.454±0.029 | 0.010 | [-0.007, 0.027] | 0.822 |
| MLF_left | 0.381±0.026 | 0.403±0.031 | 0.389±0.023 | 0.388±0.025 | 0.004 | [-0.013, 0.021] | 0.657 |
| MLF_right | 0.380±0.026 | 0.403±0.036 | 0.387±0.025 | 0.388±0.027 | 0.002 | [-0.015, 0.019] | 0.586 |
| OR_left | 0.400±0.032 | 0.434±0.042 | 0.412±0.031 | 0.412±0.024 | 0.000 | [-0.017, 0.017] | 0.506 |
| OR_right | 0.396±0.028 | 0.442±0.036 | 0.407±0.030 | 0.420±0.026 | -0.008 | [-0.026, 0.010] | 0.233 |
| POPT_left | 0.456±0.028 | 0.474±0.033 | 0.464±0.023 | 0.458±0.027 | 0.008 | [-0.009, 0.025] | 0.778 |
| POPT_right | 0.451±0.030 | 0.470±0.037 | 0.460±0.027 | 0.453±0.030 | 0.008 | [-0.010, 0.024] | 0.763 |
| SCP_left | 0.448±0.033 | 0.473±0.028 | 0.458±0.028 | 0.455±0.025 | 0.006 | [-0.012, 0.022] | 0.708 |
| SCP_right | 0.425±0.030 | 0.445±0.029 | 0.433±0.026 | 0.428±0.024 | 0.007 | [-0.010, 0.024] | 0.746 |
| SLF_III_left | 0.375±0.027 | 0.410±0.049 | 0.388±0.023 | 0.387±0.035 | 0.001 | [-0.016, 0.019] | 0.550 |
| SLF_III_right | 0.387±0.032 | 0.410±0.046 | 0.398±0.026 | 0.390±0.038 | 0.008 | [-0.009, 0.026] | 0.775 |
| SLF_II_left | 0.354±0.031 | 0.366±0.032 | 0.362±0.026 | 0.352±0.026 | 0.011 | [-0.007, 0.028] | 0.842 |
| SLF_II_right | 0.342±0.032 | 0.369±0.039 | 0.352±0.026 | 0.350±0.028 | 0.004 | [-0.014, 0.021] | 0.632 |
| SLF_I_left | 0.318±0.025 | 0.336±0.029 | 0.325±0.021 | 0.323±0.025 | 0.006 | [-0.011, 0.023] | 0.721 |
| SLF_I_right | 0.332±0.029 | 0.360±0.028 | 0.341±0.027 | 0.343±0.022 | 0.001 | [-0.016, 0.019] | 0.557 |
| STR_left | 0.486±0.038 | 0.506±0.042 | 0.494±0.036 | 0.490±0.036 | 0.008 | [-0.009, 0.025] | 0.773 |
| STR_right | 0.467±0.036 | 0.471±0.042 | 0.473±0.033 | 0.461±0.037 | 0.015 | [-0.002, 0.033] | 0.918 |
| UF_left | 0.429±0.032 | 0.461±0.031 | 0.440±0.028 | 0.440±0.021 | 0.002 | [-0.015, 0.019] | 0.591 |
| UF_right | 0.420±0.031 | 0.449±0.035 | 0.430±0.030 | 0.430±0.026 | 0.002 | [-0.015, 0.019] | 0.588 |

[1] Units for raw and adjusted means, difference, and 95% HDI are arbitrary.

Data are presented as mean ± SD. Raw values are unadjusted sex-wise means. Adjusted values are sex-wise mean residual values after adjusting for age, education, intracranial volume, APOE e4 carriership, and family history of Alzheimer's disease. Bayesian model outputs include the mean difference and 95% highest density interval estimates as well as P+. P+ values approaching 1 indicate stronger evidence of female > male while P+ values approaching 0 indicate stronger evidence of male > female. The hierarchical model includes all tracts in a singular model and leverages partial pooling of the data, therefore the mean difference will not always be exactly equal to the difference in covariate adjusted means.

### Supplemental table 9: Fractional anisotropy differences between pre-menopausal women and age-matched men

| | Raw means[1] | | Covariate adjusted means[1] | | Hierarchical Bayesian model[1] | | |
|---|---|---|---|---|---|---|---|
| Tract | Females | Males | Females | Males | Mean Difference | 95% HDI | P+ |
| AF_left | 0.388±0.014 | 0.393±0.014 | 0.387±0.012 | 0.393±0.013 | -0.003 | [-0.010, 0.003] | 0.205 |
| AF_right | 0.388±0.014 | 0.389±0.014 | 0.387±0.013 | 0.390±0.011 | -0.002 | [-0.008, 0.004] | 0.326 |
| ATR_left | 0.363±0.010 | 0.369±0.013 | 0.363±0.010 | 0.370±0.013 | -0.004 | [-0.011, 0.002] | 0.153 |
| ATR_right | 0.343±0.009 | 0.346±0.014 | 0.343±0.009 | 0.347±0.014 | -0.003 | [-0.010, 0.003] | 0.224 |
| CA | 0.397±0.052 | 0.406±0.046 | 0.400±0.048 | 0.402±0.047 | -0.008 | [-0.015, -0.001] | 0.035 |
| CC_1 | 0.437±0.019 | 0.442±0.021 | 0.436±0.018 | 0.442±0.019 | -0.003 | [-0.009, 0.003] | 0.228 |
| CC_2 | 0.413±0.014 | 0.411±0.014 | 0.412±0.013 | 0.412±0.012 | -0.001 | [-0.007, 0.005] | 0.393 |
| CC_3 | 0.406±0.016 | 0.402±0.013 | 0.405±0.014 | 0.403±0.012 | 0.000 | [-0.007, 0.006] | 0.448 |
| CC_4 | 0.451±0.017 | 0.446±0.015 | 0.450±0.017 | 0.447±0.014 | 0.000 | [-0.007, 0.006] | 0.461 |
| CC_5 | 0.454±0.018 | 0.449±0.018 | 0.454±0.018 | 0.450±0.017 | 0.000 | [-0.006, 0.007] | 0.519 |
| CC_6 | 0.480±0.016 | 0.480±0.018 | 0.480±0.015 | 0.480±0.016 | -0.002 | [-0.008, 0.005] | 0.343 |
| CC_7 | 0.496±0.026 | 0.496±0.024 | 0.496±0.026 | 0.496±0.024 | -0.002 | [-0.009, 0.004] | 0.307 |
| CG_left | 0.414±0.018 | 0.423±0.017 | 0.413±0.016 | 0.424±0.015 | -0.004 | [-0.011, 0.002] | 0.137 |
| CG_right | 0.404±0.014 | 0.414±0.018 | 0.403±0.013 | 0.415±0.017 | -0.004 | [-0.011, 0.002] | 0.133 |
| CST_left | 0.491±0.014 | 0.496±0.013 | 0.492±0.014 | 0.496±0.011 | -0.004 | [-0.011, 0.002] | 0.158 |
| CST_right | 0.485±0.012 | 0.490±0.012 | 0.485±0.011 | 0.489±0.011 | -0.004 | [-0.010, 0.003] | 0.174 |
| FPT_left | 0.457±0.014 | 0.465±0.014 | 0.458±0.013 | 0.464±0.013 | -0.005 | [-0.011, 0.002] | 0.114 |
| FPT_right | 0.448±0.012 | 0.454±0.017 | 0.448±0.011 | 0.454±0.016 | -0.004 | [-0.011, 0.002] | 0.143 |
| FX_left | 0.397±0.037 | 0.386±0.043 | 0.394±0.035 | 0.390±0.036 | 0.005 | [-0.002, 0.012] | 0.865 |
| FX_right | 0.399±0.040 | 0.375±0.044 | 0.397±0.039 | 0.379±0.041 | 0.007 | [0.000, 0.015] | 0.944 |
| ICP_left | 0.415±0.013 | 0.426±0.021 | 0.415±0.012 | 0.425±0.020 | -0.006 | [-0.012, 0.001] | 0.071 |
| ICP_right | 0.422±0.019 | 0.425±0.020 | 0.422±0.017 | 0.424±0.020 | -0.004 | [-0.010, 0.002] | 0.158 |
| IFO_left | 0.420±0.016 | 0.424±0.013 | 0.420±0.015 | 0.423±0.013 | -0.003 | [-0.010, 0.003] | 0.191 |
| IFO_right | 0.427±0.017 | 0.430±0.014 | 0.427±0.016 | 0.430±0.013 | -0.003 | [-0.009, 0.003] | 0.215 |
| ILF_left | 0.412±0.019 | 0.415±0.015 | 0.412±0.018 | 0.415±0.015 | -0.004 | [-0.010, 0.002] | 0.169 |
| ILF_right | 0.411±0.017 | 0.411±0.015 | 0.411±0.016 | 0.411±0.015 | -0.002 | [-0.009, 0.004] | 0.269 |
| MCP | 0.479±0.011 | 0.485±0.016 | 0.479±0.010 | 0.485±0.015 | -0.004 | [-0.010, 0.003] | 0.184 |
| MLF_left | 0.386±0.016 | 0.384±0.014 | 0.386±0.014 | 0.385±0.014 | -0.001 | [-0.007, 0.005] | 0.394 |
| MLF_right | 0.376±0.014 | 0.378±0.017 | 0.376±0.012 | 0.378±0.015 | -0.002 | [-0.009, 0.004] | 0.276 |
| OR_left | 0.421±0.016 | 0.422±0.017 | 0.421±0.016 | 0.422±0.016 | -0.002 | [-0.009, 0.004] | 0.277 |
| OR_right | 0.431±0.015 | 0.431±0.016 | 0.431±0.015 | 0.432±0.015 | -0.002 | [-0.008, 0.005] | 0.339 |
| POPT_left | 0.452±0.012 | 0.456±0.013 | 0.452±0.012 | 0.455±0.012 | -0.003 | [-0.010, 0.003] | 0.202 |
| POPT_right | 0.458±0.013 | 0.463±0.014 | 0.458±0.012 | 0.463±0.012 | -0.004 | [-0.010, 0.003] | 0.185 |
| SCP_left | 0.428±0.015 | 0.435±0.012 | 0.428±0.013 | 0.434±0.013 | -0.005 | [-0.012, 0.001] | 0.088 |
| SCP_right | 0.406±0.015 | 0.415±0.015 | 0.407±0.012 | 0.414±0.013 | -0.006 | [-0.012, 0.001] | 0.075 |
| SLF_III_left | 0.396±0.016 | 0.402±0.013 | 0.396±0.014 | 0.403±0.012 | -0.003 | [-0.010, 0.003] | 0.199 |
| SLF_III_right | 0.387±0.013 | 0.391±0.014 | 0.386±0.012 | 0.391±0.013 | -0.003 | [-0.009, 0.003] | 0.232 |
| SLF_II_left | 0.355±0.018 | 0.346±0.017 | 0.354±0.017 | 0.347±0.017 | 0.001 | [-0.005, 0.008] | 0.609 |
| SLF_II_right | 0.359±0.018 | 0.356±0.013 | 0.359±0.015 | 0.357±0.013 | 0.000 | [-0.007, 0.006] | 0.468 |
| SLF_I_left | 0.380±0.016 | 0.377±0.016 | 0.379±0.015 | 0.378±0.016 | -0.001 | [-0.007, 0.006] | 0.409 |
| SLF_I_right | 0.376±0.017 | 0.373±0.013 | 0.376±0.016 | 0.374±0.012 | -0.001 | [-0.007, 0.006] | 0.440 |
| STR_left | 0.397±0.017 | 0.400±0.014 | 0.397±0.017 | 0.400±0.013 | -0.003 | [-0.009, 0.003] | 0.227 |
| STR_right | 0.400±0.015 | 0.401±0.015 | 0.400±0.015 | 0.401±0.015 | -0.002 | [-0.009, 0.004] | 0.280 |
| UF_left | 0.363±0.015 | 0.377±0.014 | 0.363±0.014 | 0.376±0.014 | -0.007 | [-0.014, 0.000] | 0.049 |
| UF_right | 0.354±0.014 | 0.360±0.014 | 0.354±0.013 | 0.360±0.013 | -0.005 | [-0.011, 0.002] | 0.130 |

[1] Units for raw and adjusted means, difference, and 95% HDI are arbitrary.

Data are presented as mean ± SD. Raw values are unadjusted sex-wise means. Adjusted values are sex-wise mean residual values after adjusting for age, education, APOE e4 carriership, and family history of Alzheimer's disease. Bayesian model outputs include the mean difference and 95% highest density interval estimates as well as P+. P+ indicates the proportion of the Bayesian samples (N = 80000) where the difference between women and men was > 0. P+ values approaching 1 indicate stronger evidence of female > male while P+ values approaching 0 indicate stronger evidence of male > female. The hierarchical model includes all tracts in a singular model and leverages partial pooling of the data, therefore the mean difference will not always be exactly equal to the difference in covariate adjusted means.

### Supplemental table 10: Mean diffusivity differences between pre-menopausal women and age-matched men

| | Raw means[1] | | Covariate adjusted means[1] | | Hierarchical Bayesian model[1] | | |
|---|---|---|---|---|---|---|---|
| Tract | Females | Males | Females | Males | Mean Difference | 95% HDI | P+ |
| AF_left | 7.113±0.219 | 7.055±0.182 | 7.112±0.215 | 7.058±0.172 | 0.062 | [-0.075, 0.199] | 0.768 |
| AF_right | 7.195±0.183 | 7.109±0.214 | 7.194±0.176 | 7.111±0.203 | 0.063 | [-0.071, 0.202] | 0.772 |
| ATR_left | 7.530±0.228 | 7.511±0.213 | 7.529±0.227 | 7.512±0.208 | 0.025 | [-0.108, 0.167] | 0.617 |
| ATR_right | 7.809±0.194 | 7.828±0.273 | 7.812±0.198 | 7.821±0.246 | -0.002 | [-0.137, 0.135] | 0.487 |
| CA | 8.101±0.369 | 7.932±0.293 | 8.081±0.340 | 7.971±0.273 | 0.018 | [-0.125, 0.159] | 0.573 |
| CC_1 | 8.154±0.371 | 8.046±0.325 | 8.153±0.361 | 8.047±0.331 | 0.002 | [-0.136, 0.139] | 0.503 |
| CC_2 | 8.449±0.363 | 8.463±0.267 | 8.451±0.350 | 8.458±0.271 | -0.046 | [-0.180, 0.093] | 0.289 |
| CC_3 | 8.697±0.469 | 8.839±0.383 | 8.710±0.452 | 8.816±0.386 | -0.090 | [-0.225, 0.048] | 0.146 |
| CC_4 | 8.369±0.428 | 8.424±0.342 | 8.370±0.419 | 8.423±0.347 | -0.051 | [-0.187, 0.086] | 0.275 |
| CC_5 | 8.728±0.456 | 8.800±0.396 | 8.727±0.456 | 8.802±0.367 | -0.079 | [-0.217, 0.058] | 0.177 |
| CC_6 | 7.851±0.312 | 7.847±0.301 | 7.848±0.304 | 7.853±0.281 | -0.002 | [-0.141, 0.131] | 0.491 |
| CC_7 | 9.080±0.681 | 9.400±0.869 | 9.084±0.663 | 9.393±0.882 | -0.160 | [-0.307, -0.023] | 0.034 |
| CG_left | 7.265±0.202 | 7.183±0.211 | 7.260±0.194 | 7.192±0.198 | 0.056 | [-0.077, 0.195] | 0.747 |
| CG_right | 7.250±0.193 | 7.159±0.263 | 7.249±0.184 | 7.161±0.244 | 0.060 | [-0.076, 0.197] | 0.763 |
| CST_left | 6.970±0.184 | 6.920±0.124 | 6.966±0.174 | 6.928±0.124 | 0.069 | [-0.068, 0.206] | 0.794 |
| CST_right | 7.112±0.129 | 7.022±0.172 | 7.108±0.121 | 7.030±0.161 | 0.069 | [-0.068, 0.204] | 0.793 |
| FPT_left | 7.159±0.156 | 7.096±0.133 | 7.155±0.150 | 7.103±0.129 | 0.060 | [-0.078, 0.196] | 0.760 |
| FPT_right | 7.362±0.150 | 7.275±0.168 | 7.358±0.143 | 7.281±0.166 | 0.051 | [-0.084, 0.188] | 0.727 |
| FX_left | 15.597±1.580 | 16.327±1.783 | 15.679±1.542 | 16.173±1.574 | -0.676 | [-0.855, -0.499] | 0.000 |
| FX_right | 15.218±1.504 | 16.309±1.865 | 15.286±1.500 | 16.183±1.731 | -0.738 | [-0.914, -0.560] | 0.000 |
| ICP_left | 7.167±0.177 | 7.011±0.193 | 7.156±0.163 | 7.031±0.197 | 0.079 | [-0.059, 0.218] | 0.821 |
| ICP_right | 6.932±0.235 | 6.802±0.231 | 6.919±0.216 | 6.828±0.217 | 0.088 | [-0.050, 0.225] | 0.850 |
| IFO_left | 7.826±0.328 | 7.794±0.258 | 7.818±0.318 | 7.808±0.241 | 0.005 | [-0.135, 0.139] | 0.524 |
| IFO_right | 7.872±0.273 | 7.897±0.236 | 7.873±0.255 | 7.896±0.218 | -0.010 | [-0.146, 0.124] | 0.453 |
| ILF_left | 7.800±0.387 | 7.821±0.287 | 7.792±0.370 | 7.836±0.257 | -0.006 | [-0.143, 0.130] | 0.473 |
| ILF_right | 7.705±0.285 | 7.795±0.202 | 7.706±0.255 | 7.793±0.197 | -0.012 | [-0.148, 0.125] | 0.445 |
| MCP | 6.925±0.129 | 6.864±0.163 | 6.925±0.132 | 6.862±0.144 | 0.076 | [-0.059, 0.215] | 0.813 |
| MLF_left | 7.572±0.263 | 7.541±0.218 | 7.569±0.252 | 7.547±0.215 | 0.024 | [-0.112, 0.159] | 0.611 |
| MLF_right | 7.413±0.236 | 7.300±0.253 | 7.411±0.217 | 7.303±0.220 | 0.054 | [-0.084, 0.190] | 0.734 |
| OR_left | 8.135±0.448 | 8.237±0.465 | 8.133±0.444 | 8.239±0.441 | -0.047 | [-0.183, 0.091] | 0.291 |
| OR_right | 7.912±0.293 | 8.009±0.347 | 7.915±0.273 | 8.004±0.346 | -0.028 | [-0.167, 0.106] | 0.375 |
| POPT_left | 7.626±0.184 | 7.631±0.187 | 7.623±0.178 | 7.637±0.164 | 0.011 | [-0.124, 0.149] | 0.555 |
| POPT_right | 7.610±0.177 | 7.558±0.226 | 7.606±0.169 | 7.565±0.200 | 0.026 | [-0.110, 0.162] | 0.620 |
| SCP_left | 8.065±0.222 | 8.109±0.269 | 8.058±0.203 | 8.122±0.264 | -0.029 | [-0.164, 0.110] | 0.367 |
| SCP_right | 8.337±0.229 | 8.303±0.284 | 8.325±0.196 | 8.326±0.279 | -0.030 | [-0.167, 0.112] | 0.361 |
| SLF_III_left | 7.070±0.242 | 7.014±0.193 | 7.069±0.237 | 7.018±0.176 | 0.064 | [-0.074, 0.201] | 0.778 |
| SLF_III_right | 7.324±0.195 | 7.261±0.233 | 7.321±0.189 | 7.267±0.221 | 0.048 | [-0.086, 0.186] | 0.716 |
| SLF_II_left | 7.062±0.203 | 7.063±0.209 | 7.065±0.194 | 7.058±0.202 | 0.053 | [-0.084, 0.191] | 0.734 |
| SLF_II_right | 7.210±0.183 | 7.154±0.241 | 7.211±0.169 | 7.153±0.230 | 0.055 | [-0.079, 0.193] | 0.742 |
| SLF_I_left | 7.229±0.201 | 7.228±0.249 | 7.230±0.186 | 7.226±0.234 | 0.041 | [-0.095, 0.179] | 0.688 |
| SLF_I_right | 7.365±0.213 | 7.319±0.265 | 7.364±0.195 | 7.320±0.255 | 0.042 | [-0.094, 0.180] | 0.688 |
| STR_left | 6.994±0.178 | 6.997±0.178 | 6.989±0.173 | 7.006±0.145 | 0.055 | [-0.084, 0.191] | 0.743 |
| STR_right | 7.015±0.147 | 6.938±0.195 | 7.014±0.142 | 6.939±0.179 | 0.074 | [-0.065, 0.210] | 0.804 |
| UF_left | 7.678±0.276 | 7.474±0.210 | 7.666±0.272 | 7.497±0.176 | 0.054 | [-0.087, 0.192] | 0.732 |
| UF_right | 7.911±0.219 | 7.798±0.170 | 7.910±0.216 | 7.801±0.169 | 0.020 | [-0.119, 0.156] | 0.589 |

[1] Units for raw and adjusted means, difference, and 95% HDI are $10^{-4}$ $mm^2/s$.

Data are presented as mean ± SD. Raw values are unadjusted sex-wise means. Adjusted values are sex-wise mean residual values after adjusting for age, education, APOE e4 carriership, and family history of Alzheimer's disease. Bayesian model outputs include the mean difference and 95% highest density interval estimates as well as P+. P+ indicates the proportion of the Bayesian samples (N = 80000) where the difference between women and men was > 0. P+ values approaching 1 indicate stronger evidence of female > male while P+ values approaching 0 indicate stronger evidence of male > female. The hierarchical model includes all tracts in a singular model and leverages partial pooling of the data, therefore the mean difference will not always be exactly equal to the difference in covariate adjusted means.

### Supplemental table 11: Fiber density differences between peri-menopausal women and age-matched men

| | Raw means[1] | | Covariate adjusted means[1] | | Hierarchical Bayesian model[1] | | |
|---|---|---|---|---|---|---|---|
| **Tract** | **Females** | **Males** | **Females** | **Males** | **Mean Difference** | **95% HDI** | **P+** |
| AF_left | 0.368±0.016 | 0.359±0.013 | 0.367±0.015 | 0.361±0.012 | 0.008 | [0.001, 0.015] | 0.967 |
| AF_right | 0.393±0.020 | 0.382±0.016 | 0.392±0.020 | 0.384±0.014 | 0.008 | [0.001, 0.016] | 0.970 |
| ATR_left | 0.366±0.017 | 0.361±0.016 | 0.365±0.015 | 0.362±0.012 | 0.008 | [0.001, 0.015] | 0.962 |
| ATR_right | 0.367±0.015 | 0.360±0.015 | 0.366±0.014 | 0.362±0.014 | 0.008 | [0.001, 0.015] | 0.958 |
| CA | 0.441±0.028 | 0.439±0.022 | 0.440±0.026 | 0.442±0.023 | 0.007 | [0.000, 0.015] | 0.942 |
| CC_1 | 0.455±0.020 | 0.436±0.017 | 0.453±0.019 | 0.438±0.016 | 0.010 | [0.002, 0.017] | 0.982 |
| CC_2 | 0.384±0.016 | 0.372±0.014 | 0.384±0.015 | 0.374±0.012 | 0.009 | [0.002, 0.016] | 0.974 |
| CC_3 | 0.400±0.021 | 0.386±0.020 | 0.399±0.018 | 0.389±0.017 | 0.009 | [0.002, 0.017] | 0.980 |
| CC_4 | 0.458±0.023 | 0.439±0.019 | 0.456±0.022 | 0.444±0.015 | 0.010 | [0.002, 0.017] | 0.984 |
| CC_5 | 0.433±0.022 | 0.414±0.019 | 0.431±0.021 | 0.419±0.014 | 0.010 | [0.003, 0.017] | 0.986 |
| CC_6 | 0.411±0.019 | 0.393±0.014 | 0.409±0.018 | 0.396±0.013 | 0.010 | [0.002, 0.017] | 0.984 |
| CC_7 | 0.385±0.018 | 0.375±0.012 | 0.384±0.017 | 0.378±0.011 | 0.008 | [0.001, 0.016] | 0.970 |
| CG_left | 0.351±0.014 | 0.348±0.013 | 0.351±0.014 | 0.347±0.012 | 0.007 | [0.000, 0.014] | 0.936 |
| CG_right | 0.365±0.018 | 0.357±0.012 | 0.365±0.017 | 0.358±0.012 | 0.008 | [0.001, 0.015] | 0.961 |
| CST_left | 0.555±0.021 | 0.548±0.019 | 0.554±0.021 | 0.549±0.017 | 0.007 | [-0.001, 0.014] | 0.926 |
| CST_right | 0.520±0.021 | 0.513±0.019 | 0.519±0.021 | 0.515±0.018 | 0.007 | [0.000, 0.015] | 0.943 |
| FPT_left | 0.478±0.017 | 0.474±0.018 | 0.478±0.016 | 0.473±0.016 | 0.007 | [-0.001, 0.014] | 0.925 |
| FPT_right | 0.467±0.015 | 0.462±0.020 | 0.467±0.015 | 0.463±0.019 | 0.007 | [0.000, 0.014] | 0.935 |
| FX_left | 0.519±0.068 | 0.499±0.070 | 0.515±0.058 | 0.506±0.059 | 0.013 | [0.004, 0.021] | 0.993 |
| FX_right | 0.536±0.066 | 0.508±0.072 | 0.531±0.055 | 0.516±0.064 | 0.014 | [0.005, 0.022] | 0.996 |
| ICP_left | 0.418±0.019 | 0.412±0.016 | 0.417±0.019 | 0.413±0.016 | 0.007 | [0.000, 0.014] | 0.944 |
| ICP_right | 0.382±0.018 | 0.372±0.017 | 0.381±0.018 | 0.372±0.016 | 0.008 | [0.001, 0.015] | 0.963 |
| IFO_left | 0.388±0.016 | 0.377±0.013 | 0.387±0.015 | 0.379±0.011 | 0.009 | [0.001, 0.016] | 0.972 |
| IFO_right | 0.384±0.018 | 0.373±0.011 | 0.383±0.016 | 0.375±0.009 | 0.009 | [0.001, 0.016] | 0.973 |
| ILF_left | 0.389±0.017 | 0.381±0.011 | 0.388±0.016 | 0.383±0.011 | 0.008 | [0.001, 0.015] | 0.958 |
| ILF_right | 0.391±0.021 | 0.384±0.014 | 0.390±0.020 | 0.386±0.014 | 0.008 | [0.001, 0.015] | 0.959 |
| MCP | 0.456±0.020 | 0.447±0.014 | 0.456±0.020 | 0.448±0.013 | 0.008 | [0.001, 0.015] | 0.960 |
| MLF_left | 0.377±0.019 | 0.364±0.011 | 0.376±0.019 | 0.367±0.011 | 0.009 | [0.002, 0.016] | 0.974 |
| MLF_right | 0.382±0.021 | 0.366±0.015 | 0.380±0.020 | 0.369±0.013 | 0.009 | [0.002, 0.017] | 0.982 |
| OR_left | 0.392±0.016 | 0.384±0.014 | 0.391±0.015 | 0.386±0.013 | 0.008 | [0.001, 0.015] | 0.967 |
| OR_right | 0.401±0.020 | 0.390±0.014 | 0.400±0.018 | 0.393±0.013 | 0.009 | [0.002, 0.016] | 0.974 |
| POPT_left | 0.463±0.021 | 0.451±0.014 | 0.462±0.021 | 0.453±0.014 | 0.008 | [0.001, 0.015] | 0.966 |
| POPT_right | 0.464±0.019 | 0.449±0.015 | 0.462±0.018 | 0.452±0.014 | 0.009 | [0.002, 0.016] | 0.973 |
| SCP_left | 0.463±0.015 | 0.467±0.018 | 0.463±0.015 | 0.467±0.017 | 0.005 | [-0.002, 0.013] | 0.874 |
| SCP_right | 0.439±0.014 | 0.440±0.017 | 0.439±0.014 | 0.439±0.017 | 0.006 | [-0.002, 0.013] | 0.900 |
| SLF_III_left | 0.370±0.017 | 0.362±0.016 | 0.369±0.016 | 0.364±0.014 | 0.008 | [0.001, 0.016] | 0.967 |
| SLF_III_right | 0.380±0.020 | 0.367±0.016 | 0.379±0.020 | 0.368±0.014 | 0.009 | [0.001, 0.016] | 0.973 |
| SLF_II_left | 0.338±0.017 | 0.327±0.014 | 0.337±0.017 | 0.329±0.014 | 0.008 | [0.001, 0.016] | 0.969 |
| SLF_II_right | 0.333±0.020 | 0.324±0.015 | 0.332±0.019 | 0.325±0.014 | 0.008 | [0.001, 0.015] | 0.961 |
| SLF_I_left | 0.310±0.016 | 0.308±0.014 | 0.310±0.016 | 0.308±0.013 | 0.007 | [-0.001, 0.014] | 0.927 |
| SLF_I_right | 0.330±0.019 | 0.324±0.010 | 0.330±0.018 | 0.325±0.010 | 0.008 | [0.000, 0.015] | 0.953 |
| STR_left | 0.495±0.023 | 0.486±0.018 | 0.494±0.024 | 0.488±0.017 | 0.007 | [0.000, 0.015] | 0.947 |
| STR_right | 0.471±0.023 | 0.456±0.019 | 0.470±0.023 | 0.459±0.017 | 0.009 | [0.002, 0.016] | 0.974 |
| UF_left | 0.427±0.018 | 0.418±0.014 | 0.426±0.017 | 0.419±0.013 | 0.008 | [0.001, 0.015] | 0.962 |
| UF_right | 0.418±0.018 | 0.406±0.012 | 0.417±0.017 | 0.407±0.012 | 0.009 | [0.002, 0.016] | 0.973 |

[1] Units for raw and adjusted means, difference, and 95% HDI are arbitrary.

Data are presented as mean ± SD. Raw values are unadjusted sex-wise means. Adjusted values are sex-wise mean residual values after adjusting for age, education, APOE e4 carriership, and family history of Alzheimer's disease. Bayesian model outputs include the mean difference and 95% highest density interval estimates as well as P+. P+ indicates the proportion of the Bayesian samples (N = 80000) where the difference between women and men was > 0. P+ values approaching 1 indicate stronger evidence of female > male while P+ values approaching 0 indicate stronger evidence of male > female. The hierarchical model includes all tracts in a singular model and leverages partial pooling of the data, therefore the mean difference will not always be exactly equal to the difference in covariate adjusted means.

### Supplemental table 12: Fiber cross-section differences between peri-menopausal women and age-matched men

| | Raw means[1] | | Covariate adjusted means[1] | | Hierarchical Bayesian model[1] | | |
|---|---|---|---|---|---|---|---|
| Tract | Females | Males | Females | Males | Mean Difference | 95% HDI | P+ |
| AF_left | 0.010±0.055 | 0.105±0.071 | 0.042±0.037 | 0.045±0.030 | -0.005 | [-0.020, 0.011] | 0.311 |
| AF_right | 0.009±0.054 | 0.104±0.069 | 0.041±0.033 | 0.044±0.036 | -0.005 | [-0.020, 0.011] | 0.311 |
| ATR_left | -0.013±0.067 | 0.081±0.081 | 0.022±0.046 | 0.016±0.042 | -0.004 | [-0.020, 0.011] | 0.319 |
| ATR_right | -0.009±0.064 | 0.088±0.091 | 0.027±0.046 | 0.021±0.050 | -0.005 | [-0.020, 0.011] | 0.311 |
| CA | -0.008±0.059 | 0.085±0.084 | 0.020±0.047 | 0.032±0.068 | -0.005 | [-0.020, 0.011] | 0.303 |
| CC_1 | -0.014±0.080 | 0.057±0.086 | 0.014±0.060 | 0.004±0.068 | -0.003 | [-0.019, 0.013] | 0.390 |
| CC_2 | -0.001±0.065 | 0.089±0.079 | 0.032±0.040 | 0.027±0.044 | -0.004 | [-0.020, 0.012] | 0.342 |
| CC_3 | 0.004±0.065 | 0.103±0.101 | 0.039±0.059 | 0.038±0.062 | -0.005 | [-0.021, 0.010] | 0.305 |
| CC_4 | -0.016±0.061 | 0.071±0.065 | 0.011±0.053 | 0.019±0.043 | -0.005 | [-0.020, 0.011] | 0.316 |
| CC_5 | -0.022±0.056 | 0.062±0.065 | 0.005±0.047 | 0.010±0.038 | -0.005 | [-0.020, 0.011] | 0.312 |
| CC_6 | -0.031±0.052 | 0.075±0.074 | 0.004±0.034 | 0.011±0.041 | -0.006 | [-0.022, 0.010] | 0.266 |
| CC_7 | 0.010±0.069 | 0.139±0.094 | 0.052±0.053 | 0.063±0.054 | -0.007 | [-0.023, 0.009] | 0.233 |
| CG_left | -0.029±0.059 | 0.066±0.074 | 0.003±0.035 | 0.007±0.046 | -0.005 | [-0.021, 0.010] | 0.301 |
| CG_right | -0.024±0.055 | 0.063±0.073 | 0.006±0.037 | 0.007±0.047 | -0.005 | [-0.020, 0.011] | 0.319 |
| CST_left | -0.026±0.061 | 0.045±0.055 | -0.003±0.049 | 0.002±0.044 | -0.004 | [-0.019, 0.012] | 0.342 |
| CST_right | -0.028±0.057 | 0.049±0.066 | -0.001±0.042 | 0.000±0.047 | -0.004 | [-0.020, 0.011] | 0.331 |
| FPT_left | -0.019±0.051 | 0.061±0.060 | 0.009±0.029 | 0.008±0.037 | -0.004 | [-0.020, 0.011] | 0.330 |
| FPT_right | -0.014±0.053 | 0.058±0.062 | 0.013±0.034 | 0.008±0.034 | -0.003 | [-0.019, 0.012] | 0.355 |
| FX_left | -0.097±0.106 | 0.055±0.131 | -0.047±0.083 | -0.038±0.079 | -0.009 | [-0.027, 0.009] | 0.217 |
| FX_right | -0.100±0.101 | 0.060±0.142 | -0.048±0.078 | -0.037±0.087 | -0.010 | [-0.028, 0.010] | 0.205 |
| ICP_left | -0.018±0.066 | 0.022±0.044 | -0.003±0.058 | -0.006±0.043 | -0.002 | [-0.018, 0.015] | 0.430 |
| ICP_right | -0.026±0.059 | 0.021±0.059 | -0.008±0.049 | -0.013±0.056 | -0.002 | [-0.019, 0.014] | 0.404 |
| IFO_left | 0.003±0.058 | 0.108±0.075 | 0.039±0.036 | 0.042±0.032 | -0.006 | [-0.021, 0.010] | 0.281 |
| IFO_right | 0.007±0.055 | 0.114±0.078 | 0.043±0.033 | 0.047±0.040 | -0.006 | [-0.021, 0.010] | 0.277 |
| ILF_left | 0.021±0.058 | 0.133±0.079 | 0.058±0.036 | 0.064±0.036 | -0.006 | [-0.022, 0.010] | 0.271 |
| ILF_right | 0.020±0.059 | 0.131±0.083 | 0.057±0.038 | 0.061±0.042 | -0.006 | [-0.022, 0.010] | 0.276 |
| MCP | -0.012±0.068 | 0.045±0.054 | 0.007±0.057 | 0.010±0.056 | -0.003 | [-0.019, 0.013] | 0.388 |
| MLF_left | -0.011±0.050 | 0.093±0.067 | 0.022±0.030 | 0.032±0.037 | -0.006 | [-0.021, 0.010] | 0.273 |
| MLF_right | -0.020±0.053 | 0.087±0.074 | 0.015±0.032 | 0.024±0.043 | -0.006 | [-0.021, 0.010] | 0.268 |
| OR_left | 0.007±0.063 | 0.119±0.093 | 0.045±0.051 | 0.050±0.048 | -0.006 | [-0.022, 0.009] | 0.258 |
| OR_right | -0.004±0.062 | 0.113±0.090 | 0.035±0.043 | 0.040±0.049 | -0.007 | [-0.022, 0.009] | 0.250 |
| POPT_left | -0.033±0.051 | 0.050±0.060 | -0.005±0.034 | -0.002±0.034 | -0.005 | [-0.021, 0.011] | 0.306 |
| POPT_right | -0.036±0.053 | 0.046±0.065 | -0.007±0.036 | -0.007±0.036 | -0.005 | [-0.020, 0.011] | 0.317 |
| SCP_left | -0.029±0.051 | 0.018±0.052 | -0.009±0.039 | -0.019±0.036 | -0.002 | [-0.019, 0.014] | 0.404 |
| SCP_right | -0.033±0.050 | 0.018±0.054 | -0.012±0.036 | -0.020±0.040 | -0.003 | [-0.019, 0.013] | 0.388 |
| SLF_III_left | 0.009±0.059 | 0.109±0.095 | 0.044±0.044 | 0.044±0.058 | -0.005 | [-0.021, 0.010] | 0.297 |
| SLF_III_right | 0.012±0.062 | 0.099±0.074 | 0.043±0.045 | 0.043±0.052 | -0.004 | [-0.020, 0.011] | 0.329 |
| SLF_II_left | 0.014±0.059 | 0.105±0.073 | 0.045±0.042 | 0.046±0.041 | -0.004 | [-0.020, 0.011] | 0.323 |
| SLF_II_right | 0.010±0.055 | 0.113±0.078 | 0.044±0.037 | 0.049±0.041 | -0.005 | [-0.021, 0.010] | 0.288 |
| SLF_I_left | 0.004±0.050 | 0.087±0.066 | 0.032±0.036 | 0.035±0.042 | -0.004 | [-0.020, 0.011] | 0.322 |
| SLF_I_right | -0.003±0.052 | 0.084±0.069 | 0.027±0.031 | 0.028±0.039 | -0.005 | [-0.020, 0.011] | 0.316 |
| STR_left | -0.042±0.059 | 0.043±0.068 | -0.017±0.050 | -0.002±0.053 | -0.006 | [-0.021, 0.011] | 0.290 |
| STR_right | -0.025±0.058 | 0.052±0.072 | 0.000±0.050 | 0.006±0.052 | -0.004 | [-0.020, 0.011] | 0.321 |
| UF_left | -0.007±0.059 | 0.086±0.063 | 0.023±0.039 | 0.029±0.033 | -0.005 | [-0.020, 0.011] | 0.314 |
| UF_right | 0.009±0.052 | 0.092±0.070 | 0.037±0.037 | 0.041±0.046 | -0.004 | [-0.020, 0.011] | 0.342 |

[1] Units for raw and adjusted means, difference, and 95% HDI are arbitrary.

Data are presented as mean ± SD. Raw values are unadjusted sex-wise means. Adjusted values are sex-wise mean residual values after adjusting for age, education, intracranial volume, APOE e4 carriership, and family history of Alzheimer's disease. Bayesian model outputs include the mean difference and 95% highest density interval estimates as well as P+. P+ values approaching 1 indicate stronger evidence of female > male while P+ values approaching 0 indicate stronger evidence of male > female. The hierarchical model includes all tracts in a singular model and leverages partial pooling of the data, therefore the mean difference will not always be exactly equal to the difference in covariate adjusted means.

### Supplemental table 13: FDC differences between peri-menopausal women and age-matched men

| | Raw means[1] | | Covariate adjusted means[1] | | Hierarchical Bayesian model[1] | | |
|---|---|---|---|---|---|---|---|
| Tract | Females | Males | Females | Males | Mean Difference | 95% HDI | P+ |
| AF_left | 0.375±0.033 | 0.403±0.036 | 0.387±0.025 | 0.382±0.022 | 0.004 | [-0.011, 0.018] | 0.663 |
| AF_right | 0.398±0.033 | 0.427±0.038 | 0.409±0.027 | 0.407±0.026 | 0.003 | [-0.012, 0.018] | 0.638 |
| ATR_left | 0.362±0.031 | 0.392±0.034 | 0.374±0.022 | 0.370±0.021 | 0.004 | [-0.011, 0.018] | 0.656 |
| ATR_right | 0.364±0.028 | 0.394±0.037 | 0.375±0.022 | 0.372±0.028 | 0.004 | [-0.011, 0.019] | 0.664 |
| CA | 0.438±0.046 | 0.480±0.057 | 0.451±0.040 | 0.456±0.050 | -0.001 | [-0.016, 0.014] | 0.463 |
| CC_1 | 0.451±0.046 | 0.460±0.050 | 0.458±0.040 | 0.446±0.044 | 0.008 | [-0.007, 0.024] | 0.804 |
| CC_2 | 0.385±0.033 | 0.407±0.036 | 0.395±0.026 | 0.390±0.027 | 0.005 | [-0.009, 0.020] | 0.723 |
| CC_3 | 0.402±0.036 | 0.429±0.044 | 0.412±0.031 | 0.411±0.037 | 0.004 | [-0.011, 0.018] | 0.652 |
| CC_4 | 0.454±0.044 | 0.474±0.039 | 0.462±0.041 | 0.460±0.034 | 0.005 | [-0.010, 0.019] | 0.689 |
| CC_5 | 0.426±0.041 | 0.442±0.037 | 0.433±0.038 | 0.429±0.029 | 0.005 | [-0.010, 0.020] | 0.723 |
| CC_6 | 0.400±0.035 | 0.427±0.034 | 0.410±0.030 | 0.409±0.027 | 0.003 | [-0.011, 0.018] | 0.642 |
| CC_7 | 0.388±0.038 | 0.431±0.040 | 0.402±0.030 | 0.403±0.023 | -0.001 | [-0.016, 0.014] | 0.469 |
| CG_left | 0.343±0.028 | 0.374±0.034 | 0.353±0.022 | 0.355±0.028 | 0.003 | [-0.012, 0.018] | 0.631 |
| CG_right | 0.358±0.028 | 0.384±0.032 | 0.367±0.023 | 0.367±0.024 | 0.004 | [-0.011, 0.019] | 0.676 |
| CST_left | 0.543±0.046 | 0.573±0.040 | 0.554±0.040 | 0.552±0.035 | 0.001 | [-0.014, 0.016] | 0.550 |
| CST_right | 0.510±0.040 | 0.538±0.039 | 0.520±0.035 | 0.519±0.036 | 0.001 | [-0.014, 0.016] | 0.561 |
| FPT_left | 0.469±0.033 | 0.502±0.037 | 0.482±0.026 | 0.479±0.029 | 0.002 | [-0.013, 0.016] | 0.568 |
| FPT_right | 0.460±0.031 | 0.489±0.038 | 0.471±0.025 | 0.468±0.030 | 0.003 | [-0.012, 0.018] | 0.622 |
| FX_left | 0.466±0.052 | 0.516±0.055 | 0.476±0.050 | 0.497±0.051 | -0.006 | [-0.022, 0.011] | 0.292 |
| FX_right | 0.481±0.054 | 0.527±0.055 | 0.492±0.051 | 0.506±0.051 | -0.004 | [-0.019, 0.012] | 0.361 |
| ICP_left | 0.412±0.039 | 0.423±0.021 | 0.417±0.036 | 0.413±0.022 | 0.008 | [-0.008, 0.023] | 0.793 |
| ICP_right | 0.373±0.030 | 0.380±0.026 | 0.379±0.027 | 0.370±0.025 | 0.009 | [-0.006, 0.025] | 0.833 |
| IFO_left | 0.388±0.030 | 0.418±0.034 | 0.400±0.022 | 0.397±0.016 | 0.003 | [-0.012, 0.018] | 0.626 |
| IFO_right | 0.385±0.031 | 0.414±0.033 | 0.396±0.024 | 0.393±0.018 | 0.003 | [-0.012, 0.017] | 0.625 |
| ILF_left | 0.397±0.030 | 0.435±0.037 | 0.410±0.024 | 0.410±0.020 | 0.001 | [-0.014, 0.016] | 0.539 |
| ILF_right | 0.397±0.036 | 0.436±0.043 | 0.411±0.029 | 0.409±0.026 | 0.001 | [-0.014, 0.016] | 0.533 |
| MCP | 0.456±0.046 | 0.472±0.030 | 0.463±0.041 | 0.458±0.031 | 0.006 | [-0.009, 0.021] | 0.735 |
| MLF_left | 0.375±0.033 | 0.402±0.030 | 0.385±0.028 | 0.384±0.022 | 0.004 | [-0.011, 0.019] | 0.663 |
| MLF_right | 0.376±0.034 | 0.402±0.035 | 0.385±0.029 | 0.385±0.026 | 0.004 | [-0.011, 0.018] | 0.653 |
| OR_left | 0.392±0.032 | 0.428±0.041 | 0.405±0.026 | 0.404±0.024 | 0.001 | [-0.014, 0.016] | 0.536 |
| OR_right | 0.397±0.035 | 0.433±0.037 | 0.410±0.028 | 0.408±0.021 | 0.001 | [-0.014, 0.016] | 0.536 |
| POPT_left | 0.450±0.037 | 0.475±0.033 | 0.460±0.031 | 0.456±0.025 | 0.003 | [-0.012, 0.018] | 0.640 |
| POPT_right | 0.449±0.035 | 0.471±0.034 | 0.458±0.030 | 0.455±0.027 | 0.003 | [-0.011, 0.018] | 0.648 |
| SCP_left | 0.450±0.033 | 0.476±0.028 | 0.460±0.027 | 0.456±0.025 | 0.003 | [-0.011, 0.018] | 0.642 |
| SCP_right | 0.425±0.030 | 0.449±0.029 | 0.435±0.024 | 0.430±0.024 | 0.004 | [-0.010, 0.019] | 0.685 |
| SLF_III_left | 0.376±0.034 | 0.408±0.047 | 0.389±0.028 | 0.385±0.034 | 0.003 | [-0.012, 0.018] | 0.622 |
| SLF_III_right | 0.388±0.037 | 0.409±0.042 | 0.397±0.032 | 0.392±0.034 | 0.005 | [-0.010, 0.020] | 0.707 |
| SLF_II_left | 0.347±0.033 | 0.368±0.032 | 0.356±0.026 | 0.351±0.027 | 0.006 | [-0.009, 0.021] | 0.745 |
| SLF_II_right | 0.340±0.032 | 0.368±0.038 | 0.350±0.028 | 0.350±0.030 | 0.004 | [-0.011, 0.019] | 0.663 |
| SLF_I_left | 0.315±0.026 | 0.341±0.030 | 0.323±0.024 | 0.325±0.024 | 0.005 | [-0.010, 0.020] | 0.701 |
| SLF_I_right | 0.333±0.028 | 0.358±0.027 | 0.342±0.023 | 0.342±0.021 | 0.005 | [-0.010, 0.019] | 0.695 |
| STR_left | 0.478±0.045 | 0.508±0.044 | 0.489±0.041 | 0.488±0.037 | 0.001 | [-0.014, 0.016] | 0.554 |
| STR_right | 0.461±0.045 | 0.480±0.044 | 0.468±0.041 | 0.466±0.040 | 0.004 | [-0.011, 0.019] | 0.673 |
| UF_left | 0.424±0.034 | 0.456±0.031 | 0.435±0.027 | 0.436±0.020 | 0.002 | [-0.013, 0.017] | 0.585 |
| UF_right | 0.421±0.033 | 0.445±0.036 | 0.431±0.028 | 0.428±0.027 | 0.004 | [-0.011, 0.019] | 0.670 |

[1] Units for raw and adjusted means, difference, and 95% HDI are arbitrary.

Data are presented as mean ± SD. Raw values are unadjusted sex-wise means. Adjusted values are sex-wise mean residual values after adjusting for age, education, intracranial volume, APOE e4 carriership, and family history of Alzheimer's disease. Bayesian model outputs include the mean difference and 95% highest density interval estimates as well as P+. P+ values approaching 1 indicate stronger evidence of female > male while P+ values approaching 0 indicate stronger evidence of male > female. The hierarchical model includes all tracts in a singular model and leverages partial pooling of the data, therefore the mean difference will not always be exactly equal to the difference in covariate adjusted means.

### Supplemental table 14: Fractional anisotropy differences between peri-menopausal women and age-matched men

| | Raw means[1] | | Covariate adjusted means[1] | | Hierarchical Bayesian model[1] | | |
|---|---|---|---|---|---|---|---|
| Tract | Females | Males | Females | Males | Mean Difference | 95% HDI | P+ |
| AF_left | 0.388±0.016 | 0.389±0.013 | 0.389±0.015 | 0.387±0.012 | 0.000 | [-0.006, 0.006] | 0.468 |
| AF_right | 0.388±0.016 | 0.387±0.012 | 0.389±0.015 | 0.385±0.011 | 0.000 | [-0.006, 0.006] | 0.476 |
| ATR_left | 0.363±0.012 | 0.370±0.011 | 0.363±0.011 | 0.368±0.012 | -0.003 | [-0.009, 0.003] | 0.245 |
| ATR_right | 0.345±0.011 | 0.348±0.013 | 0.345±0.011 | 0.347±0.013 | -0.001 | [-0.007, 0.005] | 0.387 |
| CA | 0.388±0.044 | 0.407±0.054 | 0.387±0.043 | 0.408±0.053 | -0.009 | [-0.017, -0.001] | 0.025 |
| CC_1 | 0.436±0.018 | 0.442±0.021 | 0.436±0.018 | 0.441±0.021 | -0.003 | [-0.009, 0.003] | 0.220 |
| CC_2 | 0.410±0.020 | 0.412±0.014 | 0.411±0.020 | 0.412±0.014 | -0.002 | [-0.008, 0.004] | 0.309 |
| CC_3 | 0.400±0.015 | 0.404±0.015 | 0.401±0.015 | 0.403±0.015 | -0.002 | [-0.008, 0.004] | 0.258 |
| CC_4 | 0.451±0.016 | 0.447±0.014 | 0.451±0.015 | 0.447±0.014 | 0.000 | [-0.007, 0.006] | 0.466 |
| CC_5 | 0.451±0.017 | 0.447±0.016 | 0.450±0.016 | 0.448±0.015 | -0.001 | [-0.008, 0.005] | 0.362 |
| CC_6 | 0.478±0.012 | 0.480±0.017 | 0.478±0.012 | 0.479±0.017 | -0.002 | [-0.008, 0.004] | 0.318 |
| CC_7 | 0.492±0.019 | 0.494±0.023 | 0.491±0.018 | 0.495±0.022 | -0.003 | [-0.010, 0.003] | 0.196 |
| CG_left | 0.409±0.018 | 0.421±0.016 | 0.411±0.017 | 0.418±0.014 | -0.004 | [-0.010, 0.003] | 0.187 |
| CG_right | 0.401±0.017 | 0.411±0.018 | 0.403±0.017 | 0.408±0.016 | -0.003 | [-0.009, 0.003] | 0.228 |
| CST_left | 0.496±0.018 | 0.497±0.013 | 0.498±0.016 | 0.494±0.011 | 0.000 | [-0.007, 0.006] | 0.481 |
| CST_right | 0.490±0.016 | 0.492±0.012 | 0.490±0.014 | 0.491±0.012 | -0.001 | [-0.007, 0.006] | 0.431 |
| FPT_left | 0.459±0.015 | 0.468±0.013 | 0.461±0.013 | 0.465±0.012 | -0.003 | [-0.009, 0.003] | 0.234 |
| FPT_right | 0.451±0.015 | 0.458±0.015 | 0.452±0.013 | 0.456±0.015 | -0.002 | [-0.008, 0.004] | 0.305 |
| FX_left | 0.383±0.054 | 0.384±0.046 | 0.381±0.048 | 0.388±0.039 | -0.006 | [-0.013, 0.001] | 0.091 |
| FX_right | 0.379±0.050 | 0.379±0.044 | 0.377±0.046 | 0.381±0.042 | -0.006 | [-0.012, 0.001] | 0.093 |
| ICP_left | 0.416±0.015 | 0.423±0.018 | 0.417±0.014 | 0.422±0.019 | -0.003 | [-0.009, 0.003] | 0.223 |
| ICP_right | 0.423±0.016 | 0.425±0.018 | 0.423±0.015 | 0.425±0.018 | -0.002 | [-0.008, 0.004] | 0.333 |
| IFO_left | 0.420±0.013 | 0.421±0.011 | 0.421±0.012 | 0.420±0.012 | -0.001 | [-0.007, 0.005] | 0.407 |
| IFO_right | 0.427±0.014 | 0.427±0.011 | 0.428±0.013 | 0.427±0.010 | 0.000 | [-0.006, 0.006] | 0.461 |
| ILF_left | 0.414±0.016 | 0.413±0.014 | 0.415±0.015 | 0.412±0.015 | 0.000 | [-0.006, 0.006] | 0.509 |
| ILF_right | 0.413±0.017 | 0.410±0.013 | 0.413±0.016 | 0.409±0.012 | 0.001 | [-0.005, 0.007] | 0.608 |
| MCP | 0.480±0.013 | 0.483±0.012 | 0.480±0.012 | 0.483±0.012 | -0.003 | [-0.009, 0.003] | 0.250 |
| MLF_left | 0.385±0.013 | 0.383±0.013 | 0.385±0.012 | 0.382±0.013 | 0.001 | [-0.005, 0.007] | 0.599 |
| MLF_right | 0.375±0.014 | 0.376±0.015 | 0.376±0.013 | 0.375±0.015 | 0.000 | [-0.006, 0.006] | 0.515 |
| OR_left | 0.419±0.017 | 0.421±0.016 | 0.419±0.016 | 0.421±0.016 | -0.001 | [-0.007, 0.005] | 0.418 |
| OR_right | 0.428±0.016 | 0.431±0.013 | 0.429±0.014 | 0.430±0.014 | -0.001 | [-0.007, 0.005] | 0.431 |
| POPT_left | 0.453±0.015 | 0.455±0.012 | 0.454±0.013 | 0.454±0.011 | -0.001 | [-0.007, 0.005] | 0.422 |
| POPT_right | 0.459±0.013 | 0.464±0.013 | 0.460±0.013 | 0.462±0.012 | -0.001 | [-0.007, 0.005] | 0.370 |
| SCP_left | 0.429±0.012 | 0.439±0.013 | 0.430±0.012 | 0.437±0.013 | -0.004 | [-0.010, 0.002] | 0.158 |
| SCP_right | 0.409±0.012 | 0.420±0.017 | 0.411±0.012 | 0.417±0.014 | -0.004 | [-0.010, 0.003] | 0.172 |
| SLF_III_left | 0.396±0.013 | 0.398±0.013 | 0.397±0.013 | 0.397±0.012 | -0.001 | [-0.007, 0.005] | 0.348 |
| SLF_III_right | 0.387±0.013 | 0.389±0.013 | 0.387±0.013 | 0.387±0.012 | -0.002 | [-0.008, 0.004] | 0.311 |
| SLF_II_left | 0.352±0.017 | 0.345±0.015 | 0.353±0.017 | 0.345±0.015 | 0.002 | [-0.004, 0.009] | 0.706 |
| SLF_II_right | 0.359±0.017 | 0.356±0.012 | 0.360±0.016 | 0.355±0.012 | 0.001 | [-0.005, 0.007] | 0.585 |
| SLF_I_left | 0.377±0.014 | 0.376±0.013 | 0.378±0.013 | 0.374±0.012 | 0.001 | [-0.006, 0.007] | 0.556 |
| SLF_I_right | 0.375±0.015 | 0.374±0.012 | 0.376±0.014 | 0.373±0.012 | 0.001 | [-0.006, 0.007] | 0.564 |
| STR_left | 0.396±0.018 | 0.399±0.015 | 0.397±0.017 | 0.398±0.014 | 0.000 | [-0.006, 0.006] | 0.465 |
| STR_right | 0.399±0.018 | 0.402±0.016 | 0.400±0.017 | 0.401±0.016 | 0.000 | [-0.006, 0.006] | 0.466 |
| UF_left | 0.365±0.012 | 0.373±0.016 | 0.366±0.012 | 0.371±0.016 | -0.003 | [-0.009, 0.003] | 0.218 |
| UF_right | 0.355±0.012 | 0.358±0.013 | 0.356±0.011 | 0.357±0.013 | -0.001 | [-0.007, 0.005] | 0.364 |

[1] Units for raw and adjusted means, difference, and 95% HDI are arbitrary.

Data are presented as mean ± SD. Raw values are unadjusted sex-wise means. Adjusted values are sex-wise mean residual values after adjusting for age, education, APOE e4 carriership, and family history of Alzheimer's disease. Bayesian model outputs include the mean difference and 95% highest density interval estimates as well as P+. P+ indicates the proportion of the Bayesian samples (N = 80000) where the difference between women and men was > 0. P+ values approaching 1 indicate stronger evidence of female > male while P+ values approaching 0 indicate stronger evidence of male > female. The hierarchical model includes all tracts in a singular model and leverages partial pooling of the data, therefore the mean difference will not always be exactly equal to the difference in covariate adjusted means.

### Supplemental table 15: Mean diffusivity differences between peri-menopausal women and age-matched men

| | Raw means[1] | | Covariate adjusted means[1] | | Hierarchical Bayesian model[1] | | |
|---|---|---|---|---|---|---|---|
| Tract | Females | Males | Females | Males | Mean Difference | 95% HDI | P+ |
| AF_left | 7.132±0.206 | 7.071±0.179 | 7.123±0.197 | 7.088±0.185 | 0.029 | [-0.073, 0.129] | 0.680 |
| AF_right | 7.172±0.201 | 7.116±0.197 | 7.167±0.199 | 7.125±0.193 | 0.029 | [-0.071, 0.130] | 0.683 |
| ATR_left | 7.552±0.221 | 7.510±0.202 | 7.548±0.208 | 7.516±0.210 | 0.029 | [-0.071, 0.130] | 0.678 |
| ATR_right | 7.818±0.231 | 7.827±0.254 | 7.824±0.226 | 7.816±0.232 | 0.027 | [-0.072, 0.128] | 0.670 |
| CA | 8.156±0.328 | 7.948±0.291 | 8.153±0.323 | 7.954±0.277 | 0.035 | [-0.067, 0.136] | 0.708 |
| CC_1 | 8.130±0.295 | 7.991±0.305 | 8.128±0.288 | 7.995±0.314 | 0.032 | [-0.069, 0.133] | 0.696 |
| CC_2 | 8.490±0.427 | 8.450±0.260 | 8.492±0.422 | 8.446±0.256 | 0.029 | [-0.071, 0.130] | 0.680 |
| CC_3 | 8.782±0.391 | 8.816±0.384 | 8.786±0.367 | 8.808±0.380 | 0.026 | [-0.074, 0.126] | 0.664 |
| CC_4 | 8.385±0.328 | 8.421±0.319 | 8.388±0.299 | 8.415±0.317 | 0.026 | [-0.074, 0.127] | 0.666 |
| CC_5 | 8.804±0.409 | 8.840±0.368 | 8.825±0.361 | 8.802±0.321 | 0.027 | [-0.071, 0.130] | 0.669 |
| CC_6 | 7.903±0.233 | 7.853±0.283 | 7.907±0.225 | 7.845±0.268 | 0.029 | [-0.071, 0.130] | 0.683 |
| CC_7 | 9.053±0.546 | 9.382±0.827 | 9.105±0.532 | 9.284±0.747 | 0.016 | [-0.090, 0.119] | 0.606 |
| CG_left | 7.264±0.200 | 7.158±0.219 | 7.250±0.197 | 7.183±0.208 | 0.031 | [-0.071, 0.131] | 0.688 |
| CG_right | 7.222±0.223 | 7.148±0.255 | 7.212±0.221 | 7.166±0.244 | 0.030 | [-0.071, 0.131] | 0.685 |
| CST_left | 6.931±0.159 | 6.934±0.118 | 6.922±0.150 | 6.950±0.123 | 0.027 | [-0.074, 0.127] | 0.668 |
| CST_right | 7.092±0.152 | 7.028±0.158 | 7.087±0.151 | 7.037±0.159 | 0.030 | [-0.069, 0.132] | 0.683 |
| FPT_left | 7.118±0.177 | 7.087±0.133 | 7.106±0.167 | 7.110±0.134 | 0.028 | [-0.073, 0.128] | 0.675 |
| FPT_right | 7.312±0.193 | 7.261±0.153 | 7.306±0.190 | 7.272±0.157 | 0.029 | [-0.071, 0.130] | 0.679 |
| FX_left | 16.218±2.083 | 16.426±1.960 | 16.313±1.813 | 16.249±1.653 | 0.026 | [-0.091, 0.143] | 0.646 |
| FX_right | 15.974±1.907 | 16.199±1.907 | 16.062±1.636 | 16.037±1.776 | 0.021 | [-0.093, 0.140] | 0.624 |
| ICP_left | 7.110±0.185 | 7.024±0.176 | 7.104±0.182 | 7.034±0.181 | 0.030 | [-0.070, 0.131] | 0.687 |
| ICP_right | 6.864±0.204 | 6.790±0.211 | 6.865±0.204 | 6.788±0.208 | 0.030 | [-0.071, 0.131] | 0.685 |
| IFO_left | 7.838±0.218 | 7.776±0.242 | 7.835±0.216 | 7.780±0.240 | 0.029 | [-0.069, 0.132] | 0.682 |
| IFO_right | 7.827±0.215 | 7.871±0.218 | 7.840±0.210 | 7.847±0.197 | 0.026 | [-0.075, 0.126] | 0.664 |
| ILF_left | 7.811±0.245 | 7.812±0.278 | 7.809±0.246 | 7.815±0.268 | 0.027 | [-0.072, 0.129] | 0.672 |
| ILF_right | 7.663±0.226 | 7.767±0.195 | 7.672±0.218 | 7.749±0.191 | 0.023 | [-0.077, 0.124] | 0.650 |
| MCP | 6.887±0.163 | 6.843±0.156 | 6.885±0.160 | 6.846±0.149 | 0.028 | [-0.073, 0.129] | 0.678 |
| MLF_left | 7.587±0.198 | 7.545±0.226 | 7.580±0.186 | 7.559±0.228 | 0.028 | [-0.072, 0.128] | 0.677 |
| MLF_right | 7.391±0.202 | 7.319±0.232 | 7.384±0.199 | 7.332±0.217 | 0.030 | [-0.071, 0.129] | 0.684 |
| OR_left | 8.085±0.266 | 8.232±0.448 | 8.109±0.271 | 8.188±0.392 | 0.023 | [-0.080, 0.122] | 0.647 |
| OR_right | 7.897±0.269 | 7.985±0.318 | 7.908±0.254 | 7.964±0.307 | 0.024 | [-0.076, 0.126] | 0.655 |
| POPT_left | 7.621±0.167 | 7.645±0.175 | 7.616±0.165 | 7.654±0.170 | 0.026 | [-0.075, 0.126] | 0.666 |
| POPT_right | 7.618±0.152 | 7.572±0.214 | 7.613±0.154 | 7.582±0.203 | 0.029 | [-0.072, 0.129] | 0.679 |
| SCP_left | 8.003±0.229 | 8.091±0.248 | 8.007±0.225 | 8.085±0.249 | 0.024 | [-0.075, 0.126] | 0.653 |
| SCP_right | 8.297±0.265 | 8.268±0.250 | 8.293±0.253 | 8.275±0.248 | 0.028 | [-0.074, 0.128] | 0.675 |
| SLF_III_left | 7.091±0.220 | 7.037±0.192 | 7.083±0.216 | 7.052±0.189 | 0.029 | [-0.070, 0.131] | 0.680 |
| SLF_III_right | 7.323±0.231 | 7.263±0.222 | 7.319±0.233 | 7.270±0.214 | 0.030 | [-0.072, 0.129] | 0.684 |
| SLF_II_left | 7.090±0.198 | 7.084±0.212 | 7.081±0.192 | 7.101±0.213 | 0.027 | [-0.074, 0.127] | 0.670 |
| SLF_II_right | 7.203±0.206 | 7.158±0.223 | 7.196±0.204 | 7.171±0.212 | 0.029 | [-0.070, 0.131] | 0.681 |
| SLF_I_left | 7.286±0.240 | 7.244±0.236 | 7.270±0.234 | 7.274±0.223 | 0.028 | [-0.072, 0.129] | 0.677 |
| SLF_I_right | 7.381±0.224 | 7.319±0.242 | 7.372±0.218 | 7.336±0.235 | 0.030 | [-0.071, 0.130] | 0.684 |
| STR_left | 7.020±0.199 | 6.985±0.172 | 7.015±0.196 | 6.993±0.170 | 0.029 | [-0.071, 0.130] | 0.679 |
| STR_right | 7.023±0.177 | 6.933±0.194 | 7.016±0.173 | 6.945±0.190 | 0.031 | [-0.069, 0.133] | 0.691 |
| UF_left | 7.665±0.228 | 7.459±0.184 | 7.649±0.220 | 7.489±0.200 | 0.034 | [-0.068, 0.134] | 0.703 |
| UF_right | 7.874±0.212 | 7.773±0.153 | 7.865±0.206 | 7.790±0.158 | 0.030 | [-0.070, 0.131] | 0.686 |

[1] Units for raw and adjusted means, difference, and 95% HDI are $10^{-4}$ mm²/s.

Data are presented as mean ± SD. Raw values are unadjusted sex-wise means. Adjusted values are sex-wise mean residual values after adjusting for age, education, APOE e4 carriership, and family history of Alzheimer's disease. Bayesian model outputs include the mean difference and 95% highest density interval estimates as well as P+. P+ indicates the proportion of the Bayesian samples (N = 80000) where the difference between women and men was > 0. P+ values approaching 1 indicate stronger evidence of female > male while P+ values approaching 0 indicate stronger evidence of male > female. The hierarchical model includes all tracts in a singular model and leverages partial pooling of the data, therefore the mean difference will not always be exactly equal to the difference in covariate adjusted means.

**Supplemental table 16: Fiber density differences between post-menopausal women and age-matched men**

| | Raw means[1] | | Covariate adjusted means[1] | | Hierarchical Bayesian model[1] | | |
|---|---|---|---|---|---|---|---|
| Tract | Females | Males | Females | Males | Mean Difference | 95% HDI | P+ |
| AF_left | 0.369±0.014 | 0.356±0.015 | 0.368±0.015 | 0.358±0.014 | 0.010 | [0.001, 0.018] | 0.968 |
| AF_right | 0.394±0.016 | 0.383±0.016 | 0.393±0.015 | 0.384±0.015 | 0.008 | [0.000, 0.017] | 0.945 |
| ATR_left | 0.366±0.019 | 0.357±0.022 | 0.364±0.017 | 0.360±0.018 | 0.010 | [0.002, 0.019] | 0.970 |
| ATR_right | 0.363±0.017 | 0.360±0.020 | 0.362±0.017 | 0.362±0.017 | 0.007 | [-0.002, 0.016] | 0.894 |
| CA | 0.439±0.027 | 0.429±0.029 | 0.438±0.026 | 0.429±0.029 | 0.008 | [0.000, 0.017] | 0.942 |
| CC_1 | 0.449±0.018 | 0.434±0.019 | 0.447±0.018 | 0.437±0.014 | 0.012 | [0.003, 0.021] | 0.987 |
| CC_2 | 0.382±0.015 | 0.370±0.015 | 0.381±0.014 | 0.372±0.011 | 0.011 | [0.003, 0.020] | 0.980 |
| CC_3 | 0.402±0.025 | 0.383±0.025 | 0.401±0.024 | 0.385±0.021 | 0.015 | [0.006, 0.023] | 0.996 |
| CC_4 | 0.457±0.021 | 0.438±0.019 | 0.456±0.020 | 0.440±0.016 | 0.013 | [0.004, 0.022] | 0.992 |
| CC_5 | 0.430±0.019 | 0.411±0.018 | 0.429±0.018 | 0.413±0.015 | 0.013 | [0.004, 0.021] | 0.990 |
| CC_6 | 0.408±0.015 | 0.392±0.014 | 0.407±0.014 | 0.393±0.012 | 0.011 | [0.003, 0.020] | 0.983 |
| CC_7 | 0.388±0.015 | 0.369±0.013 | 0.387±0.014 | 0.370±0.012 | 0.013 | [0.004, 0.022] | 0.991 |
| CG_left | 0.353±0.014 | 0.342±0.014 | 0.352±0.014 | 0.344±0.012 | 0.008 | [0.000, 0.017] | 0.937 |
| CG_right | 0.364±0.017 | 0.351±0.013 | 0.363±0.016 | 0.353±0.011 | 0.009 | [0.000, 0.017] | 0.948 |
| CST_left | 0.553±0.022 | 0.555±0.021 | 0.553±0.020 | 0.556±0.020 | 0.001 | [-0.008, 0.010] | 0.577 |
| CST_right | 0.520±0.018 | 0.516±0.019 | 0.520±0.018 | 0.517±0.017 | 0.003 | [-0.006, 0.012] | 0.708 |
| FPT_left | 0.476±0.020 | 0.474±0.017 | 0.475±0.019 | 0.476±0.014 | 0.004 | [-0.004, 0.013] | 0.793 |
| FPT_right | 0.468±0.019 | 0.464±0.019 | 0.467±0.019 | 0.466±0.016 | 0.005 | [-0.004, 0.013] | 0.809 |
| FX_left | 0.513±0.054 | 0.462±0.067 | 0.507±0.051 | 0.470±0.061 | 0.034 | [0.024, 0.043] | 1.000 |
| FX_right | 0.517±0.060 | 0.470±0.064 | 0.510±0.054 | 0.480±0.050 | 0.036 | [0.026, 0.045] | 1.000 |
| ICP_left | 0.417±0.019 | 0.411±0.019 | 0.416±0.018 | 0.412±0.018 | 0.006 | [-0.003, 0.014] | 0.855 |
| ICP_right | 0.382±0.018 | 0.374±0.014 | 0.381±0.017 | 0.375±0.013 | 0.008 | [0.000, 0.017] | 0.941 |
| IFO_left | 0.387±0.015 | 0.373±0.018 | 0.385±0.015 | 0.375±0.014 | 0.011 | [0.003, 0.020] | 0.983 |
| IFO_right | 0.385±0.015 | 0.371±0.013 | 0.384±0.014 | 0.372±0.012 | 0.011 | [0.002, 0.019] | 0.976 |
| ILF_left | 0.388±0.017 | 0.375±0.019 | 0.387±0.017 | 0.376±0.017 | 0.010 | [0.001, 0.018] | 0.969 |
| ILF_right | 0.393±0.016 | 0.385±0.021 | 0.392±0.015 | 0.386±0.021 | 0.007 | [-0.002, 0.015] | 0.909 |
| MCP | 0.455±0.019 | 0.444±0.017 | 0.454±0.018 | 0.446±0.016 | 0.007 | [-0.001, 0.016] | 0.917 |
| MLF_left | 0.376±0.011 | 0.362±0.014 | 0.375±0.011 | 0.363±0.012 | 0.010 | [0.001, 0.019] | 0.969 |
| MLF_right | 0.382±0.016 | 0.365±0.014 | 0.381±0.015 | 0.367±0.013 | 0.012 | [0.003, 0.020] | 0.985 |
| OR_left | 0.397±0.015 | 0.384±0.014 | 0.396±0.016 | 0.385±0.012 | 0.010 | [0.001, 0.018] | 0.963 |
| OR_right | 0.408±0.016 | 0.390±0.015 | 0.407±0.016 | 0.391±0.013 | 0.012 | [0.003, 0.020] | 0.985 |
| POPT_left | 0.464±0.013 | 0.454±0.014 | 0.463±0.012 | 0.455±0.011 | 0.007 | [-0.002, 0.015] | 0.897 |
| POPT_right | 0.463±0.016 | 0.455±0.015 | 0.462±0.016 | 0.456±0.013 | 0.006 | [-0.003, 0.014] | 0.856 |
| SCP_left | 0.469±0.018 | 0.474±0.014 | 0.469±0.017 | 0.474±0.014 | 0.000 | [-0.009, 0.008] | 0.467 |
| SCP_right | 0.444±0.017 | 0.446±0.015 | 0.443±0.016 | 0.447±0.015 | 0.001 | [-0.008, 0.010] | 0.578 |
| SLF_III_left | 0.370±0.018 | 0.358±0.020 | 0.368±0.018 | 0.361±0.016 | 0.011 | [0.002, 0.019] | 0.974 |
| SLF_III_right | 0.373±0.021 | 0.366±0.019 | 0.372±0.019 | 0.368±0.017 | 0.007 | [-0.002, 0.016] | 0.906 |
| SLF_II_left | 0.335±0.019 | 0.326±0.019 | 0.334±0.018 | 0.328±0.015 | 0.009 | [0.000, 0.018] | 0.951 |
| SLF_II_right | 0.333±0.016 | 0.322±0.015 | 0.332±0.016 | 0.323±0.013 | 0.009 | [0.001, 0.018] | 0.956 |
| SLF_I_left | 0.310±0.015 | 0.304±0.012 | 0.309±0.014 | 0.306±0.009 | 0.007 | [-0.002, 0.016] | 0.906 |
| SLF_I_right | 0.331±0.015 | 0.322±0.012 | 0.330±0.014 | 0.324±0.010 | 0.008 | [-0.001, 0.016] | 0.926 |
| STR_left | 0.496±0.023 | 0.491±0.020 | 0.496±0.022 | 0.491±0.020 | 0.004 | [-0.005, 0.013] | 0.763 |
| STR_right | 0.468±0.021 | 0.464±0.020 | 0.468±0.021 | 0.464±0.019 | 0.003 | [-0.005, 0.012] | 0.732 |
| UF_left | 0.421±0.018 | 0.412±0.023 | 0.420±0.017 | 0.414±0.021 | 0.008 | [-0.001, 0.016] | 0.929 |
| UF_right | 0.411±0.020 | 0.402±0.020 | 0.410±0.020 | 0.404±0.018 | 0.009 | [0.000, 0.017] | 0.945 |

[1] Units for raw and adjusted means, difference, and 95% HDI are arbitrary.

Data are presented as mean ± SD. Raw values are unadjusted sex-wise means. Adjusted values are sex-wise mean residual values after adjusting for age, education, APOE e4 carriership, and family history of Alzheimer's disease. Bayesian model outputs include the mean difference and 95% highest density interval estimates as well as P+. P+ indicates the proportion of the Bayesian samples (N = 80000) where the difference between women and men was > 0. P+ values approaching 1 indicate stronger evidence of female > male while P+ values approaching 0 indicate stronger evidence of male > female. The hierarchical model includes all tracts in a singular model and leverages partial pooling of the data, therefore the mean difference will not always be exactly equal to the difference in covariate adjusted means.

**Supplemental table 17: Fiber cross-section differences between post-menopausal women and age-matched men**

| Tract | Raw means[1] | | Covariate adjusted means[1] | | Hierarchical Bayesian model[1] | | |
|---|---|---|---|---|---|---|---|
| | Females | Males | Females | Males | Mean Difference | 95% HDI | P+ |
| AF_left | -0.005±0.057 | 0.065±0.075 | 0.027±0.036 | 0.018±0.030 | 0.014 | [-0.003, 0.031] | 0.900 |
| AF_right | -0.005±0.053 | 0.070±0.075 | 0.027±0.031 | 0.021±0.034 | 0.013 | [-0.004, 0.031] | 0.889 |
| ATR_left | -0.034±0.064 | 0.054±0.088 | 0.003±0.051 | 0.000±0.030 | 0.010 | [-0.008, 0.028] | 0.824 |
| ATR_right | -0.032±0.065 | 0.060±0.078 | 0.003±0.047 | 0.008±0.036 | 0.010 | [-0.007, 0.029] | 0.828 |
| CA | -0.027±0.058 | 0.042±0.080 | 0.003±0.042 | -0.003±0.057 | 0.012 | [-0.006, 0.030] | 0.869 |
| CC_1 | -0.044±0.055 | 0.030±0.077 | -0.017±0.039 | -0.010±0.045 | 0.009 | [-0.010, 0.028] | 0.783 |
| CC_2 | -0.015±0.060 | 0.063±0.085 | 0.018±0.042 | 0.014±0.035 | 0.012 | [-0.006, 0.029] | 0.857 |
| CC_3 | 0.009±0.079 | 0.063±0.106 | 0.044±0.053 | 0.010±0.062 | 0.018 | [-0.002, 0.036] | 0.938 |
| CC_4 | -0.005±0.060 | 0.051±0.097 | 0.022±0.051 | 0.011±0.051 | 0.014 | [-0.003, 0.032] | 0.904 |
| CC_5 | -0.015±0.068 | 0.039±0.071 | 0.011±0.049 | 0.000±0.036 | 0.013 | [-0.004, 0.031] | 0.892 |
| CC_6 | -0.017±0.054 | 0.037±0.067 | 0.010±0.032 | -0.004±0.041 | 0.014 | [-0.003, 0.032] | 0.909 |
| CC_7 | 0.037±0.073 | 0.094±0.094 | 0.065±0.058 | 0.052±0.062 | 0.015 | [-0.003, 0.033] | 0.908 |
| CG_left | -0.035±0.059 | 0.033±0.065 | -0.007±0.040 | -0.009±0.036 | 0.011 | [-0.007, 0.029] | 0.836 |
| CG_right | -0.040±0.052 | 0.044±0.058 | -0.011±0.032 | 0.000±0.037 | 0.010 | [-0.008, 0.028] | 0.806 |
| CST_left | -0.008±0.053 | 0.026±0.067 | 0.010±0.047 | -0.001±0.040 | 0.014 | [-0.003, 0.032] | 0.908 |
| CST_right | -0.008±0.048 | 0.029±0.075 | 0.010±0.043 | 0.001±0.047 | 0.014 | [-0.003, 0.031] | 0.900 |
| FPT_left | -0.013±0.049 | 0.032±0.075 | 0.009±0.038 | -0.001±0.039 | 0.013 | [-0.004, 0.030] | 0.883 |
| FPT_right | -0.012±0.048 | 0.038±0.080 | 0.012±0.038 | 0.002±0.048 | 0.013 | [-0.004, 0.031] | 0.889 |
| FX_left | -0.080±0.114 | 0.055±0.139 | -0.019±0.081 | -0.036±0.086 | 0.012 | [-0.007, 0.032] | 0.850 |
| FX_right | -0.077±0.118 | 0.059±0.138 | -0.022±0.090 | -0.024±0.092 | 0.012 | [-0.008, 0.031] | 0.840 |
| ICP_left | -0.025±0.060 | 0.013±0.062 | -0.005±0.040 | -0.016±0.056 | 0.014 | [-0.003, 0.032] | 0.901 |
| ICP_right | -0.020±0.066 | 0.019±0.054 | -0.002±0.041 | -0.008±0.044 | 0.015 | [-0.004, 0.033] | 0.909 |
| IFO_left | -0.001±0.058 | 0.076±0.066 | 0.029±0.040 | 0.030±0.030 | 0.011 | [-0.006, 0.029] | 0.852 |
| IFO_right | 0.007±0.057 | 0.075±0.067 | 0.035±0.040 | 0.033±0.037 | 0.012 | [-0.005, 0.030] | 0.875 |
| ILF_left | 0.014±0.057 | 0.098±0.063 | 0.047±0.035 | 0.048±0.039 | 0.012 | [-0.006, 0.030] | 0.861 |
| ILF_right | 0.010±0.063 | 0.101±0.074 | 0.048±0.038 | 0.045±0.038 | 0.012 | [-0.006, 0.030] | 0.866 |
| MCP | -0.009±0.060 | 0.044±0.061 | 0.012±0.040 | 0.013±0.049 | 0.013 | [-0.005, 0.031] | 0.885 |
| MLF_left | -0.012±0.053 | 0.058±0.063 | 0.019±0.031 | 0.012±0.035 | 0.013 | [-0.004, 0.031] | 0.891 |
| MLF_right | -0.019±0.054 | 0.048±0.060 | 0.010±0.034 | 0.004±0.035 | 0.013 | [-0.004, 0.031] | 0.890 |
| OR_left | 0.014±0.067 | 0.084±0.077 | 0.044±0.045 | 0.038±0.047 | 0.013 | [-0.004, 0.031] | 0.886 |
| OR_right | 0.012±0.068 | 0.063±0.086 | 0.038±0.056 | 0.023±0.057 | 0.015 | [-0.003, 0.033] | 0.912 |
| POPT_left | -0.013±0.049 | 0.022±0.053 | 0.007±0.034 | -0.007±0.035 | 0.015 | [-0.003, 0.032] | 0.910 |
| POPT_right | -0.016±0.046 | 0.013±0.052 | 0.002±0.033 | -0.014±0.036 | 0.015 | [-0.003, 0.033] | 0.914 |
| SCP_left | -0.030±0.036 | -0.006±0.059 | -0.015±0.027 | -0.027±0.047 | 0.014 | [-0.004, 0.032] | 0.891 |
| SCP_right | -0.030±0.037 | -0.006±0.056 | -0.016±0.030 | -0.026±0.043 | 0.014 | [-0.004, 0.032] | 0.894 |
| SLF_III_left | -0.003±0.060 | 0.070±0.074 | 0.027±0.044 | 0.025±0.043 | 0.013 | [-0.005, 0.030] | 0.887 |
| SLF_III_right | -0.006±0.052 | 0.069±0.075 | 0.023±0.034 | 0.026±0.055 | 0.013 | [-0.004, 0.031] | 0.884 |
| SLF_II_left | 0.004±0.060 | 0.078±0.078 | 0.039±0.039 | 0.027±0.031 | 0.014 | [-0.003, 0.032] | 0.906 |
| SLF_II_right | -0.004±0.063 | 0.077±0.072 | 0.030±0.044 | 0.025±0.031 | 0.013 | [-0.004, 0.031] | 0.882 |
| SLF_I_left | 0.016±0.055 | 0.065±0.072 | 0.042±0.039 | 0.027±0.040 | 0.015 | [-0.003, 0.033] | 0.919 |
| SLF_I_right | 0.005±0.056 | 0.057±0.063 | 0.031±0.034 | 0.018±0.041 | 0.014 | [-0.003, 0.032] | 0.911 |
| STR_left | -0.029±0.066 | 0.020±0.072 | -0.007±0.054 | -0.013±0.052 | 0.013 | [-0.005, 0.030] | 0.881 |
| STR_right | -0.018±0.054 | 0.024±0.075 | 0.002±0.046 | -0.005±0.049 | 0.013 | [-0.004, 0.031] | 0.887 |
| UF_left | -0.025±0.051 | 0.052±0.059 | 0.005±0.035 | 0.007±0.026 | 0.010 | [-0.007, 0.028] | 0.831 |
| UF_right | -0.013±0.047 | 0.068±0.061 | 0.016±0.028 | 0.024±0.040 | 0.010 | [-0.007, 0.028] | 0.830 |

[1] Units for raw and adjusted means, difference, and 95% HDI are arbitrary.

Data are presented as mean ± SD. Raw values are unadjusted sex-wise means. Adjusted values are sex-wise mean residual values after adjusting for age, education, intracranial volume, APOE e4 carriership, and family history of Alzheimer's disease. Bayesian model outputs include the mean difference and 95% highest density interval estimates as well as P+. P+ values approaching 1 indicate stronger evidence of female > male while P+ values approaching 0 indicate stronger evidence of male > female. The hierarchical model includes all tracts in a singular model and leverages partial pooling of the data, therefore the mean difference will not always be exactly equal to the difference in covariate adjusted means.

### Supplemental table 18: FDC differences between post-menopausal women and age-matched men

| | Raw means[1] | | Covariate adjusted means[1] | | Hierarchical Bayesian model[1] | | |
|---|---|---|---|---|---|---|---|
| Tract | Females | Males | Females | Males | Mean Difference | 95% HDI | P+ |
| AF_left | 0.372±0.030 | 0.382±0.033 | 0.380±0.026 | 0.370±0.022 | 0.011 | [-0.004, 0.026] | 0.888 |
| AF_right | 0.395±0.028 | 0.412±0.034 | 0.404±0.024 | 0.398±0.024 | 0.010 | [-0.005, 0.025] | 0.850 |
| ATR_left | 0.355±0.028 | 0.377±0.036 | 0.364±0.028 | 0.364±0.022 | 0.008 | [-0.008, 0.023] | 0.788 |
| ATR_right | 0.354±0.027 | 0.381±0.031 | 0.362±0.026 | 0.368±0.023 | 0.007 | [-0.008, 0.022] | 0.772 |
| CA | 0.427±0.035 | 0.447±0.054 | 0.439±0.030 | 0.429±0.046 | 0.010 | [-0.005, 0.026] | 0.866 |
| CC_1 | 0.433±0.025 | 0.445±0.046 | 0.438±0.025 | 0.439±0.030 | 0.008 | [-0.008, 0.023] | 0.788 |
| CC_2 | 0.380±0.027 | 0.394±0.037 | 0.387±0.025 | 0.383±0.024 | 0.010 | [-0.005, 0.025] | 0.860 |
| CC_3 | 0.409±0.047 | 0.407±0.042 | 0.417±0.040 | 0.395±0.036 | 0.014 | [-0.002, 0.030] | 0.928 |
| CC_4 | 0.459±0.039 | 0.463±0.049 | 0.465±0.035 | 0.454±0.037 | 0.013 | [-0.003, 0.028] | 0.909 |
| CC_5 | 0.426±0.038 | 0.428±0.033 | 0.431±0.034 | 0.421±0.023 | 0.012 | [-0.003, 0.027] | 0.901 |
| CC_6 | 0.404±0.030 | 0.406±0.029 | 0.410±0.025 | 0.397±0.022 | 0.013 | [-0.002, 0.028] | 0.914 |
| CC_7 | 0.402±0.038 | 0.402±0.038 | 0.408±0.034 | 0.394±0.028 | 0.015 | [-0.001, 0.030] | 0.934 |
| CG_left | 0.343±0.024 | 0.354±0.026 | 0.349±0.022 | 0.346±0.020 | 0.010 | [-0.005, 0.025] | 0.863 |
| CG_right | 0.352±0.023 | 0.368±0.024 | 0.358±0.022 | 0.359±0.019 | 0.009 | [-0.006, 0.024] | 0.831 |
| CST_left | 0.551±0.044 | 0.569±0.043 | 0.560±0.040 | 0.556±0.028 | 0.009 | [-0.006, 0.025] | 0.828 |
| CST_right | 0.518±0.035 | 0.530±0.043 | 0.524±0.034 | 0.521±0.033 | 0.009 | [-0.006, 0.024] | 0.830 |
| FPT_left | 0.472±0.035 | 0.488±0.043 | 0.480±0.033 | 0.476±0.028 | 0.008 | [-0.007, 0.023] | 0.814 |
| FPT_right | 0.464±0.032 | 0.480±0.046 | 0.472±0.032 | 0.469±0.035 | 0.009 | [-0.006, 0.024] | 0.820 |
| FX_left | 0.467±0.036 | 0.475±0.053 | 0.470±0.034 | 0.469±0.051 | 0.009 | [-0.006, 0.025] | 0.845 |
| FX_right | 0.470±0.042 | 0.485±0.052 | 0.477±0.040 | 0.475±0.041 | 0.009 | [-0.006, 0.025] | 0.841 |
| ICP_left | 0.409±0.030 | 0.420±0.030 | 0.414±0.024 | 0.412±0.028 | 0.009 | [-0.007, 0.023] | 0.819 |
| ICP_right | 0.377±0.033 | 0.382±0.022 | 0.381±0.025 | 0.376±0.018 | 0.009 | [-0.006, 0.024] | 0.825 |
| IFO_left | 0.384±0.029 | 0.398±0.027 | 0.391±0.026 | 0.388±0.018 | 0.010 | [-0.005, 0.025] | 0.858 |
| IFO_right | 0.384±0.027 | 0.394±0.025 | 0.390±0.025 | 0.385±0.018 | 0.011 | [-0.004, 0.026] | 0.887 |
| ILF_left | 0.391±0.030 | 0.411±0.032 | 0.400±0.025 | 0.397±0.025 | 0.010 | [-0.005, 0.025] | 0.858 |
| ILF_right | 0.393±0.034 | 0.423±0.039 | 0.408±0.024 | 0.401±0.030 | 0.009 | [-0.006, 0.025] | 0.836 |
| MCP | 0.456±0.039 | 0.470±0.037 | 0.463±0.032 | 0.459±0.030 | 0.007 | [-0.009, 0.022] | 0.767 |
| MLF_left | 0.373±0.022 | 0.381±0.023 | 0.380±0.018 | 0.371±0.016 | 0.011 | [-0.003, 0.027] | 0.889 |
| MLF_right | 0.376±0.029 | 0.382±0.024 | 0.382±0.024 | 0.373±0.017 | 0.012 | [-0.003, 0.027] | 0.895 |
| OR_left | 0.399±0.032 | 0.411±0.029 | 0.406±0.028 | 0.400±0.021 | 0.011 | [-0.004, 0.026] | 0.888 |
| OR_right | 0.408±0.034 | 0.411±0.034 | 0.414±0.031 | 0.401±0.025 | 0.013 | [-0.002, 0.029] | 0.918 |
| POPT_left | 0.459±0.027 | 0.463±0.027 | 0.464±0.023 | 0.455±0.021 | 0.011 | [-0.004, 0.026] | 0.881 |
| POPT_right | 0.456±0.031 | 0.460±0.027 | 0.461±0.028 | 0.453±0.020 | 0.011 | [-0.004, 0.026] | 0.883 |
| SCP_left | 0.457±0.030 | 0.473±0.029 | 0.464±0.027 | 0.463±0.024 | 0.008 | [-0.007, 0.023] | 0.801 |
| SCP_right | 0.433±0.029 | 0.445±0.027 | 0.439±0.027 | 0.436±0.021 | 0.008 | [-0.007, 0.023] | 0.809 |
| SLF_III_left | 0.373±0.031 | 0.386±0.034 | 0.380±0.029 | 0.375±0.026 | 0.009 | [-0.006, 0.024] | 0.843 |
| SLF_III_right | 0.374±0.030 | 0.395±0.034 | 0.382±0.027 | 0.383±0.031 | 0.008 | [-0.007, 0.023] | 0.796 |
| SLF_II_left | 0.341±0.030 | 0.356±0.029 | 0.349±0.026 | 0.344±0.022 | 0.010 | [-0.005, 0.025] | 0.855 |
| SLF_II_right | 0.336±0.030 | 0.351±0.027 | 0.344±0.027 | 0.340±0.022 | 0.010 | [-0.005, 0.025] | 0.856 |
| SLF_I_left | 0.319±0.023 | 0.328±0.029 | 0.324±0.021 | 0.320±0.020 | 0.011 | [-0.004, 0.026] | 0.877 |
| SLF_I_right | 0.337±0.026 | 0.345±0.026 | 0.343±0.022 | 0.336±0.020 | 0.011 | [-0.004, 0.026] | 0.885 |
| STR_left | 0.486±0.047 | 0.500±0.039 | 0.495±0.042 | 0.487±0.030 | 0.010 | [-0.005, 0.025] | 0.866 |
| STR_right | 0.462±0.039 | 0.474±0.043 | 0.468±0.037 | 0.464±0.034 | 0.010 | [-0.005, 0.025] | 0.857 |
| UF_left | 0.411±0.026 | 0.432±0.032 | 0.419±0.024 | 0.420±0.026 | 0.008 | [-0.007, 0.023] | 0.799 |
| UF_right | 0.406±0.026 | 0.429±0.033 | 0.414±0.024 | 0.416±0.026 | 0.008 | [-0.008, 0.023] | 0.794 |

[1] Units for raw and adjusted means, difference, and 95% HDI are arbitrary.

Data are presented as mean ± SD. Raw values are unadjusted sex-wise means. Adjusted values are sex-wise mean residual values after adjusting for age, education, intracranial volume, APOE e4 carriership, and family history of Alzheimer's disease. Bayesian model outputs include the mean difference and 95% highest density interval estimates as well as P+. P+ values approaching 1 indicate stronger evidence of female > male while P+ values approaching 0 indicate stronger evidence of male > female. The hierarchical model includes all tracts in a singular model and leverages partial pooling of the data, therefore the mean difference will not always be exactly equal to the difference in covariate adjusted means.

### Supplemental table 19: Fractional anisotropy differences between post-menopausal women and age-matched men

| | Raw means[1] | | Covariate adjusted means[1] | | Hierarchical Bayesian model[1] | | |
|---|---|---|---|---|---|---|---|
| Tract | Females | Males | Females | Males | Mean Difference | 95% HDI | P+ |
| AF_left | 0.386±0.015 | 0.379±0.018 | 0.385±0.015 | 0.380±0.017 | 0.003 | [-0.006, 0.011] | 0.691 |
| AF_right | 0.385±0.013 | 0.382±0.017 | 0.384±0.013 | 0.383±0.015 | 0.000 | [-0.008, 0.009] | 0.533 |
| ATR_left | 0.360±0.015 | 0.365±0.013 | 0.359±0.014 | 0.366±0.013 | -0.003 | [-0.011, 0.006] | 0.316 |
| ATR_right | 0.342±0.013 | 0.344±0.013 | 0.341±0.012 | 0.345±0.013 | 0.000 | [-0.008, 0.009] | 0.521 |
| CA | 0.394±0.043 | 0.386±0.058 | 0.395±0.043 | 0.385±0.057 | 0.005 | [-0.003, 0.014] | 0.837 |
| CC_1 | 0.433±0.015 | 0.428±0.025 | 0.431±0.014 | 0.431±0.020 | 0.004 | [-0.005, 0.012] | 0.749 |
| CC_2 | 0.409±0.015 | 0.404±0.018 | 0.408±0.015 | 0.405±0.014 | 0.004 | [-0.004, 0.012] | 0.778 |
| CC_3 | 0.403±0.017 | 0.397±0.020 | 0.403±0.016 | 0.398±0.017 | 0.005 | [-0.004, 0.013] | 0.824 |
| CC_4 | 0.448±0.017 | 0.438±0.017 | 0.448±0.017 | 0.438±0.013 | 0.006 | [-0.003, 0.014] | 0.863 |
| CC_5 | 0.448±0.020 | 0.435±0.026 | 0.448±0.021 | 0.435±0.022 | 0.007 | [-0.002, 0.015] | 0.894 |
| CC_6 | 0.478±0.016 | 0.474±0.016 | 0.478±0.016 | 0.475±0.014 | 0.003 | [-0.006, 0.011] | 0.683 |
| CC_7 | 0.493±0.027 | 0.475±0.022 | 0.492±0.025 | 0.477±0.018 | 0.012 | [0.003, 0.021] | 0.984 |
| CG_left | 0.409±0.019 | 0.414±0.023 | 0.409±0.019 | 0.414±0.022 | -0.005 | [-0.013, 0.004] | 0.191 |
| CG_right | 0.399±0.017 | 0.400±0.024 | 0.398±0.017 | 0.400±0.023 | -0.003 | [-0.012, 0.005] | 0.262 |
| CST_left | 0.489±0.017 | 0.495±0.017 | 0.489±0.016 | 0.495±0.016 | -0.005 | [-0.013, 0.004] | 0.178 |
| CST_right | 0.483±0.016 | 0.492±0.014 | 0.484±0.015 | 0.492±0.013 | -0.007 | [-0.016, 0.001] | 0.081 |
| FPT_left | 0.456±0.015 | 0.465±0.015 | 0.456±0.015 | 0.466±0.014 | -0.006 | [-0.014, 0.003] | 0.127 |
| FPT_right | 0.447±0.015 | 0.457±0.016 | 0.447±0.015 | 0.457±0.015 | -0.007 | [-0.015, 0.002] | 0.098 |
| FX_left | 0.390±0.042 | 0.357±0.057 | 0.386±0.041 | 0.363±0.053 | 0.021 | [0.012, 0.030] | 1.000 |
| FX_right | 0.377±0.046 | 0.342±0.053 | 0.373±0.047 | 0.349±0.045 | 0.025 | [0.016, 0.034] | 1.000 |
| ICP_left | 0.411±0.015 | 0.417±0.017 | 0.411±0.015 | 0.417±0.016 | -0.005 | [-0.013, 0.003] | 0.172 |
| ICP_right | 0.417±0.018 | 0.422±0.019 | 0.416±0.017 | 0.423±0.017 | -0.004 | [-0.012, 0.005] | 0.255 |
| IFO_left | 0.418±0.015 | 0.410±0.016 | 0.417±0.015 | 0.411±0.014 | 0.005 | [-0.004, 0.013] | 0.821 |
| IFO_right | 0.425±0.016 | 0.418±0.017 | 0.424±0.016 | 0.420±0.015 | 0.004 | [-0.004, 0.012] | 0.788 |
| ILF_left | 0.409±0.016 | 0.401±0.021 | 0.408±0.016 | 0.402±0.018 | 0.005 | [-0.003, 0.014] | 0.849 |
| ILF_right | 0.409±0.016 | 0.408±0.013 | 0.409±0.015 | 0.409±0.012 | 0.001 | [-0.008, 0.009] | 0.561 |
| MCP | 0.480±0.015 | 0.482±0.016 | 0.480±0.015 | 0.482±0.015 | -0.004 | [-0.012, 0.005] | 0.228 |
| MLF_left | 0.384±0.014 | 0.377±0.016 | 0.383±0.014 | 0.378±0.014 | 0.004 | [-0.005, 0.012] | 0.752 |
| MLF_right | 0.376±0.015 | 0.372±0.014 | 0.375±0.015 | 0.372±0.013 | 0.003 | [-0.006, 0.011] | 0.690 |
| OR_left | 0.423±0.017 | 0.413±0.014 | 0.422±0.017 | 0.415±0.013 | 0.005 | [-0.003, 0.014] | 0.839 |
| OR_right | 0.431±0.019 | 0.425±0.018 | 0.430±0.020 | 0.426±0.015 | 0.004 | [-0.005, 0.012] | 0.767 |
| POPT_left | 0.452±0.013 | 0.455±0.012 | 0.452±0.012 | 0.455±0.011 | -0.003 | [-0.011, 0.005] | 0.278 |
| POPT_right | 0.458±0.014 | 0.464±0.013 | 0.458±0.013 | 0.464±0.012 | -0.005 | [-0.013, 0.004] | 0.171 |
| SCP_left | 0.433±0.013 | 0.445±0.010 | 0.433±0.013 | 0.444±0.010 | -0.009 | [-0.017, 0.000] | 0.050 |
| SCP_right | 0.413±0.015 | 0.428±0.015 | 0.413±0.015 | 0.428±0.014 | -0.010 | [-0.018, -0.001] | 0.032 |
| SLF_III_left | 0.391±0.016 | 0.387±0.016 | 0.390±0.016 | 0.388±0.015 | 0.003 | [-0.006, 0.011] | 0.686 |
| SLF_III_right | 0.380±0.015 | 0.380±0.017 | 0.380±0.014 | 0.381±0.015 | -0.001 | [-0.009, 0.008] | 0.446 |
| SLF_II_left | 0.351±0.017 | 0.345±0.018 | 0.350±0.017 | 0.346±0.017 | 0.003 | [-0.006, 0.011] | 0.700 |
| SLF_II_right | 0.357±0.015 | 0.354±0.014 | 0.357±0.016 | 0.354±0.013 | 0.001 | [-0.007, 0.010] | 0.592 |
| SLF_I_left | 0.374±0.018 | 0.376±0.016 | 0.374±0.016 | 0.376±0.014 | -0.001 | [-0.009, 0.008] | 0.446 |
| SLF_I_right | 0.375±0.016 | 0.374±0.015 | 0.375±0.016 | 0.374±0.014 | 0.000 | [-0.008, 0.009] | 0.523 |
| STR_left | 0.394±0.020 | 0.398±0.018 | 0.395±0.020 | 0.396±0.016 | -0.004 | [-0.012, 0.005] | 0.244 |
| STR_right | 0.393±0.018 | 0.405±0.020 | 0.394±0.017 | 0.403±0.020 | -0.008 | [-0.017, 0.000] | 0.057 |
| UF_left | 0.359±0.015 | 0.358±0.021 | 0.358±0.015 | 0.360±0.020 | 0.000 | [-0.009, 0.008] | 0.498 |
| UF_right | 0.349±0.014 | 0.348±0.019 | 0.348±0.014 | 0.350±0.017 | 0.000 | [-0.008, 0.009] | 0.538 |

[1] Units for raw and adjusted means, difference, and 95% HDI are arbitrary.

Data are presented as mean ± SD. Raw values are unadjusted sex-wise means. Adjusted values are sex-wise mean residual values after adjusting for age, education, APOE e4 carriership, and family history of Alzheimer's disease. Bayesian model outputs include the mean difference and 95% highest density interval estimates as well as P+. P+ indicates the proportion of the Bayesian samples (N = 80000) where the difference between women and men was > 0. P+ values approaching 1 indicate stronger evidence of female > male while P+ values approaching 0 indicate stronger evidence of male > female. The hierarchical model includes all tracts in a singular model and leverages partial pooling of the data, therefore the mean difference will not always be exactly equal to the difference in covariate adjusted means.

**Supplemental table 20: Mean diffusivity differences between post-menopausal women and age-matched men**

| | Raw means[1] | | Covariate adjusted means[1] | | Hierarchical Bayesian model[1] | | |
|---|---|---|---|---|---|---|---|
| Tract | Females | Males | Females | Males | Mean Difference | 95% HDI | P+ |
| AF_left | 7.141±0.193 | 7.204±0.203 | 7.145±0.198 | 7.199±0.189 | -0.024 | [-0.157, 0.110] | 0.386 |
| AF_right | 7.218±0.174 | 7.184±0.245 | 7.217±0.181 | 7.186±0.211 | -0.012 | [-0.148, 0.121] | 0.440 |
| ATR_left | 7.568±0.202 | 7.675±0.217 | 7.575±0.208 | 7.664±0.190 | -0.062 | [-0.195, 0.070] | 0.222 |
| ATR_right | 7.840±0.183 | 7.944±0.210 | 7.847±0.187 | 7.934±0.161 | -0.082 | [-0.212, 0.054] | 0.161 |
| CA | 8.128±0.398 | 8.253±0.395 | 8.146±0.380 | 8.227±0.380 | -0.086 | [-0.225, 0.049] | 0.157 |
| CC_1 | 8.123±0.320 | 8.074±0.334 | 8.127±0.313 | 8.068±0.249 | -0.074 | [-0.210, 0.062] | 0.188 |
| CC_2 | 8.397±0.244 | 8.554±0.265 | 8.402±0.228 | 8.547±0.225 | -0.134 | [-0.267, 0.001] | 0.056 |
| CC_3 | 8.670±0.353 | 8.904±0.329 | 8.680±0.325 | 8.889±0.309 | -0.168 | [-0.300, -0.032] | 0.023 |
| CC_4 | 8.365±0.345 | 8.639±0.318 | 8.378±0.328 | 8.620±0.276 | -0.155 | [-0.291, -0.021] | 0.033 |
| CC_5 | 8.834±0.398 | 9.239±0.638 | 8.850±0.394 | 9.215±0.578 | -0.212 | [-0.352, -0.071] | 0.007 |
| CC_6 | 7.912±0.242 | 8.072±0.295 | 7.923±0.237 | 8.054±0.262 | -0.098 | [-0.231, 0.034] | 0.115 |
| CC_7 | 9.242±0.776 | 9.637±0.724 | 9.277±0.662 | 9.584±0.679 | -0.250 | [-0.398, -0.106] | 0.003 |
| CG_left | 7.298±0.181 | 7.253±0.252 | 7.294±0.191 | 7.259±0.218 | -0.013 | [-0.145, 0.122] | 0.434 |
| CG_right | 7.272±0.181 | 7.239±0.215 | 7.267±0.188 | 7.246±0.183 | -0.014 | [-0.147, 0.119] | 0.431 |
| CST_left | 6.967±0.129 | 6.987±0.120 | 6.970±0.126 | 6.983±0.116 | -0.001 | [-0.136, 0.133] | 0.493 |
| CST_right | 7.107±0.131 | 7.061±0.139 | 7.106±0.134 | 7.064±0.130 | 0.001 | [-0.134, 0.135] | 0.503 |
| FPT_left | 7.140±0.123 | 7.131±0.169 | 7.137±0.128 | 7.135±0.157 | -0.009 | [-0.146, 0.121] | 0.457 |
| FPT_right | 7.354±0.130 | 7.278±0.169 | 7.348±0.137 | 7.287±0.151 | -0.010 | [-0.145, 0.123] | 0.445 |
| FX_left | 16.112±1.634 | 17.298±2.252 | 16.271±1.532 | 17.060±2.148 | -0.807 | [-0.998, -0.614] | 0.000 |
| FX_right | 16.180±1.838 | 17.346±2.033 | 16.380±1.692 | 17.047±1.727 | -0.820 | [-1.011, -0.628] | 0.000 |
| ICP_left | 7.141±0.166 | 7.100±0.161 | 7.139±0.149 | 7.104±0.168 | 0.001 | [-0.134, 0.135] | 0.505 |
| ICP_right | 6.873±0.162 | 6.819±0.161 | 6.876±0.147 | 6.814±0.167 | 0.023 | [-0.112, 0.157] | 0.609 |
| IFO_left | 7.819±0.241 | 7.988±0.265 | 7.831±0.242 | 7.971±0.212 | -0.094 | [-0.227, 0.039] | 0.127 |
| IFO_right | 7.905±0.322 | 8.013±0.394 | 7.920±0.309 | 7.990±0.315 | -0.091 | [-0.226, 0.044] | 0.136 |
| ILF_left | 7.824±0.283 | 8.006±0.299 | 7.841±0.282 | 7.980±0.216 | -0.096 | [-0.229, 0.038] | 0.125 |
| ILF_right | 7.770±0.349 | 7.855±0.382 | 7.790±0.337 | 7.826±0.294 | -0.074 | [-0.207, 0.062] | 0.185 |
| MCP | 6.857±0.133 | 6.768±0.156 | 6.855±0.120 | 6.771±0.162 | 0.030 | [-0.106, 0.166] | 0.635 |
| MLF_left | 7.576±0.190 | 7.689±0.256 | 7.584±0.194 | 7.677±0.229 | -0.063 | [-0.197, 0.068] | 0.224 |
| MLF_right | 7.409±0.194 | 7.437±0.288 | 7.415±0.196 | 7.429±0.259 | -0.037 | [-0.170, 0.095] | 0.323 |
| OR_left | 8.125±0.340 | 8.531±0.520 | 8.145±0.340 | 8.501±0.484 | -0.163 | [-0.303, -0.025] | 0.026 |
| OR_right | 7.968±0.443 | 8.173±0.521 | 7.995±0.404 | 8.134±0.402 | -0.119 | [-0.259, 0.019] | 0.084 |
| POPT_left | 7.667±0.172 | 7.747±0.168 | 7.673±0.172 | 7.737±0.163 | -0.060 | [-0.192, 0.075] | 0.234 |
| POPT_right | 7.629±0.139 | 7.674±0.211 | 7.636±0.141 | 7.663±0.201 | -0.050 | [-0.182, 0.084] | 0.269 |
| SCP_left | 8.017±0.204 | 8.051±0.191 | 8.014±0.186 | 8.055±0.201 | -0.072 | [-0.208, 0.063] | 0.194 |
| SCP_right | 8.270±0.231 | 8.268±0.197 | 8.277±0.232 | 8.257±0.186 | -0.083 | [-0.216, 0.052] | 0.159 |
| SLF_III_left | 7.116±0.217 | 7.176±0.203 | 7.122±0.220 | 7.168±0.187 | -0.022 | [-0.156, 0.112] | 0.391 |
| SLF_III_right | 7.369±0.207 | 7.356±0.251 | 7.369±0.217 | 7.356±0.213 | -0.024 | [-0.158, 0.109] | 0.383 |
| SLF_II_left | 7.104±0.166 | 7.155±0.240 | 7.108±0.170 | 7.149±0.228 | -0.018 | [-0.153, 0.112] | 0.412 |
| SLF_II_right | 7.213±0.172 | 7.234±0.234 | 7.213±0.171 | 7.233±0.211 | -0.023 | [-0.155, 0.112] | 0.390 |
| SLF_I_left | 7.345±0.187 | 7.336±0.257 | 7.352±0.189 | 7.325±0.243 | -0.022 | [-0.156, 0.110] | 0.391 |
| SLF_I_right | 7.412±0.215 | 7.421±0.263 | 7.417±0.203 | 7.414±0.242 | -0.033 | [-0.166, 0.101] | 0.343 |
| STR_left | 7.053±0.200 | 7.095±0.179 | 7.051±0.200 | 7.097±0.154 | -0.017 | [-0.149, 0.117] | 0.422 |
| STR_right | 7.047±0.162 | 7.008±0.198 | 7.044±0.165 | 7.012±0.159 | 0.001 | [-0.131, 0.135] | 0.502 |
| UF_left | 7.686±0.195 | 7.624±0.200 | 7.688±0.192 | 7.620±0.187 | -0.033 | [-0.167, 0.101] | 0.341 |
| UF_right | 7.903±0.203 | 7.895±0.198 | 7.908±0.207 | 7.888±0.178 | -0.056 | [-0.191, 0.078] | 0.244 |

[1] Units for raw and adjusted means, difference, and 95% HDI are $10^{-4}$ mm²/s.

Data are presented as mean ± SD. Raw values are unadjusted sex-wise means. Adjusted values are sex-wise mean residual values after adjusting for age, education, APOE e4 carriership, and family history of Alzheimer's disease. Bayesian model outputs include the mean difference and 95% highest density interval estimates as well as P+. P+ indicates the proportion of the Bayesian samples (N = 80000) where the difference between women and men was > 0. P+ values approaching 1 indicate stronger evidence of female > male while P+ values approaching 0 indicate stronger evidence of male > female. The hierarchical model includes all tracts in a singular model and leverages partial pooling of the data, therefore the mean difference will not always be exactly equal to the difference in covariate adjusted means.
